# A regulatory module driving the recurrent evolution of irreducible molecular complexes

**DOI:** 10.1101/2024.09.16.613340

**Authors:** Polina Tikanova, James Julian Ross, Andreas Hagmüller, Florian Pühringer, Pinelopi Pliota, Daniel Krogull, Valeria Stefania, Manuel Hunold, Alevtina Koreshova, Anja Koller, Ivanna Ostapchuk, Jacqueline Okweri, Joseph Gokcezade, Peter Duchek, Gang Dong, Eyal Ben-David, Alejandro Burga

## Abstract

To sustain life, molecular complexes require the concerted action of multiple proteins, each relying on one another to perform intricate tasks. However, how such interdependent protein interactions evolve in the first place is poorly understood. To address this, we investigated the origins of a group of fast-evolving genetic parasites—toxin-antidote elements—which boil down this dilemma to a simple question: what came first, the toxin or the antidote? By integrating quantitative genetics, biochemistry, and evolutionary genomics, we discovered that toxins and antidotes can arise simultaneously through the duplication of a regulatory module comprising an F-box protein in linkage to its substrate. Our findings provide one solution to the recurrent emergence of mutual dependence in protein complexes and illustrate in detail how complexity can swiftly arise from simplicity.

## Main text

Organisms must constantly innovate to survive. While some proteins are capable of stunning biochemical feats on their own, most pivotal innovations—from the bacterial flagellar motor to the eukaryotic nuclear pore complex—require the concerted action of hundreds of them. Understanding how natural selection has given rise to such complex molecular machines through small and gradual changes is one of the major challenges in evolutionary biology. However, despite significant advances, the underlying principles are still poorly understood (*1–3*). A critical barrier remains the sheer number and intricacy of interactions between proteins, which often exhibit mutual dependence, with proteins relying on one another for proper functioning. As a result, it’s often unclear which components came first and which followed. In addition, many of these innovations are ancient, dating back hundreds of millions of years, and key intermediate steps are now extinct, making it difficult to infer their precise evolutionary path. While ancestral sequence reconstruction has offered invaluable insights by resurrecting some of these lost intermediates, this approach is generally restricted to conserved and abundant protein families (*1*, *2*, *4*). To overcome these limitations, we focused instead on a group of fast-evolving genetic parasites that succinctly encapsulates the evolution of mutual dependence and innovation: toxin-antidote elements.

Toxin-antidote elements (TAs) are selfish genes that increase their frequency in populations by poisoning individuals or gametes that did not inherit them (*5–7*). As their name suggests, TAs typically comprise two genes: a toxin and its cognate antidote. TAs are analogous to prokaryotic toxin-antitoxin elements and retrons, which mediate plasmid-addiction and phage resistance (*8–10*). Toxins and antidotes exhibit antagonistic activities, but when genetically linked, they cooperate, revealing their selfish behavior. In crosses between TA carrier and non-carrier strains, heterozygous mothers load the toxin into all their eggs. However, only progeny that inherit the TA can counteract the maternal toxin by zygotically expressing its antidote (Fig. 1A). Thus, a single TA can poison up to 25% of the F_2_ progeny and this selective removal of homozygous non-carriers leads to the fixation of the TA in the population (Fig. 1A). Despite their simplicity, the evolution of these vicious genetic parasites from host genomes poses an evolutionary conundrum. Because the toxin is lethal in the absence of its antidote and the antidote serves no apparent purpose other than neutralizing the toxin, this begs the question—what came first, the toxin or the antidote? Here we explore this causal paradox by dissecting how an enzyme that is universally essential for life—the phenylalanyl-tRNA synthetase—gave birth to three distinct selfish TAs in the nematode *Caenorhabditis tropicalis*.

**Figure 1.**
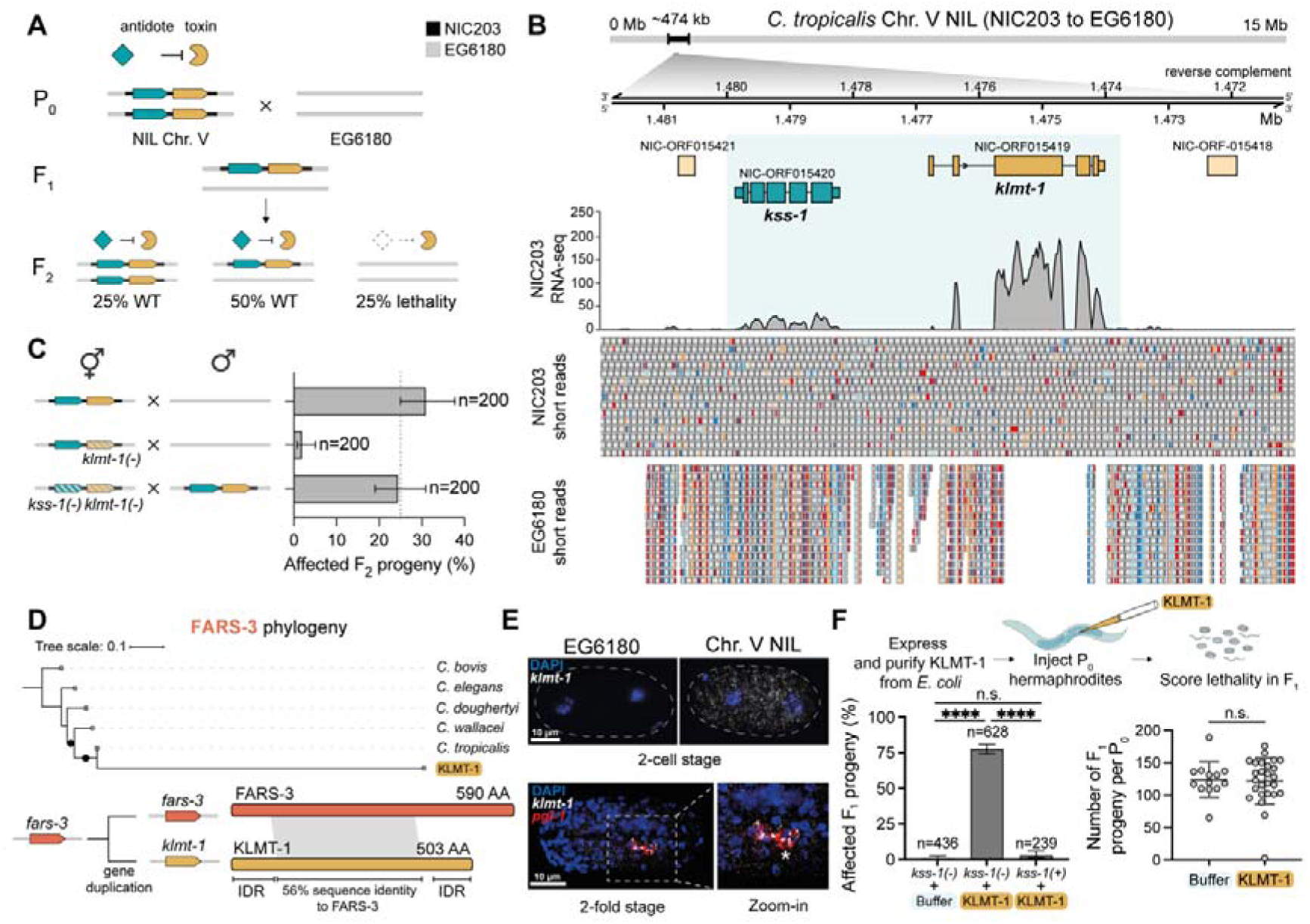
*klmt-1/kss-1* is a maternal-effect toxin-antidote element. **(A)** Mechanism of action of a toxin-antidote (TA) element. In crosses between the strain carrying the TA (Chr. V NIL, NIC203 to EG6180 background) and the susceptible strain (EG6180), all their F_2_ progeny are initially poisoned by a maternally provided toxin. Homozygous individuals for the susceptible allele (25% of the F_2_) die during embryonic development because these embryos lack the TA and cannot express the zygotic antidote. EG6180 genetic background is coloured in gray, NIC203 introgression in black. **(B)** Fine-mapping of the genes underlying the Chr. V TA. Alignment of genomic short reads and mRNA-seq from young adults to the NIC203 *de novo* assembly (see fig. S1C for corresponding locus in EG6180). Highlighted in light blue are the two top candidate genes: *kss-1 (ORF015420)* and *klmt-1 (ORF015419)*. **(C)** The Chr. V TA kills ∼25% of the F_2_ progeny (top). A frameshift mutation in *klmt-1* is sufficient to abolish this lethality (middle). A *kss-1(-) klmt-1(-)* double mutant allele is poisoned by the WT allele (bottom). Error bars indicate 95% confidence intervals calculated with the hybrid Wilson/Brown method. **(D)** A phylogenetic tree of FARS-3 indicates that the KLMT-1 toxin is a fast-evolving paralog of *C. tropicalis* FARS-3 (top). Comparison of FARS-3 and KLMT-1 protein domains and shared homology (bottom). Intrinsically disordered region (IDR). **(E)** *klmt-1* expression prior to zygotic genome activation and during late embryogenesis in Chr. V NIL by smFISH. *pgl-1* is expressed in germ cells. EG6180 serves as a negative control for *klmt-1* probes. **(F)** KLMT-1 protein purified from *E. coli* kills embryos when injected into the gonad of hermaphrodites, but embryos expressing the KSS-1 antidote from a single copy transgene are protected from KLMT-1 toxicity (*P* < 0.0001 for injection of KLMT-1 vs buffer, and for injection of KLMT-1 into EG6180 vs *kss-1(+)* strain; *P* = 0.0597 for buffer injection vs KLMT-1 injection into *kss-1(+)* strain, Fisher’s exact test). At least 8 hermaphrodites were injected per condition. Error bars indicate 95% confidence intervals calculated with the hybrid Wilson/Brown method (left). KLMT-1 does not affect the fecundity of injected mothers (two-sided unpaired t-test; *P* = 0.8542, n.s. – not significant). Error bars indicate mean with standard deviation (right).

### *klmt-1/kss-1* is a maternal-effect toxin-antidote element

TAs engage in sustained molecular arms races with their hosts. As a result, TAs typically code for fast-evolving proteins of unknown molecular function, making it difficult to trace their origins. To overcome this barrier, we set out to characterize novel TAs in the nematode *C. tropicalis*, where they cause extensive genetic incompatibilities (*7*, *11*). Through studying crosses between two *C. tropicalis* wild isolates from the Caribbean, NIC203 (Guadeloupe, France) and EG6180 (Puerto Rico, USA), we discovered that NIC203 carries three TAs located in Chr. II, III, and V, respectively (*7*). Each of these TAs poisons individuals that are homozygous for the corresponding EG6180 susceptible allele. We previously identified the genes that comprise the Chr. III TA, which we named *slow-1/grow-1* (*12*). However, the genes that make up the Chr. II and V TAs were unknown.

To facilitate their identification and characterization, we took advantage of near-isogenic lines (NILs) each carrying a single TA (NIC203 introgression) in an otherwise EG6180 parental background. Due to the smaller size of the introgressed region, we first focused on mapping the Chr. V TA. In crosses between the Chr. V TA NIL and the EG6180 parental line, ∼25% of the F_2_ progeny dies during embryonic development or shortly after hatching (L1 stage) (*7*). In agreement with the TA model, the vast majority of these affected individuals were homozygous carriers of the EG6180 allele (91.3%, n=92; Fig. 1A, fig. S1A, records for all crosses available in Data S1). To identify the genes underlying this incompatibility, we leveraged Nanopore long-read de novo genome assemblies and RNA-seq expression data to define a list of candidate loci within the ∼474 kb (1.26–1.74 Mb) NIC203 introgression. Among the 176 genes we found within this region, we sought gene pairs that were either (i) absent, mutated or divergent in EG6180 and (ii) in tight genetic linkage. These criteria were matched by one pair of genes: *NIC-ORF015419* and *NIC-ORF015420* (Data S2).

Since toxins are typically expressed at higher levels than antidotes (fig. S1B), we hypothesized that *ORF015419* is the toxin-encoding gene (Fig. 1B). To test this, we first generated an *ORF015419* knockout allele in the Chr. V NIL background using CRISPR/Cas. The mutation introduced a frameshift and premature stop codon resulting in a truncated protein (fig. S1D). *ORF015419*(*-*) mutants were phenotypically wild-type (95%, n=100) indicating that *ORF015419* is dispensable for the normal development of *C. tropicalis*. To test if *ORF015419* is the toxin, we crossed *ORF015419*(*-*) NIL hermaphrodites to EG6180 males and checked whether the TA was still active. We observed background levels of embryonic lethality among the F_2_ progeny (2%, n=200) in contrast to the ∼25% affected progeny in the wild type cross (31%, n=200) (Fig. 1C). Furthermore, F_2_ progeny homozygous for the susceptible allele (EG/EG) were recovered at the expected Mendelian ratio and were perfectly viable and fertile (26.5%, n=200). Thus, we concluded that *NIC-ORF015419* codes for the toxin and named the novel locus *klmt-1* (for *<u>K</u>iller of embryos and <u>L</u>arvae <u>M</u>aternal <u>T</u>oxin;* pronounced *klimt)*.

Next, we determined that the gene immediately upstream of *klmt-1*, *NIC-ORF015420*, encodes its antidote, which we named *kss-1* (for *<u>K</u>lmt rescue by zygotically expre<u>SS</u>ed gene*; pronounced *kiss*). If *klmt-1* and *kss-1* make up the TA, then we would expect the *klmt-1*(*-*) *kss-1*(*-*) double mutant haplotype to phenocopy a susceptible EG6180 allele, which lacks both. To show this, we mutated *kss-1* in the background of the mutant toxin using CRISPR/Cas (fig. S1D) and then crossed *klmt-1*(*-*) *kss-1*(*-*) double mutant hermaphrodites to wild type NIL males. Consistent with *kss-1* being the antidote, we observed ∼25% embryonic/L1 lethality among the F_2_ progeny (24.5%, n=200), and all genotyped individuals homozygous for the double mutant allele were embryonic or larval lethal (n=17) (Fig. 1C). As a control, only background levels of lethality were observed in the double mutant NIL parental strain (4%, n=100). Furthermore, overexpression of KSS-1 was sufficient to suppress the genetic incompatibility and prevented the death of F_2_ embryos homozygous for the susceptible allele (4.3%, n=23; fig. S1E-F). Thus, *klmt-1/kss-1* is a novel maternal-effect TA.

### The *klmt-1* toxin arose from *fars-3* through gene duplication

The toxin, *klmt-1,* codes for a 503-amino-acid protein and its closest paralog is the highly conserved and essential gene, *fars-3*, a subunit of the phenylalanyl tRNA-synthetase (PheRS) (Fig. 1D and fig. S2A). In bacteria, archaea and eukaryotes, PheRS is a heterotetrameric enzyme made up of two alpha and two beta subunits, which in nematodes are encoded by *fars-1* and *fars-3*, respectively (*13*, *14*) (fig. S2B). Like all other aminoacyl-tRNA synthetases, PheRS plays a critical role in mRNA translation, as it is responsible for charging tRNA^Phe^ with its cognate amino acid, L-phenylalanine. FARS-3 has two main domains: N-terminal (AA:1-376) and C-terminal (AA:390-590), which are connected by a short linker (fig. S2C). Phylogenetic analyses of FARS-3 across *Caenorhabditis* species indicated that KLMT-1 arose from a recent gene duplication event (Fig. 1D). However, KLMT-1 lacks the FARS-3 C-terminal domain and is instead predicted to have intrinsically disordered domains on its N- and C-terminal regions (Fig. 1D and fig. S2D). In agreement with its role as a maternal-effect toxin, we detected *klmt-1* mRNA expression in NIC203 embryos prior to zygotic genome activation (<4-cell stage) using single-molecule fluorescent in situ hybridization (smFISH). At the comma stage, *klmt-1* mRNA was only detectable in a pair of cells corresponding to the Z2 and Z3 germline precursors (Fig. 1E). As a negative control, we did not detect *klmt-1* mRNA in EG6180 embryos (Fig. 1E, fig. S3A).

To study the localization of KLMT-1 at the protein level, we introduced a C-terminal 3xFLAG tag at the *klmt-1* endogenous locus in the NIL background using CRISPR/Cas homology-mediated repair. The resulting KLMT-1::3xFLAG toxin could be readily detected by western blot and closely matched its predicted molecular weight of 59.3 kDa (fig. S3B). To test whether KLMT-1::3xFLAG was active, we crossed *klmt-1::3xflag* NIL hermaphrodites to EG6180 males and scored their F_2_ progeny. We observed 24.5% of affected F_2_ progeny (n=200) and the majority of affected progeny were homozygous EG/EG embryos (87.5%, n=24; fig. S3C). Immunofluorescence staining revealed that KLMT-1::3xFLAG protein is maternally loaded into eggs prior to fertilization (fig. S3D). KLMT-1 protein levels quickly declined during embryogenesis and could only be detected in the germ cell precursor cells following the mid gastrulation stage (fig. S3E). We observed the same expression pattern when staining embryos with a KLMT-1 monoclonal antibody (fig. S3D-F).

Since both *klmt-1* mRNA and protein are maternally loaded into eggs, we next asked whether maternal KLMT-1 protein is sufficient to poison embryos. To do this, we first expressed KLMT-1 in *E. coli*, purified it using a 6xHis-tag, and confirmed its identity using liquid chromatography mass spectrometry (LC-MS) (fig. S4A). We then co-injected the purified KLMT-1 protein and a plasmid-encoded fluorescent marker into the gonad of EG6180 hermaphrodites and characterized in detail their progeny following selfing. Analogously to the toxicity caused by the *klmt-1/kss-1* TA in genetic crosses, injection of purified KLMT-1—but not the vehicle buffer—caused high levels of embryonic and L1 lethality among their F_1_ progeny (77.7%, n=628, table S2, Fig. 1F). To test whether the toxicity was specifically caused by KLMT-1, we injected the purified toxin into a line overexpressing KSS-1 from a single copy transgene. We observed only background levels of embryonic lethality in the presence of its antidote (2.9%, n=239, table S2) indicating that KLMT-1 was responsible for the toxicity (Fig. 1F). Despite the drastic difference in F_1_ survival following KLMT-1 injection, injected mothers showed no obvious physiological problems or differences in the number of eggs laid (Fig. 1F), suggesting that KLMT-1 does not affect the fertility of adult mothers. We conclude that duplication of the essential gene *fars-3* gave rise to *klmt-1*, a selfish toxin maternally loaded into eggs.

### KSS-1 is an F-box protein that binds KLMT-1 and recruits the SCF complex

In prokaryotes, antitoxins utilize various mechanisms to neutralize toxins. For instance some antitoxins are antisense RNAs that block toxin synthesis, while others are proteins that bind to the toxin, preventing it from interacting with its target (*15*, *16*). Since the mechanisms by which eukaryotic antidotes neutralize toxins are largely unexplored (*17*), we next investigated how *kss-1* neutralizes *klmt-1*. KSS-1 is a predicted 389 AA long protein. An InterPro search revealed that the central region of KSS-1 [AA:201-252] contains a nematode-specific domain, IPR012885, which is typically found in F-box proteins, but does not provide evidence for the presence of the F-box domain itself (fig. S5A). F-box proteins are the substrate recognition subunits of the SKP1–CUL1–F-box (SCF) E3 ubiquitin ligase complexes, which target proteins for proteasome-mediated degradation.

To better understand the molecular role of KSS-1, we predicted its tertiary structure using AlphaFold2 (*18*). The resulting model had a high accuracy and revealed that KSS-1 adopts a horseshoe fold with parallel beta sheets running on its inner face and alpha helices on its exterior (Fig. 2A and fig. S5B- D). We then used the AlphaFold2 KSS-1 model as a template to search for proteins with a similar structure in the Protein Data Bank (PDB) (*19*). The repetitive solenoid arrangement bore a striking structural resemblance to the Leucine-Rich-Repeat (LRR) domain found in F-box proteins such as S-Phase Kinase Associated Protein 2 (SKP2; z-score 7.8) and *Arabidopsis* Coronatine-insensitive protein 1 (COI1; z-score 8.4) (Fig. 2A and fig. S5E). Furthermore, the N-terminal F-box domain of KSS-1 adopted a very similar fold to the N-terminal F-box domain of SKP2 and COI1 despite sharing only 18% and 17% sequence identity, respectively (Fig. 2B and fig. S5E).

**Figure 2.**
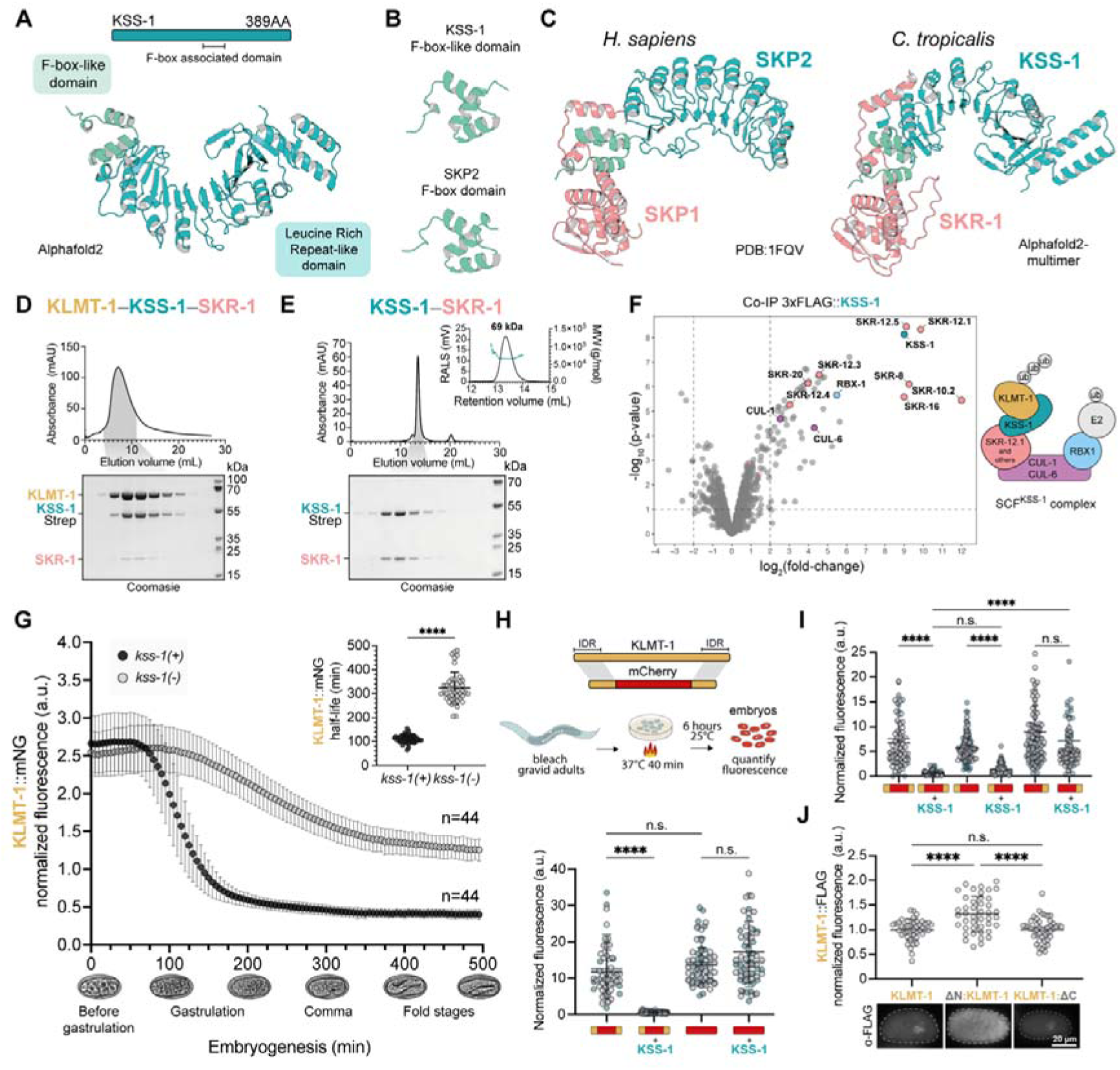
KSS-1 is an F-box that binds KLMT-1 and mediates its degradation. **(A)** Predicted protein domains and AlphaFold2 model of KSS-1. **(B)** Comparison of F-box domain from human SKP2 (PDB:1FQV) and predicted F-box-like domain from KSS-1. **(C)** Comparison of human SKP1– SKP2 protein complex (PDB:1FQV) and predicted by Alphafold2-multimer *C. tropicalis* SKR-1– KSS-1 complex. **(D)** Strep-affinity purification of KLMT-1–KSS-1::Strep–SKR-1 complex. KSS-1::Strep was co-expressed in Sf9 insect cells together with untagged versions of KLMT-1 and SKR-1 (top). Elution fractions from the StrepTrapXT column were visualized by SDS-PAGE and Coomassie staining (bottom). **(E)** SEC elution profile of the KSS-1–SKR-1 complex (top). SEC elution fractions from the Superdex 200 Increase 10/300 GL column were visualized by SDS-PAGE and Coomassie staining (bottom). Static Light Scattering (SLS) determination of the molecular weight of the KSS-1– SKR-1 complex. The molecular weight of the complex (69 kDa) closely matches that of a 1:1 heterodimer (47.1+19.8 kDa) (inset). **(F)** Volcano plot of 3xFLAG::KSS-1 co-immunoprecipitation followed by mass spectrometry (left). Members of the SCF complex are color-coded according to their function in the complex (right). Co-IPs were performed in biological triplicates. Lysate from Chr. V NIL served as a control for fold enrichment. **(G)** Quantification of endogenous KLMT-1::mNG levels during embryonic development in the presence or absence of KSS-1. Fluorescence quantification starting time corresponds to the 8 to 16-cell stages. KLMT-1::mNG fluorescence of both strains is normalized to mean fluorescence of INK169, *kss-1(+)*, strain. Error bars indicate mean with standard deviation. Half-life of the toxin is extended in the absence of the antidote (exact Wilcoxon rank sum test, *P* < 2.2^-16^). Error bars indicate mean with standard deviation (inset). **(H)** Fluorescent reporter assays to identify the KLMT-1 region necessary for KSS-1-mediated degradation. Fluorescent mCherry reporters are expressed under the control of a heat-shock inducible promoter (*Ctr-hsp-16.11p*), whereas the antidote is expressed under a constitutive promoter (*Ctr*-*rpl-36p*). Reporter and antidote genes were integrated as single copy transgenes in synthetic landing pads on Chr. IV and Chr. I, respectively. Fusing KLMT-1 N-and C-terminal IDRs to mCherry is sufficient for its specific degradation (*P_adj_* < 0.0001), while mCherry on its own is not degraded by KSS-1 (*P_ad_*_j_ > 0.9999). Fusing disordered regions to mCherry does not change its signal intensity (*P_adj_* = 0.52). Difference between group means was analyzed using Kruskal-Wallis test (*H*(4) = 123.3, *P* < 0.0001), followed by Dunn’s post hoc test. Error bars indicate mean with standard deviation. Experiment was performed twice, individual repeats are coloured in gray and cyan. **(I)** Similar experiments as in (H) identify the N-terminal IDR of KLMT-1 (AA:1-69) as sufficient for KSS-1-mediated degradation. KSS-1 degrades mCherry fused to N-terminal IDR with similar efficiency as it does mCherry with both IDRs (*P_ad_*_j_ < 0.0001). Difference between group means was analyzed using Kruskal-Wallis test (*H*(6) = 306.8, *P* < 0.0001), followed by Dunn’s post hoc test. Error bars indicate mean with standard deviation. Experiment was performed twice, individual repeats are coloured in gray and cyan. **(J)** Immunofluorescence staining of endogenous KLMT-1. The N-terminal IDR of KLMT-1 is necessary for its degradation *in vivo* (*P_adj_* < 0.0001), unlike C-terminal IDR (*P_adj_* > 0.9999). KLMT-1 levels were quantified at the late gastrulation and bean stages using an anti-FLAG antibody. Difference between group means was analyzed using Brown-Forsythe ANOVA test (*F\**(2, 102.8) = 18.52, *P* < 0.0001), followed by Games-Howell post hoc test. Error bars indicate mean with standard deviation. For H–J n.s. means not significant. Representative embryos of late gastrulation stage (bottom).

SKP1 serves as a key adapter that links dozens of different F-box proteins to the scaffold protein CUL1, which in turn binds the ubiquitin conjugating enzyme (*20*, *21*). While humans and yeast possess a single SKP1 gene, nematodes have undergone a substantial expansion of this protein family. For instance, the genomes of *C. elegans* and *C. tropicalis* code for 20 and 23 SKP1-related (*skr*) genes, respectively (*22*, *23*) (fig. S6). Among these, *skr-1* and its close paralog *skr-2*, are essential for embryonic development in *C. elegans* (*22*, *23*). We used AlphaFold-Multimer to test whether C. *tropicalis* SKR-1 interacted with the antidote. We identified a high confidence interaction between the C-terminus of SKR-1 (alpha helices H5-H8) and the N-terminal F-box-like domain of KSS-1, analogous to the interaction found in SKP1–SKP2 and other known SCF complexes (Fig. 2C and fig. S7A-C) (*20*, *24*). The structural similarity between KSS-1 and F-box proteins, combined with the sharp decrease of KLMT-1 levels coinciding with KSS-1 expression in early embryos (fig. S7D), led us to hypothesize that KSS-1 targets KLMT-1 for degradation by binding the toxin and recruiting the SCF complex. To test this model *in vitro*, we co-expressed a C-terminally tagged KSS-1::Strep together with untagged versions of KLMT-1 and SKR-1 in Sf9 insect cells. Affinity purification of KSS-1::Strep followed by size exclusion chromatography revealed that KSS-1 stably binds both KLMT-1 and SKR-1 (Fig. 2D and fig. S7E). KLMT-1’s strong tendency to aggregate through its disordered regions prevented us from identifying the precise stoichiometry of this complex (fig. S7F); however, in agreement with SKP1–SKP2 structures (*24*), we found that KSS-1 and SKR-1 form a stable 1:1 heterodimer (Fig. 2E).

To test whether KSS-1 binds components of the SCF complex *in vivo*, we performed co-immunoprecipitation of a N-terminally tagged 3xFLAG::KSS-1 transgene followed by quantitative mass-spectrometry (Co-IP–MS). As expected, KSS-1 was highly enriched in the IP samples compared to a WT control line (∼500 fold-change; *P* < 10^-8^; Fig. 2F). In agreement with our model, we identified eight SKR proteins among the most significant KSS-1 interactors, many of which were closely related paralogs, such as SKR-10.2, SKR-12.1, and SKR-12.5 (∼500-4000 fold-change; *P* < 10^-5^; Fig. 2F, fig. S6, and data S3). We independently validated a direct interaction between KSS-1 and *Ctr*-SKR-20 using yeast two-hybrid assay (fig. S7G). Furthermore, two cullin scaffolds, CUL-1 and CUL-6—the two closest worm paralogs to human CUL1—were also enriched in the IP samples (∼5-fold-change and ∼20-fold-change, respectively; *P* < 10^-4^; Fig. 2F), as was RBX1 (∼45-fold-change; *P* < 10^-5^; Fig. 2F). CUL1 is a core component of the SCF complex that together with RBX1 recruits the E2 ubiquitin-conjugating enzyme (*25*). Since differences in *klmt-1* mRNA levels between *kss-1*(*-*) and WT embryos could not account for the differences at the protein level (fig. S7H), overall, our results strongly suggest that KSS-1 exerts its antidote activity post-transcriptionally by binding KLMT-1, recruiting the SCF complex, and targeting KLMT-1 for degradation.

### KSS-1 mediates KLMT-1 degradation via an N-terminal degron

To test whether KSS-1 targets KLMT-1 for degradation in embryos, we first built a reporter strain carrying a C-terminal mNeonGreen (mNG) tag in the *klmt-1* endogenous locus. In agreement with KLMT-1 immunostaining, we observed strong fluorescent signal in unfertilized eggs and early embryos (fig. S8A). Furthermore, KLMT-1::mNG was present in the gonads of adult hermaphrodites consistent with its maternal deposition (fig. S8A). While the addition of this fluorescent tag abrogated KLMT-1 toxicity (2.2% lethality in F_2_, n=90) (fig. S8B), this inactive reporter allowed us to quantify KLMT-1 levels in *kss-1*(*-*) embryos, which would otherwise be dead. In line with this, we generated a *kss-1*(*-*) null allele in the *klmt-1*::*mNG* background using CRISPR/Cas and the resulting strain was healthy and viable (94% wild type, n=100) (fig. S8C). Next, we quantified the expression dynamics of KLMT-1::mNG in both *kss-1*(*-*) and WT *kss-1(+)* embryos. In agreement with our hypothesis, we found that KLMT-1::mNG protein levels were largely stabilized in *kss-1*(*-*) embryos leading to a 2.9- fold increase in the toxin’s half-life (Fig. 2G). While KLMT-1::mNG was no longer detectable in the soma of WT embryos by the comma stage, *kss-1*(*-*) embryos retained somatic KLMT-1::mNG expression throughout embryogenesis; even during advanced L2/L3 larval stages (fig. S8D). Reintroduction of WT antidote activity via a single copy transgene restored prompt KLMT-1::mNG degradation, fully rescuing the *kss-1*(*-*) mutant phenotype (fig. S8C).

Next, we investigated how KSS-1 recognizes the toxin. We hypothesized that the disordered domains of KLMT-1 are crucial for this interaction, as they are absent in its paralog FARS-3 (fig. S2C). To test this, we designed a chimeric construct by fusing both N-terminal and C-terminal disordered domains of KLMT-1 to mCherry (AA:1-69 and AA:460-503, respectively) (Fig. 2H; fig. S2E). We then quantified the abundance of the chimeric protein in embryos in the presence and absence of KSS-1. Our results showed that the disordered domains are sufficient to target mCherry for degradation by KSS-1 (Fig. 2H). As a negative control, KSS-1 had no effect on the expression levels of wild-type mCherry (Fig. 2H). Next, we tested whether having only one disordered domain attached to mCherry is sufficient for degradation and found that the N-terminal disordered region of KLMT-1 was sufficient for KSS-1-mediated degradation (Fig. 2I).

To test whether the N-terminal disordered region was also necessary for degradation, we deleted either the N- or C-terminal disordered domains of *klmt-1* at its endogenous locus (fig. S9A). We confirmed expression of the truncated toxins, ΔN::KLMT-1 and KLMT-1::ΔC, via western blot (fig. S9B), and genetic crosses revealed that these were no longer toxic (fig. S9C). Immunofluorescent staining of KLMT-1 in embryos showed that the expression dynamics of KLMT-1::ΔC were indistinguishable from those of the WT toxin (Fig. 2J). In contrast, ΔN::KLMT-1 persisted at high levels in late embryonic stages, mimicking *kss-1*(*-*) mutants (Fig. 2J). These results indicate that the N-terminal domain corresponds to a degron, which is both necessary and sufficient for KSS-1- mediated degradation. Altogether, *in vitro* and *in vivo* evidence strongly suggests that KSS-1 functions similarly to canonical F-box proteins, binding the N-terminal disordered region of KLMT-1 and targeting the toxin for degradation via an SCF E3 ligase complex.

### The Chr. II TA is evolutionarily related to *klmt-1/kss-1*

Having identified the genes underlying the Chr. V TA, we set out to map the toxin and the antidote responsible for the incompatibility in Chr. II. Using the Chr. II NIL and a similar strategy as before (see Methods and Data S4), we found that two neighboring genes, *NIC-ORF006816* and *NIC- ORF006815*, encode the toxin and its antidote, respectively (Fig 3A and fig. S10A). The toxin codes for a predicted 90 kDa protein derived from the fusion of three different genes: *mec-15*, *zyg-9*, and— to our surprise—again*, fars-3* (Fig. 3B–C and fig. S10B). Thus, we named the new toxin *pzl-1* (for *<u>P</u>heRS beta subunit, zyg-9, and mec-15 fusion maternal toxin,* pronounced *puzzle*). We tagged the endogenous *pzl-1* with a C-terminal 3xFLAG using CRISPR/Cas and confirmed the presence of a protein of the expected molecular weight (Fig. 3D). Although both KLMT-1 and PZL-1 originated from FARS-3 through gene duplication, their domain architectures differ significantly. KLMT-1 shares homology with only the N-terminal domain of FARS-3, while PZL-1 is homologous to the C-terminal domain (Fig. 1D and 3B). This non-overlapping homology suggests distinct mechanisms of action for these toxins. Supporting this, KLMT-1 overexpression was lethal only when induced during early embryonic stages, whereas PZL-1 overexpression was lethal at all embryonic and early larval (L1, L2) stages (Fig. 3E). Furthermore, the resulting phenotypes differed: KLMT-1 poisoning produced small, crumpled L1 larvae, whereas PZL-1 overexpression resulted in bloated L1 larvae (fig. S10C).

**Figure 3.**
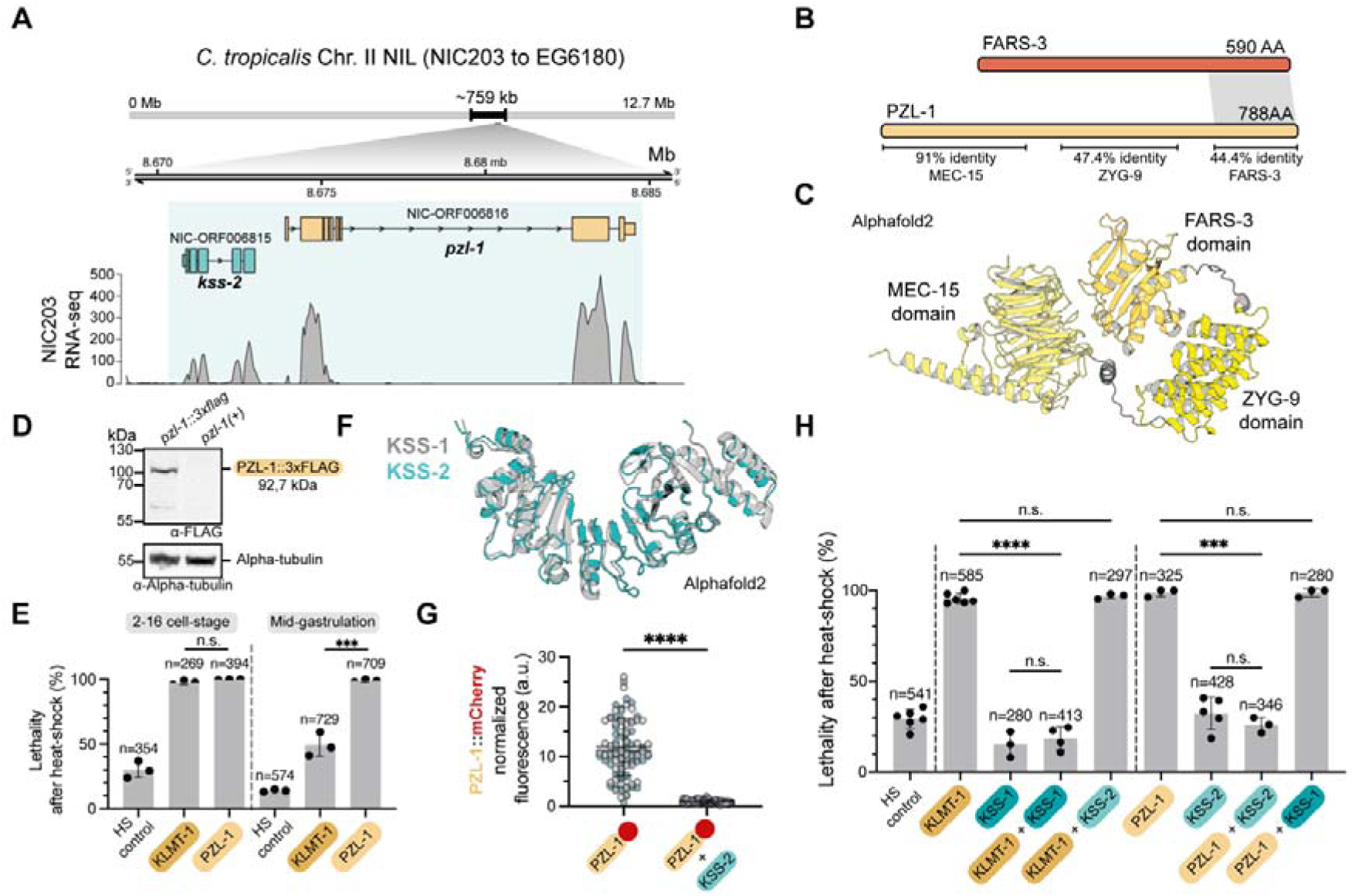
PZL-1, like KLMT-1, evolved from FARS-3, an essential tRNA synthetase subunit. **(A)** Fine-mapping of the TA element present in the NIC203 Chr. II NIL. The TA comprises the toxin *pzl-1* and its antidote *kss-2*. Alignment of mRNA-seq from early embryos to the NIC203 *de novo* assembly. **(B)** Comparison of FARS-3 and PZL-1 protein domains and homology. PZL-1 is a chimeric protein with homology to three conserved proteins: MEC-15, ZYG-9, and FARS-3. Percent identity value is calculated based on the local alignment. **(C)** Predicted AlphaFold2 model of the PZL-1 toxin. **(D)** The endogenous *pzl-1* locus was tagged with a C-terminal 3xFLAG (INK577) using CRISPR/Cas. A band of the expected molecular weight (92.7 kDa) was confirmed via western blot. Chr. II NIL serves as a negative control. Western blot against alpha-tubulin serves as a loading control. **(E)** Comparison of KLMT-1 and PZL-1 overexpression phenotype in early (up to 16 cell stage) embryos and after 3 hours of development, corresponding to mid-gastrulation stage embryos. Toxins are expressed from a single copy transgene in a synthetic landing pad on Chr. IV under the control of a heat-shock inducible promoter (*Ctr-hsp-16.11p*). Both toxins kill early embryos (two-sided unpaired t-test; *P* = 0.0737), however, unlike PZL-1, KLMT-1 is less potent during later stages of embryonic development (two-sided unpaired t-test; *P* = 0.0008). Each dot represents an individual replicate. At least 50 embryos were screened per replicate. Error bars indicate mean with standard deviation. **(F)** Protein alignment of KSS-1 (gray) and KSS-2 (cyan) Alphafold2 models. **(G)** Fluorescent reporter assay in embryos. KSS-2 overexpression leads to reduced PZL-1::mCherry expression levels (two-sided unpaired Welch’s t-test; *P* < 0.0001). Fluorescent toxin is expressed from a single copy transgene in a synthetic landing pad on Chr. IV under the control of a heat-shock inducible promoter (*Ctr-hsp-16.11p*), whereas the antidote is expressed under a constitutive promoter (*Ctr*-*rpl-36p*). Experiment was performed twice, individual repeats are coloured in gray and cyan. Error bars indicate mean with standard deviation. **(H)** Functional assays testing the rescue activity and specificity of KSS-1 and KSS-2 antidotes. Toxins and antidotes are under the control of a heat-shock inducible promoter (see (E)), expressed as single copy transgenes from synthetic landing pads on Chr. IV and Chr. I, respectively. Heat shock treatment in WT worms causes ∼30% embryonic lethality in early embryos (HS control), which is considered the background level in these experiments. Only cognate antidotes rescue from KLMT-1 and PZL-1 toxicity (*P_adj_* < 0.0001 for both). Overexpression of antidotes alone does not differ from overexpression of toxins with their cognate antidotes (n.s. – not significant, *P_adj_* > 0.48). Difference between group means was analyzed using Brown-Forsythe ANOVA test (for KLMT-1 comparisons – *F\**(3, 5.195) = 315.5, *P* < 0.0001; for PZL-1 comparisons – *F\**(3, 6.903) = 222.6, *P* < 0.0001), followed by Dunnett post hoc test. At least 50 embryos were screened per replicate. Error bars indicate mean with standard deviation.

The evolutionary connection between these two TAs extends beyond *fars-3*, as their antidotes are also homologous. ORF006815 and KSS-1 share 64,6% sequence identity at the protein level and their predicted structures are highly similar (RMSD 1.6 Å; Fig. 3F). Thus, we named the new antidote KSS-2. Co-IP experiments revealed that KSS-2—like KSS-1—interacts with components of the SCF complex (fig. S10D and Data S5). Further, we found that KSS-2 overexpression led to a drastic reduction in the levels of a PZL-1::mCherry reporter and identified a natural variant in a solvent-exposed residue of KSS-2, Arg297, which led to decreased clearance of the toxin (Fig. 3G and fig. S10E-G). Altogether, these results strongly suggest that, analogously to the *klmt-1/kss-1* TA, KSS-2 binds PZL-1 and targets it for degradation. While very similar, KSS-1 and KSS-2 are highly specific. In controlled transgenic overexpression experiments, these antidotes could efficiently neutralize their cognate toxins but conferred no cross-resistance (Fig. 3H). Thus, *klmt-1/kss-1* and *pzl-1/kss-2* are related but distinct TAs.

### The *fars-3* locus harbors a cryptic TA in *C. tropicalis*

Intrigued by the recurrent ability of *fars-3* to give rise to genetic parasites, we examined the *C. tropicalis fars-3* locus in closer detail. In *C. elegans*, *fars-3* (Chr. II: 8.4 Mb) is part of a polycistronic operon along with its upstream neighbor *zyg-9* and downstream neighbors, *F22B5.10* and *M05D6.2* (*26*). Overall, the synteny of these four genes is highly conserved among *Caenorhabditis* nematodes including *C. tropicalis* and its sister species, *C. wallacei* (Bali, Indonesia) (Fig. 4A) (*27*). However, we found a large ∼34 kb insertion between *C. tropicalis zyg-9.2* and *fars-3* that is missing from *C. wallacei* and other representative *Caenorhabditis* species. Within this insertion, we identified three divergent paralogs of *fars-3*, including at least two copies that exhibited clear signs of pseudogenization (Fig. 4A). Remarkably, we also found several closely related paralogs with significant homology to *kss-1* and *kss-2*, which we collectively named *kss-3* (Fig. 4A, fig. S11A). The number of these paralogs varied depending on the isolate. For instance, EG6180 carried a cluster of four paralogs: *kss-3.1*, *kss-3.2*, *kss-3.3*, and *kss-3.4*, whereas NIC203 carried three: *kss-3.2*, *kss-3.3*, and *kss-3.4* (fig. S11B). All of these paralogs code for intact F-box proteins (fig. S11C), and RNA-seq profiling revealed a similar expression pattern to that of *kss-1* and *kss-2*, suggesting that they may act during early embryonic development (fig. S11D).

**Figure 4.**
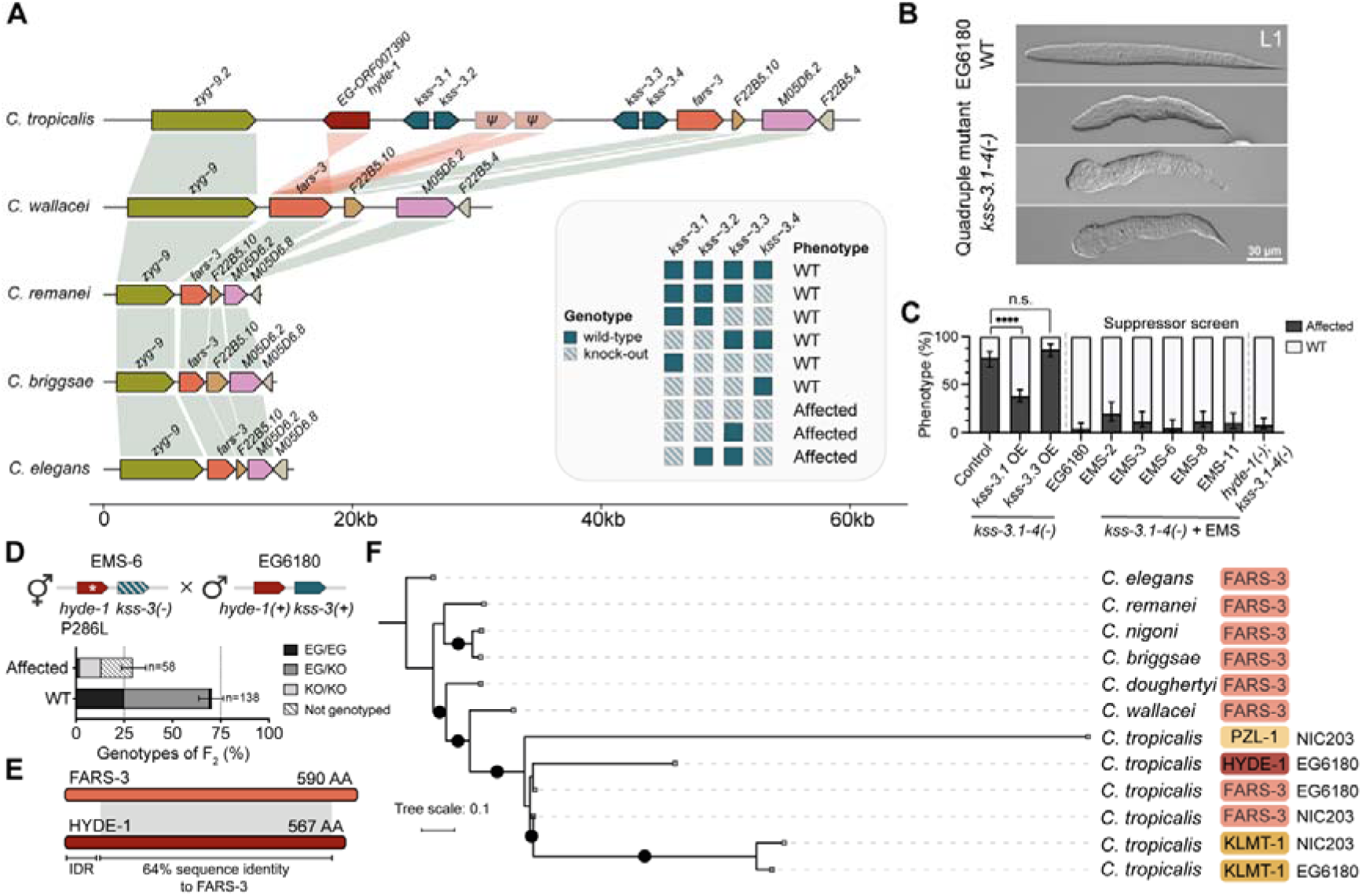
A cryptic TA in linkage to the *fars-3* parental locus. **(A)** Comparison of the *fars-3* locus across *Caenorhabditis* species. Orthologous genes are connected by shading. Paralogs with homology to *fars-3* are indicated by red shading. Four paralogs of *kss-1* and *kss-2* were identified in *C. tropicalis* EG6180 in between *zyg-9.2* and *fars-3*. Inset summarizes the genotype and phenotype of lines carrying various combinations of mutations in *kss-3* paralogs. **(B)** Morphological defects observed in L1 larvae lacking all four *kss-3* paralogs, denoted as *kss-3.1-4(-)*. EG6180 is a wild type reference. **(C)** Barplots summarizing the percentage of affected *kss-3.1-4(-)* embryos in various genetic backgrounds: i) overexpression (OE) of *kss-3.1* or *kss-3.3* (P < 0.0001 and P = 0.0967, respectively, Fisher’s exact test), ii) suppressor alleles recovered from an EMS mutagenesis screen (EMS-2, EMS-3, EMS-6, EMS-8, and EMS-11), and iii) a *ORF007390 (hyde-1)* null allele generated using CRISPR/Cas. Error bars indicate 95% confidence intervals calculated with the hybrid Wilson/Brown method. **(D)** Genetic cross between the suppressor line EMS-6 (INK1095) and the EG6180 wild isolate. EMS-6 carries a putative null mutation of *hyde-1*[P286L] as well as null mutations of all four *kss-3* paralogs. As expected if *hyde-1/kss-3* was a TA, ∼25% of their F_2_ progeny is affected and these individuals are homozygous for the susceptible haplotype. Error bars indicate 95% confidence intervals calculated with the hybrid Wilson/Brown method. KO/KO genotype corresponds to *hyde-1*[P286L] *kss-3.1-4(-).* **(E)** Comparison of FARS-3 and HYDE-1 protein domains and shared homology. Percent identity value is calculated based on the local alignment. **(F)** DNA-based phylogenetic tree showing the evolutionary relationship between *klmt-1*, *pzl-1*, *hyde-1,* and *fars-3*. Black dots denote branches with a bootstrap value (SH-aLRT) > 80%.

To gain insights into the function of the *kss-3* paralogs, and anticipating redundancy given their similarity, we systematically generated knockout alleles of these genes in the EG6180 background using CRISPR/Cas. Most double and triple knockout mutants did not display any obvious embryonic or larval phenotypes (Fig. 4A). However, double or triple mutant combinations missing *kss-3.1* and *kss-3.4*, as well as the knockout of all four *kss-3* paralogs, exhibited a high proportion of dead and sick progeny. For instance, in the quadruple mutant, we observed 57.2% of embryonic and larval lethality, and 20% of strong delay or poor fecundity (n=180) (Fig. 4A, Table S1). The phenotype of the surviving larvae included a wide range of anterior and posterior morphological defects and protrusions along the body axis, phenotypes that were never observed in embryos poisoned by KLMT-1 or PZL-1 (Fig. 4B). Overexpression of *kss-3.1*—but not *kss-3.3*— significantly rescued the mutant phenotype of the quadruple mutant (Fig. 4C), indicating that these morphological defects were specifically caused by the targeted disruption of *kss* paralogs rather than off-target mutations induced by CRISPR/Cas.

Having confirmed the specificity of the quadruple knock-out strain, we investigated why loss of *kss-3.1* and *kss-3.4* resulted in embryonic and larval defects. Given their sequence and structural similarity to KSS-1 and KSS-2, we hypothesized that these genes might also function as F-box proteins, degrading an unknown target. Considering that the *kss* paralogs are located immediately upstream of *fars-3*—echoing the architecture of *klmt-1/kss-1* and *pzl-1/kss-2* loci—we first tested whether these genes could degrade FARS-3. To do so, we quantified FARS-3 protein levels upon deletion or overexpression of *kss* paralogs using an endogenously tagged *fars-3*::*mScarlet* reporter line but found no significant differences compared to controls (fig. S11E–F). Next, we adopted an unbiased genome-wide approach. The quadruple knockout strain is very sick but viable; those few embryos that reach adulthood are usually fertile, allowing the mutant strain to be propagated indefinitely. Hence, we reasoned that a EMS-forward genetic screen could help us identify the putative target among its suppressors (*28*). To do this, we mutagenized the quadruple mutants, isolated their F_2_ progeny, and propagated them for approximately 10 generations. Reasoning that full or strong suppressors would outcompete the mutants and quickly increase their frequency in the population, we isolated single hermaphrodites from the fastest-growing mutagenized plates and established candidate suppressor lines from each of them. To validate these lines, we quantified their percent of embryonic and larval lethality, ultimately obtaining five independent suppressor lines, which closely resembled the wild-type line, EG6180 (Fig. 4C). Finally, we sequenced the whole genome of the parental and these five suppressors lines, identified EMS-derived *de novo* mutations, and predicted their coding impact. Remarkably, 4 out of 5 suppressor lines carried either non-synonymous or splicing-altering variants in the same gene: *EG-ORF007390* (Data S6).

*ORF007390* is one of the *fars-3* paralogs located downstream of *zyg-9.2* in *C. tropicalis* (Fig. 4A). To confirm that mutations in *ORF007390* suppressed the mutant phenotype, we generated *ORF007390*(*-*) null mutants using CRISPR/Cas (fig. S12A). The quadruple *kss-3* knockout strain was fully viable and healthy in an *ORF007390*(*-*) genetic background, thus validating the candidate suppressor (Fig. 4C). The discovery that *ORF007390* and the *kss-3* paralogs were together dispensable for embryonic development led to an unexpected revelation: ORF007390 was not a pseudogene but a toxin, and *kss-3.1* and *kss-3.4* were its cognate antidotes. In agreement with the TA model, we crossed EG6180 with the EMS-6 suppressor line and observed developmental defects in ∼25% of the F_2_ progeny and affected embryos were homozygous for the EMS-derived suppressor allele (Fig. 4D). The same pattern of F_2_ lethality was observed when crossing the suppressor EMS-3 with EG6180 and the CRISPR derived *ORF007390*(*-*) *kss-3.1-4*(*-*) mutant line with EG6180 (fig. S12B). Furthermore, reciprocal maternal and paternal backcrosses indicated that ORF007390 was a maternal-effect toxin, like *klmt-1* and *pzl-1* (fig. S12C). Based on these findings, we named the new toxin *hyde-1* (for *H*armful phen*Y*lalanine-tRNA synthetase *D*uplicat*E*). Unlike all other TAs, *hyde-1/kss-3.1,4* is tightly linked to its parental locus (Fig. 4A). As a result, *fars-3* haplotypes associated to an active *hyde-1/kss-3* TA are predicted to increase their frequency in populations due to genetic hitchhiking (*29*), showcasing a unique case of a selfish element conferring a selective advantage to its non-selfish parental gene.

HYDE-1 is 567 amino acids long and, unlike KLMT-1 and PZL-1, shares significant homology with both the N-terminal and C-terminal domains of FARS-3 (Fig. 4E, fig. S12D, and S13). Among the suppressors of *kss-3* loss of function, we identified a Pro286Leu substitution (line EMS-6). This proline is highly conserved across FARS-3 orthologs, and likely hinders the correct folding of the toxin (fig. S12E). Also, we found a Glu277Lys substitution (line EMS-3), which changes a negative charge for a positive one in a solvent exposed residue (fig. S12F). The parental NIC203 carries a divergent copy of *hyde-1*, which is truncated and not toxic (fig. S11B, S12G). The *hyde-1/kss-3-1,4* TA does not cause a genetic incompatibility in crosses between NIC203 and EG6180, likely because NIC203 *kss-3.4* expression is sufficient to neutralize HYDE-1 (fig. S11B, D). KSS-3.1 is 84.3% identical to KSS-1 but it does not confer resistance to KLMT-1. To gain insights into their substrate specificity, we exchanged a short region from a highly variable segment between KSS-1 and KSS-3.1 (fig. S14A) and found that this 6AA loop is necessary for the degradation of KLMT-1 and likely involved in substrate recognition (fig. S14B-D). Thus, we identified a third maternal-effect TA derived from the *fars-3* locus. Although related, the toxins from all three TAs appear to have different mechanisms, and the KSS antidotes are highly specific for their toxins.

### Duplication of the *kss/fars-3* module led to recurrent TA evolution

To better understand the evolutionary relationships between *klmt-1*, *pzl-1*, *hyde-1,* and *fars-3*, we constructed a DNA-based phylogenetic tree (Fig. 4F). To ensure accuracy and reliability, we leveraged sequence conservation and structural models to exclude predicted intrinsically disordered regions from the toxins. Further, for *pzl-1*, we included only the region homologous to *fars-3* (see Methods). The phylogeny revealed that all three toxins originated from *fars-3* following the divergence of *C. tropicalis* and its sister species, *C. wallacei*. Remarkably, the toxins did not form a monophyletic group, instead, we found strong statistical support for at least two independent duplication events of *fars-3*. The first duplication event gave rise to *pzl-1.* Subsequent duplications led to the emergence of either *klmt-1* and *hyde-1*, or their common ancestor; however, distinguishing between these models was not possible. The oldest age of the *pzl-1/kss-2* TA is also consistent with the fact that PZL-1 is the most derived toxin, including homology to MEC-15, which was likely a secondary acquisition (Fig. 3B).

For TAs to evolve *de novo*, a series of exceptionally rare events must take place. First, the toxin and the antidote must emerge simultaneously to avoid a causal paradox. Second, both genes must be genetically linked to enable their selfish behavior, and spread within populations. If they are found on different chromosomes or on opposite arms of the same chromosome, gene drive cannot take place. Despite the daunting odds, TAs are not only common in nematodes but a single essential gene, *fars-3*, duplicated at least twice, giving rise to three independent acting TAs in *C. tropicalis*. What is the basis for the evolution of these recurrent genetic parasites? To address this question, we examined the evolutionary origins of KSS antidotes in *Caenorhabditis* species. An exploratory screen for homologous sequences across 55 *Caenorhabditis* species revealed that these were primarily restricted to the *Elegans* group—with only a few significant matches in *C. sulstoni*, which belongs to the *Japonica* group. Thus, we selected 11 *Elegans*-group species with chromosome-level assemblies, as well as *C. sulstoni*, annotated genes homologous to KSS-1 using a curated hidden Markov model (HMM), and constructed a comprehensive phylogenetic tree (fig. S15). In total, we identified 560 homologs across 12 species, which we named *ksl* (for *<u>KS</u>*s-*<u>L</u>*ike genes) (Data S7). The phylogeny of KSL proteins revealed significant expansion and diversification of this family. Most KSL proteins were species-specific, a pattern similar to that observed in canonical F-box proteins from *C. elegans* and other nematodes (fig. S15) (*30*, *31*). The species with the highest number of KSL proteins were *C. remanei* (100), *C. latens* (98), and *C. doughertyi* (90). In contrast, *C. briggsae* and *C. inopinata* contained only 2 KSL proteins each, and no homologs were identified in *C. elegans*. The low number or absence of KSL homologs in these and other species could, in part, be explained by the rapid diversification of F-box proteins, which makes sequence-based identification challenging. Supporting this view, a structural-based search found 287 *C. elegans* proteins with significant similarity to KSS- 1 (fig. S14E and Data S8; Foldseek E-Value < 10^-5^).

Including known antidotes, we identified 30 KSL proteins in *C. tropicalis* (fig. S15). The most divergent residues across *C. tropicalis* KSL proteins were found in the inner surface of the solenoid, overlapping residues involved in toxin recognition, suggesting that their substrates are quickly evolving (fig. S14F). We also identified sister clades to the antidotes in *C. wallacei* and *C. doughertyi,* the two closest relatives to *C. tropicalis* (fig. S15) . Although representative KSL proteins, such as *Cwal-ksl-7* and *Cdou-ksl-30,* were only 35% and 28% identical to KSS-1, Alphafold2 models of these proteins revealed extensive structural homology to KSS-1, confirming their shared ancestry (Fig. 5A and fig. S15-16). Surprisingly, we found that *ksl* genes were not randomly distributed across all six equally-sized chromosomes: 78.4% (439/560) of *ksl* genes were located on Chr. II (Fig. 5A). This bias was present in all 10 species from the *Elegans* group where *ksl* were identified, including *C. tropicalis* (90%), *C. wallacei* (82.4%), *C. doughertyi* (90%). Further, *ksl* genes were not evenly distributed within Chr. II, but formed large clusters (Fig. 5B). Remarkably, these clusters were found in close proximity to their respective *fars-3* orthologs—approximately 1 Mb or less in the two closest relatives of *C. tropicalis* (Fig. 5B, Data S7). Altogether, these results strongly suggest that the *ksl* family originated in Chr. II and that genetic linkage between *fars-3* and the ancestor of all three antidotes predated the evolution of their selfish behavior.

**Figure 5.**
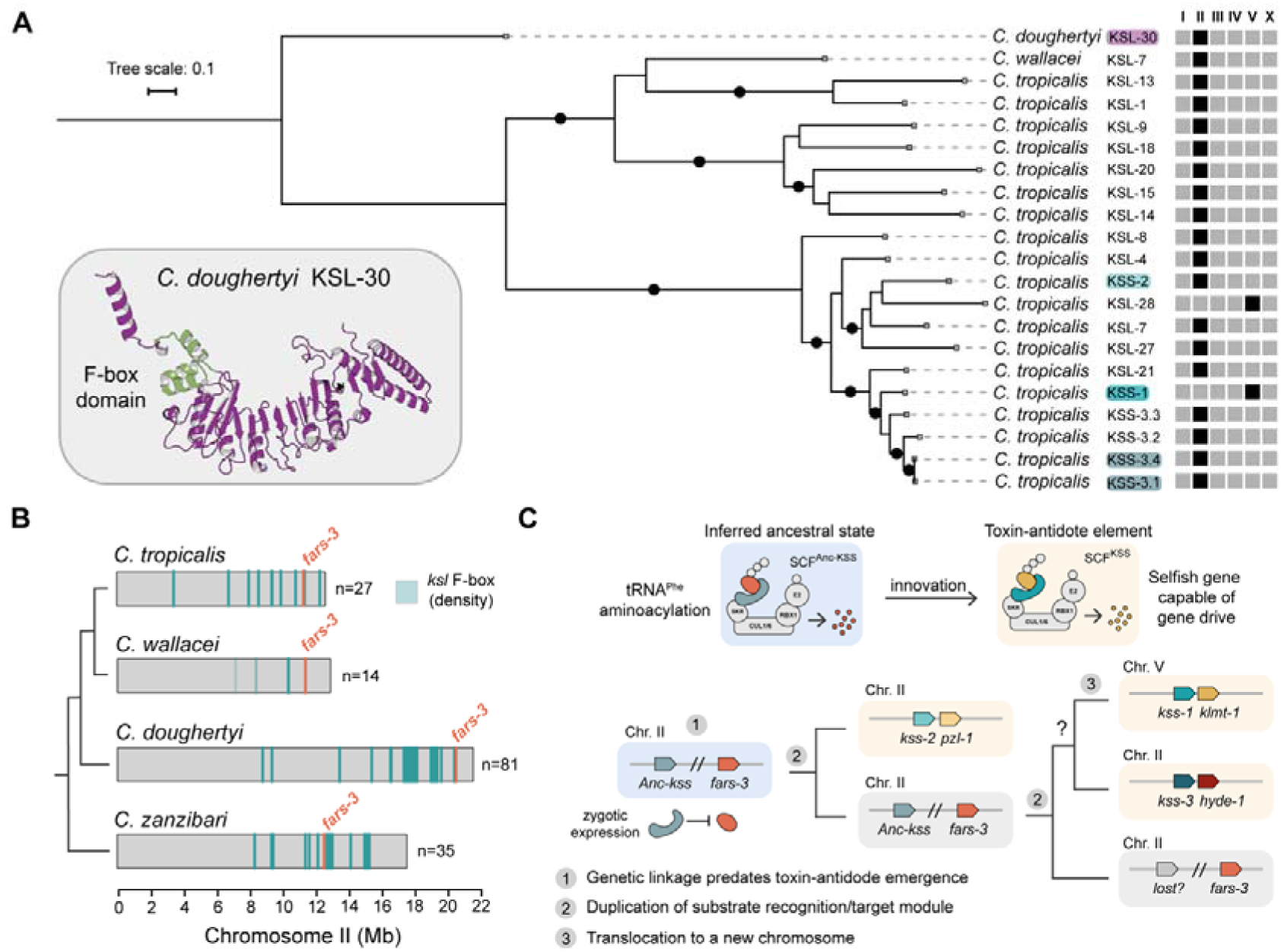
The emergence of KSS antidotes and a model for the recurrent evolution of TAs from *fars-3*. **(A)** Protein-based phylogenetic tree of the KSS antidotes, their closest paralogs in *C. tropicalis* and representative homologs of *C. wallacei* and *C. doughertyi*. *C.dou-*KSL-30 was designated as the outgroup. Black dots denote branches with a bootstrap value (SH-aLRT) > 80%. The chromosome where each KSS paralog is located is indicated in the diagram on the right. **(B)** Relative position of *fars-3* and *ksl* F-box genes on Chr. II across representative species of the *Elegans* group. Position of ksl genes is semi-transparent to emphasize the positions of clusters. N denotes the number of *ksl* genes in Chr. II. **(C)** A KSL F-box protein, part of a rapidly evolving cluster on Chr. II, randomly acquired affinity for FARS-3, becoming the ancestor of all three KSS antidotes (Anc-KSS). Anc-KSS was likely expressed exclusively in the early zygote and linked to FARS-3. After the duplication of the *fars-3*/Anc-*kss* genetic module and under relaxed selection, the divergent FARS-3 paralog became toxic, with the linked F-box protein simultaneously evolving into its antidote. Over time, increased toxicity and antidote specificity developed. The first duplication of the *fars-3*/Anc-*kss* module led to the emergence of *pzl-1/kss-2.* Subsequent duplications of this module gave rise to either the common ancestor of *klmt-1/kss-1* and *hyde-1/kss-3* or to each independently.

The phylogeny of the KSS antidotes mirrors that of the toxins and also supports the view that *pzl- 1/kss-2* was the first TA to emerge from the *fars-3* locus (Fig. 5A). Given that the ancestral *fars-3* gene duplicated at least twice, and considering that all three antidotes share a common evolutionary origin, there are two possible scenarios that could account for our observations: either ancestral KSS F-box proteins independently evolved the ability to bind different toxins on at least two occasions, or the binding evolved only once and involved the common ancestor of all these toxins, FARS-3 (fig. S17). Since the second scenario is the most parsimonious and given that linkage between the ancestral *kss* F-box proteins and *fars-3* predated the emergence of the TAs, our results strongly suggest that it was the duplication of the module consisting of both the ancestral F-box protein (Anc-*kss*) and *fars-3*, that drove the recurrent evolution of these selfish elements (Fig. 5C). A module consisting of a substrate-recognition protein and its target provides an ideal ground for the evolution of TAs. It only takes a copy of the substrate becoming slightly poisonous for the toxin and antidote functions to arise simultaneously, thereby escaping the causal paradox (Fig. 5C).

Given the essential role of FARS-3, how could Anc-KSS evolve an affinity for FARS-3 if doing so would result in its degradation? To explore this, we examined the expression patterns of *C. tropicalis* KSL paralogs (Fig. 5A). We found that 10 out of 13 *C. tropicalis* KSL paralogs (including also representatives of the more distant sister clade) are only transcribed during a small window in early development and show no detectable levels at the L4 stage, just like the KSS antidotes (Fig. 5A and fig. S16). This pattern strongly suggests that transient zygotic expression is the ancestral state of this protein family and did not evolve as a consequence of their role as an antidote. Moreover, this explains how the degradation of FARS-3 by Anc-KSS could have been tolerated. Since FARS-3 is constitutively expressed throughout all embryonic and larval stages, the ancestral F-box would have had only a transient effect on its dosage.

## Discussion

Much like new species emerge from a common ancestor through divergence and adaptation, complexes evolve from ancestral proteins that acquire new functions and interactions. Intrigued by the rate at which new species evolve, Eldredge and Gould proposed that—contrary to the prevailing gradualist view—species remain largely stable over extended geological periods (*32*, *33*). New species, they argued, arise instead in rapid bursts, leading to novel forms coexisting with the old, a process they termed punctuated equilibrium (*32*, *33*). Here, we provide evidence that this rapid differentiation tempo is mirrored at the molecular level by the evolution of *fars-3*. This essential gene did not show any obvious signs of neofunctionalization for hundreds of millions of years of nematode evolution. However, following the split of *C. tropicalis* and *C. wallacei*, *fars-3* duplicated multiple times and quickly gave rise to three selfish TAs. Like species occupying new niches (*34*, *35*), these TAs translocated to different genomic regions, breaking free from linkage, allowing for further diversification and specialization. But what ignited this frantic evolutionary sprint?

Recent studies indicate that, contrary to intuition, proteins are only one or a few substitutions away from evolving multimerization, allostery, and new folds—features associated with innovation and increased complexity (*36*). Our results further extend this view by showing that even two-component complexes exhibiting mutual dependence can evolve *de novo* in a few simple steps (Fig. 5C). A key genomic feature that likely facilitated this process is the massive expansion and diversification of F-box proteins in nematodes, providing a large repertoire of protein binders on which natural selection can act. In agreement with this idea, the two TAs found in *C. elegans*—*sup-35/pha-1* and *peel-1/zeel- 1*—likely evolved by duplication of analogous modules (*6*, *37*). Using Alphafold2, we found that the antidote PHA-1 is structurally analogous to KSS-1 (fig. S18). Furthermore, the antidote ZEEL-1, is homologous to a conserved substrate-recognition subunit of the Cullin-2-RING E3 ubiquitin ligase complex (fig. S18) (*38*, *39*).

The involvement of substrate-recognition proteins in the evolution of TAs also extends to plants. For instance, HTA, the antidote of a TA causing pollen sterility in rice hybrids, also codes for an F-box protein (*40*). Furthermore, our model may explain the recurrent evolution of self-incompatibility loci (SI), which ensure plants are only fertilized by genetically diverse pollen (*41*). Analogous to TAs, SI loci achieve this through the combined action of cytotoxic enzymes tightly linked with detoxifying F-box proteins. Like nematodes, plants have also experienced a large expansion of F-box proteins (*31*, *42*), suggesting that the independent amplification of this gene family may have fueled the convergent evolution of these modules across kingdoms. Beyond plants and nematodes, many other species encode hundreds of substrate-recognition proteins of unknown function, which we anticipate could have roles beyond simply fine-tuning protein levels. In life’s constant struggle for survival, from species to selfish genes, complexity swiftly arises from simplicity, poised delicately at its edge, unseen.

## Acknowledgments

We thank members of the Burga lab for critical reading of the manuscript. Library preparation and sequencing were performed at the VBCF NGS Unit. Proteomics analyses were performed by the Proteomics Facility at IMBA using the VBCF instrument pool. Annotation of KSL proteins was done by Thomas Burkand and Alexander Schleiffer from the VBCF Bioinformatics core. We thank the Max Perutz Laboratories Monoclonal Antibody Facility for generating α-KLMT-1 antibody (clone 1A3-3E5). We thank IMP-IMBA-GMI Biooptics Core Facility for support during the project. Research in the Burga lab is supported by the Austrian Academy of Sciences, the city of Vienna, the European Research Council (ERC) Starting Grant under the European Union’s Horizon 2020 program (ERC-2019-StG-851470), and the Stand-Alone Austrian Science Fund (FWF) grant P34880. Raw sequencing reads are available in the SRA database (PRJNA1159832).

## List of Supplementary Materials

Materials and Methods Figs. S1 to S18

Tables S1 to S5 References (*46–69*)

