## Supplementary files for "A regulatory module driving the recurrent evolution of irreducible molecular complexes"

**The PDF file includes:**

Materials and Methods

Figs. S1 to S18

Tables S1 to S5

References

**Other Supplementary Materials for this manuscript include the following:**

Data S1 to S9

**Materials and methods**

**Maintenance of worm strains**

Nematodes were grown on modified nematode growth medium (NGM) plates with 1% agar/0.7% agarose to prevent *C. tropicalis* burrowing. Plates were seeded with OP50. Experiments were conducted at 25°C. In case of bacterial contamination, worms were bleached using 500 µl of 1:1 mix of 1M NaOH and 5% bleach added to 750 µl of worms in M9, bleaching was stopped by washing embryos twice in 14 ml of M9. Table S3 lists all the strains used in this study, some of which were provided by the CGC, funded by the NIH Office of Research Infrastructure Programs (P40 OD010440).

**Phenotyping and genotyping of crosses**

For crosses, 4–5 L4 hermaphrodites were mated with 40–50 males (1:10 ratio) in a single well of a 12-well plate with modified NGM. After two days 15 L4 F_1_ progeny were transferred in bulk to a fresh plate, and then the next morning singled into individual plates, to ensure synchronized laying of F_2_ generation embryos. After 3 hours of active laying F_1_ hermaphrodites were collected for genotyping by PCR, and at least 10 embryos from each heterozygous hermaphrodite were transferred to individual plates the same day. F_2_ were staged and phenotyped daily until they either died or reached sexual maturity. Embryonic lethality, larval lethality, arrested development, delayed reproduction and sterility were assessed. For maternal and paternal backrosses F_1_s were additionally mated overnight with males or hermaphrodites of the appropriate parental strain. For genotyping by PCR worms were digested with lysis buffer (10 mM Tris-Cl pH8, 50 mM KCl, 2.5 mM MgCl_2_, 200 µg/ml Proteinase K). PCRs were performed using in-house hot-start Taq polymerase (Molecular Biology Service, IMBA). If necessary, PCR products were additionally analyzed by Sanger sequencing. Efficiency of PCR was low for progeny arrested as embryos or at early larval stages. Detailed information on genotyping and phenotyping of crosses can be found in Data S1.

**Generation of *C. tropicalis* transgenic lines**

For CRISPR/Cas gene editing, we adapted previously described protocols (*43*). In brief, 250 ng/µl Cas9 or Cas12a proteins (IDT) were incubated with 200 ng/µl crRNA (IDT) and 333 ng/µl tracrRNA (IDT) at 37°C for 10 min before adding 2.5 ng/µl co-injection marker plasmid (pCFJ90-mScarlet-I, or pCFJ90-mNeonGreen for editing of lines expressing mScarlet-tagged proteins). For homology-directed repair, donor oligos (IDT) or biotinylated and melted PCR products were added at a final concentration of 200 ng/µl or 100 ng/µl, respectively. Following injections into young hermaphrodites, mScarlet or mNeonGreen-positive F_1_ progeny were singled, and their offspring screened by PCR and Sanger sequencing to detect successful editing. To insert single-copy transgenes we adapted a split selection system developed for *C. elegans* (*44*). We introduced two synthetic landing pads (SLPs) with split hygromycin B resistance in the *C. tropicalis* EG6180 Chr. I and Chr. IV. For homology-directed repair we used plasmids carrying an insert of interest followed by the N-terminal portion of the hygromycin resistance gene under ribosomal promoter *Ctr-rps-20p*, at a final concentration of 50–60 ng/µl. On day 3 or 4 post-injection we supplemented plates with 600 µl of 5 mg/ml Hygromycin B. After 7–10 days after poisoning, survivors were singled, propagated and screened by PCR and Sanger sequencing to detect successful editing. All gRNAs and HDR templates as well as plasmids used for SLP injections are listed in Table S4 and Table S5 and are available upon request. Lines carrying genes of interest in both SLPs were obtained by crossing strains carrying single Chr. I and Chr. IV SLP insertions. Double carriers were screened by PCR and Sanger sequencing to ensure homozygosity of both alleles of interest. To obtain homozygous *kss-3.1-3.4* quadruple knock-out, *kss-3.1; kss-3.2; kss-3.4* triple knock-out, and *kss-3.1; kss-3.4* double knock-out lines additional steps were necessary, as standard screening procedure following CRISPR/Cas gene editing only resulted in heterozygous lines, suggesting that homozygosity in this locus is highly detrimental for the worms. We screened a large number of offspring from heterozygous hermaphrodites to obtain low frequency homozygous mutant survivors. All transgenic lines and wild isolates used in this study were bulk phenotyped to test if any abnormal phenotypes were present. L4 hermaphrodites were collected, grown overnight, and the next singled onto new plates for laying for 3 hours. Synchronized embryos were transferred to new plates, 5-10 embryos per plate, and phenotyped until adulthood. Phenotypes like embryonic and larval arrest, developmental delay, and sterility were assessed. At least 60 embryos were screened per strain. Data can be found in Table S1.

**Genetic mapping of *klmt-1/kss-1* and *pzl-1/kss-2***

To facilitate the mapping of the NIC203 Chr. V TA, we have previously generated a Near Isogenic Line (NIL) carrying a NIC203 introgression (Chr. V: 1,260,950–1,735,109 Mb) in an otherwise EG6180 genetic background. The Chr. V NIL parental strain was named QX2343 (*7*). Within this introgression, we identified 176 genes (Data S2). Next, we sought gene pairs in NIC203 that were either (i) absent, mutated or divergent in EG6180 and (ii) in tight genetic linkage. From the handful of examples available in nematodes, genes coding for toxins and antidotes tend to have weak or no homology to conserved nematodes genes. Thus, to identify candidates, we defined *C. tropicalis* “divergent” genes as those showing ≤ 90% sequence identity ≤ 80% sequence coverage to their best reciprocal BLAST hit in *C. elegans* and also ≤ 90% sequence identity ≤ 80% sequence coverage between *C. tropicalis* NIC203 and EG6180. This criteria was matched by one pair of genes: *NIC-ORF015419* and *NIC-ORF015420*. To test whether *ORF015419* coded for the toxin, we generated a putative *ORF015419(-)* null allele using CRISPR/Cas (strain INK303). To test whether *ORF015420* coded for the antidote, we generated a putative *ORF015420(-)* null allele in the *ORF015419(-)* genetic background (strain INK422). Both alleles were tested in genetic crosses. Crosses and background phenotypes are described in detail in Data S1 and Table S1. The susceptible strain, EG6180, carries remnants of *klmt-1/kss-1* TA. *EG-ORF014894* codes for EG-KSS-1, which is predicted to be 387 amino acids long protein. EG-KSS-1 has 15 substitutions and 2 amino acids deletion compared to the NIC203 KSS-1 antidote. *EG-ORF014892* is a pseudogenized version of *klmt-1.* With Mauve (*45*) we annotated an inversion starting in the beginning of exon 3 of *klmt-1* resulting in a null allele (fig. S1C). DNA and protein sequences can be found in the Data S9. To map Chr. II TA, we used QX2341, NIL carrying a NIC203 introgression (Chr. II: 8,077,521–8,836,894) in an otherwise EG6180 background. Within this introgression we identified 243 genes (Data S4). Using the same divergence and linkage criteria previously applied to the Chr. V TA did not reveal any candidate TA genes. Thus, we reasoned that the susceptible TA allele may simply carry deleterious nonsynonymous substitutions A pilot mRNA-seq differential expression analysis of NIC203 and EG6180 transcriptomes identified *ORF006816,* a gene located within this introgression, as one of the most differentially expressed genes between these two parental strains. To test whether *ORF006816* coded for the toxin, we generated a putative *ORF006816(-)* null allele using CRISPR/Cas (strain INK324). To test whether *ORF006815* coded for the antidote, we generated a putative *ORF006815(-)* null allele in the *ORF006816(-)* genetic background (strain INK485). Both alleles were tested in genetic crosses. Crosses and background phenotypes are described in detail in Data S1 and Table S1. The susceptible strain, EG6180, has *pzl-1/kss-2* TA with the following differences compared to the carrier strain. *EG-ORF006586* codes for EG-KSS-2, which has two substitutions compared to KSS-2. *EG-ORF006587* codes for EG-PZL-1 which has eight non-synonymous substitutions compared to PZL-1 toxin. In addition, there is a transposon insertion in intron 6. DNA and protein sequences can be found in the Data S9.

**Genetic mapping of *hyde-1/kss-3***

We run BLASTN (BLAST v.2.8.1+) search using the *kss-1* sequence as a query against NIC203 and EG6180 genomes. While manually annotating top hits, we noticed that three and four paralogues in NIC203 and EG6180 genomes correspondingly, were located in *fars-3* locus, downstream of *zyg-9.2* and upstream of *fars-3*. We named those paralogues *kss-3.1–3.4*. We manually characterized every gene in the region between *zyg-9.2* and *fars-3* that was predicted by Funannotate using BLASTN and BLASTP search for homologs, which resulted in annotating 3 *fars-3* homologs in both genomes, including *EG-ORF007390*. Initially, we hypothesized that KSS-3 might be important for regulation of FARS-3 expression, that’s why we tagged *fars-3* with mScarlet fluorescent tag and knocked-out *kss-3.1–3.4* in FARS-3::mScarlet background. We observed that quadruple knock-out results in a severe yet livable phenotype, which was independent of FARS-3::mScarlet expression. An EMS screen performed in the quadruple knock-out strain, along with subsequent crosses between EMS revertant lines and EG6180 parental line, indicated that *fars-3* paralogue *EG-ORF007390* and *kss-3* form a toxin-antidote pair. The *hyde-1/kss-3* locus differs between NIC203 and EG6180 isolates. Both possess functioning antidotes: EG6180 has two antidotes, *kss-3.1* and *kss-3.4*, while NIC203 has only *kss-3.4. kss-3.4* in NIC203 and EG6180 share 93% of sequence identity. *NIC-ORF007668* codes for *NIC-hyde-1*, which is a divergent variant of *hyde-1* toxin. *NIC-hyde-1* shares 56% of sequence identity with *hyde-1*. Additionally, it has a premature stop codon, leading to a non-functional toxin.

**EMS-mutagenesis screen for *kss-3* suppressors**

EMS forward mutagenesis screen was performed on quadruple *kss-3.1–3.4* knock-out strain in order to find phenotype revertants, reasoning that reversion of phenotype can be caused by mutagenesis of potential KSS-3 binding partner. EMS mutagenesis was performed according to the standard protocol (*46*) with an increased number of L4s to compensate for the high percentage of lethality in *kss-3.1–3.4* (-) strain. In short, 240 L4s were incubated in 50 mM EMS for 4 hours. After extensive washing an equal number of worms were plated on twelve 9 cm plates. Adults were washed off plates the next day, after ensuring there are at least 200 F_1_ embryos per plate laid. F_1_ gravid adults and F_2_ embryos were collected from plates after three days and bleached. F_2_ embryos from each of twelve plates were split into equal proportions and seeded onto four 9 cm plates. Plates were chunked every 4–5 days for three weeks. After that phenotypes were assessed and all plates were rated based on how long it took for a plate to starve. Based on phenotyping results and to ensure optimal sequencing coverage on the chosen Illumina flow cell lane we selected five wild-type looking revertant strains (see Table S1 for phenotyping results) and proceeded with DNA extraction and Illumina library preparation.

**DNA extraction, library preparation and sequencing**

Total DNA was extracted from 2 freshly starved plates using MasterPure Complete DNA and RNA Purification Kit (Biosearch Technologies). Lysis was performed according to the tissue sample lysis protocol from the manufacturer, with the following changes: 4 µl of proteinase K instead of 1 µl, and incubation for 30 minutes instead of 15 minutes. Next part of the protocol, precipitation of total nucleic acids, was followed without any changes. Library preparation was performed using the Illumina DNA Prep kit (Illumina Inc., San Diego, CA, USA) according to the manufacturer's instructions with unique dual indexing. The libraries were checked on a fragment analyzer, pooled at equal ratios by qPCR and sequenced on a NovaSeq X sequencer (Illumina Inc., San Diego, CA, USA) in single-end read mode for 100 cycles.

**Identification of coding variants**

Demultiplexed sequences were aligned to the *C. tropicalis* EG6180 genome using BWA (version 0.7.17) and de-duplicated using Sambamba (Samtools version 1.9). Variants were called using the GATK HaplotypeCaller (version 4.0.1.2) with default settings, and variants unique to each revertant strain were extracted with bcftools (version 1.9). Variant effects were revealed using SnpEff (version 5.2c), using a custom *C. tropicalis* EG6180 database, and moderate-/high-effect homozygous variants were extracted for further analysis.

**RNA extraction and mRNA-seq**

Total RNA was extracted from three full 9-cm plates using TRizol chloroform extraction (*47*). Phase-lock tubes were used to improve phase separation. To remove DNA samples were treated with DNAse I (37C, 30 min), enzyme was inactivated by incubation with 2,5 µM EDTA (65C, 10 min). In the case of staged mRNA-seq, RNA was extracted from three different stages: early embryos, late embryos and L4s. Early embryos were obtained by collecting per sample 20000 synchronized adults before the start of active laying. Adults were additionally gravity sedimented twice to ensure no carryover of laid embryos. Washed adults were bleached and the embryonic fraction was immediately snap-frozen. Late embryos fraction was prepared in the same fashion, but embryos were incubated for 6 hours after bleaching before snap-freezing. An aliquot of each fraction was inspected at the dissecting microscope. Early embryos fraction contains every developmental stage up to the early gastrulation, late embryos – every developmental stage starting from comme stage. For the L4 fraction 2000 L4s were collected per sample. Next, 100 ng of purified RNA was used as input for the TruSeq Stranded mRNA kit (Illumina; catalog no. 20020595), and libraries were prepared according to manufacturer instructions. Libraries were then checked on a fragment analyzer and sequenced on a NovaSeq S4 Lane (XP Workflow) for 300 cycles, paired-end reads of 150 base pairs. Library preparation and sequencing was performed by the Vienna Biocenter NGS facility. All samples were run in biological triplicate or quadruplicate for staged mRNAseq. Transcript quantification and normalization were performed using kallisto (*48*) and DESeq2 (*49*)**.**

**RT-qPCR**

Total RNA from synchronized embryos was extracted using TRizol chloroform extraction as described earlier. RNA concentrations were measured using the Qubit High-Sensitivity RNA fluorescence kit (Thermo). Absence of genomic DNA contamination in representative RNA samples was confirmed by no amplification signal in PCR reactions with gDNA specific primers. cDNA was prepared with SuperScript III reverse transcriptase (Thermo) using random Oligo(dT) primer. Primers for qPCR were validated with standard curves to ensure amplification efficiency and R^2^ value above 0.95. The following primers were used: FW-cdc-42: 5′-CGATTAAATGTGTCGTCGTAGG-3′, and RV-cdc-42: 5′-ACCGATCGTAATCTTCTTGTCC-3’, FW-klmt-1: 5’-CCAACTACTCCAGTGACATCC-3’ and RV-klmt-1: 5’-CCCTCTGAACTCAACAAATCG-3’. *cdc-42* was used as a housekeeping gene. RT–qPCR reactions were prepared with the Luna Universal qPCR and RT–qPCR kit (NEB) and run with an annealing temperature of 58 °C. All samples had three biological and four technical replicates. We used the ∆∆Ct method to calculate relative fold change.

**Single molecule in situ hybridization**

Stellaris FISH probes targeting *klmt-1* and *pgl-1* were designed using the Stellaris RNA FISH Probe Designer (Biosearch Technologies). The probes were labeled with CAL Fluor Red 610 and Quasar 670, respectively (Biosearch Technologies). Embryos were processed according to adapted Raj et al. protocol (*50*), described in more detail by the probe manufacturer available at [www.biosearchtech.com](https://www.biosearchtech.com/support/resources/stellaris-rna-fish/stellaris-protocols). Processed embryos were stained with DAPI (Merck, D9542, 6 ng/ml), mounted with Fluorshield (Sigma-Aldrich, F6182), and imaged at Axio Imager.Z2 (Zeiss) equipped with Hamamatsu Orca Flash 4 camera (pixel size: 6.5 μm) and HBO lamp. For imaging 63×/1.4 Plan-Apochromat Oil DIC objective was used, Z-stack images with 40 slices (step size 0.2 µm) were acquired. Filters used were: DAPI excitation 406/15 nm, emission 457/50 nm, CAL Fluor Red 610 excitation 545/30 nm, emission 610/75 nm, Quasar 670 excitation 620/60 nm, emission 700/70 nm. Quantification was performed using a MATLAB script from the Raj lab available at <http://bitbucket.org/arjunrajlaboratory>. Image of 2-fold stage embryo (Fig 1E) was deconvolved with Huygens Professional version 24.04.0p4 (Scientific Volume Imaging, The Netherlands, http://svi.nl).

**Heat-shock of *C. tropicalis* early embryos**

Heat-shock induction was performed on 5 cm plates with modified NGM. After dissecting gravid adults with insulin syringe needles (29G) in a drop of M9, embryos were transferred to the plates. After drying for approximately 20 minutes plates were transferred to a 37°C incubator for 40 minutes, plates were kept with their lids on top as opposed to standard upside-down plate storage, to ensure even warming up. The number of embryos was manually counted after heat-shock induction, and plates were kept in a 25°C incubator overnight. Next day the number of unhatched embryos was counted, as well as the number of affected L1s, while wild-type worms were at least L2 stage at this moment. In cases of toxins overexpression, the number of adult survivors was assessed on the second day. Percentage of affected worms was calculated using a sum of non-hatched embryos and affected L1s divided by the total number of embryos used for heat-shock for each strain. All experiments were performed in at least triplicates.

**Heat-shock of *C. tropicalis* early embryos for microscopy**

Gravid adults were washed off from three 9 cm plates, and gravity sedimented twice to ensure no carryover of laid embryos. Collected adults were bleached, and after two washes with M9 embryos were plated on non-seeded 5 cm plates. Heat-shock induction was performed not earlier than one hour after harvesting to ensure the majority of embryos developed enough to have zygotic transcription. After heat-shock plates were incubated at 25°C for 5.5–6 hours, embryos were transferred to slides with agarose pads and imaged on Axio Imager.Z2 (Zeiss), 20x/0.8 Plan-Apochromat objective. Filter used for mCherry: excitation 545/30 nm, emission 610/75 nm. All experiments were performed in at least duplicates.

**Immunohistochemistry**

Gravid adults and laid embryos were washed off plates with M9 medium, followed by bleaching to extract embryos and remove bacteria. The embryo suspension was applied to poly-L-lysine-covered slides (Sigma-Aldrich, P8920). To achieve freeze-cracking of the shells, coverslips were applied and slides were immersed into liquid nitrogen. Immediately after removal of the coverslips, slides were fixed in ice-cold methanol (15 min), followed by ice-cold acetone (10 min), and rehydrated in descending ethanol concentrations (95%, 70%, 50%, and 30% ethanol). Fixed embryos were blocked for 1 hour in 4% BSA (VWR Life Science, 422351S) in PBS-T with 1% of Tween20 (Sigma-Aldrich, P1379) at room temperature. After blocking samples were incubated overnight at 4°C with the primary antibody of choice in blocking solution. Following dilutions were used for primary antibodies: anti-FLAG M2 1:3000 (Sigma-Aldrich, F1804), anti-KLMT-1 1:80 (Monoclonal Antibody Facility, Max Perutz Labs). After washing with PBS-T secondary anti-mouse antibody Alexa Fluor 568 (ThermoFisher Scientific, A-11031) was applied for 1 hour at room temperature, the antibody was diluted 1:3000 in the blocking buffer. Samples were washed four times with PBS-T. DAPI was added to the third wash (Merck, D9542, 6 ng/ml). Processed embryos were mounted with ProLong Diamond Antifade Mountant (Invitrogen, P3696) and imaged at Axio Imager.Z2 (Zeiss) with 40x/1.3 Plan-Apochromat Oil objective. The 20x/0.8 Plan-Apochromat objective was used for imaging of large numbers of embryos that were used for fluorescence quantification experiments. Filters used were: DAPI excitation 406/15 nm, emission 457/50 nm, Alexa Fluor 568 excitation 545/30 nm, emission 610/75 nm.

**Time-lapse imaging for KLMT-1::mNeonGreen**

Synchronized gravid adults were bleached as described earlier, and obtained embryos were immediately transferred to an imaging plastic chamber for inverted microscope (Lab-TekNunc Chambered Coverglass, 155411). Celldiscoverer 7 (Zeiss) equipped with Hamamatsu Orca Flash 4.0 (pixel size: 6.5 μm) camera, LED-module 470 nm (10% intensity), filter set 92 HE, and Plan-Apochromat 20x/0.95 objective was used for imaging. Images with a Z-stack of 13.8 µm (11 slices) were taken every 5 minutes for the time course of 10 hours. Stacks where the majority of embryos were in-focus throughout the whole time-lapse were chosen for quantification. Fluorescence intensity quantification was performed as described below. Representative images of the three strains carrying *klmt-1::mNeonGreen* allele (fig. S8) were taken on Axio Imager.Z2 (Zeiss) with the 20x/0.8 Plan-Apochromat objective. Filter used: excitation 480/40 nm, emission 510LP.

**Fluorescence intensity quantification**

16-bit raw images were analyzed in Fiji. The background was subtracted using a “rolling ball” algorithm with 50 pixels of rolling ball radius for the whole dataset. Next, embryos were selected by the freehand tool, and the same selection mask was used to capture background fluorescence intensity for each embryo. To compare fluorescence intensities between strains we used corrected total cell fluorescence (CTCF) parameter (CTCF = integrated density − (area of selected cell × mean fluorescence of background readings)) normalized to the mean fluorescence value for the strain or sample with the lowest fluorescence intensities. Outliers were identified using the ROUT method (Q = 1%) and removed from the final graphs and statistical analysis. Outlier test was performed for all datasets apart from time-lapse imaging for KLMT-1::mNeonGreen. We chose statistical tests based on data distribution analysis and equality of variance tests. *P*-value was adjusted for multiple comparisons if necessary. The half-life of KLMT-1::mNeonGreen was estimated based on the average values obtained from all embryos in each dataset.

**KLMT-1 expression and purification**

pETM14 plasmid with an N-terminally 6xHis-tagged KLMT-1 with a 3C-PreScission cleavage site was transformed into LOBSTR *E. coli* expression strain. This strain was grown in 3 L of LB up to an OD600 of 0.6 – 0.9 at 37°C, then put on 4°C for 30 min and induced with 0.2 mM IPTG for overnight expression at 18°C. The cell pellets were lysed by sonication in denaturing lysis buffer (4 M guanidinium chloride, 50 mM Tris pH8, 300 mM NaCl, 20 mM imidazole, EDTA-free cOmplete protease inhibitor cocktail (Roche) and in-house 1x benzonase (10000x, 0.4 mg/ml) (Molecular Biology Service, IMBA). The lysate was cleared by centrifugation at 31,000 g for 35 min. The supernatant was then loaded onto a 5 ml HisTrap FF (Cytiva). On column refolding was performed by washing with 30 column volumes (CV) of buffer A (50 mM Tris pH8, 300 mM NaCl, 20 mM imidazole) and an additional 5 CV wash step with 5% buffer B (50 mM Tris pH8, 300 mM NaCl, 500 mM imidazole). The protein was eluted in 50% buffer B. The fractions were checked by in-house Coomassie staining of SDS-PAGE gel. For ion exchange fractions were pooled, diluted 1/10 in buffer IEX-A (50 mM Tris pH8, 0.2 mM DTT) and loaded onto a 6 ml ResourceQ (Cytiva) column. After 30 CV wash in IEX-A, the protein was eluted over a 20 CV gradient to a 100% buffer IEX-B (50 mM Tris pH8, 1 M NaCl, 0.2 mM DTT). The fractions after ion exchange were either directly run on a Superdex200 Increase 10/300 GL (Cytiva) in SEC buffer (100 mM NaPO4 pH8, 100 mM NaCl, 5% glycerol, 50 mM arginine, 50 mM glutamate, 0.2 mM DTT) or flash frozen in liquid nitrogen before that. KLMT-1 identity was confirmed by in solution mass spectrometry at 0.1 mg/ml. 20 µl of the solution containing the purified KLMT-1 in 50 mM Tris pH 8, 300 mM NaCl were mixed with 20 µl of 10 M Urea in 200 mM ammonium bicarbonate (ABC). Proteins were reduced with 10 mM DTT for 1 hour at 37°C followed by alkylation with 20 mM Iodoacetamide for 30 minutes at room temperature in the dark and 30 minutes of quenching with another 5 mM DTT. Proteins were digested with 500 ng of lysyl endopeptidase (Lys-C, Fujifilm Wako Pure Chemical Corporation) for 2 hours at 37°C. Subsequently the solution was diluted to 2 M Urea with 100 mM ABC and the proteins were digested with 500 ng trypsin (Trypsin Gold, Promega) at 37°C overnight. The digest was acidified by addition of trifluoroacetic acid (TFA, Pierce) to 1%.

**Monoclonal antibody generation against KLMT-1**

Mouse monoclonal antibody against full-length KLMT-1 (clone 1A3-3E5) was generated at the Max Perutz Laboratories Monoclonal Antibody Facility. Briefly, BALB/c mice were immunized with recombinant, His-tagged KLMT-1. Splenocytes of the best responding mouse according to serum screening by western blot were fused with X63-Ag8.653 mouse myeloma cells, and hybridoma clones were established by HAT selection. 8 days after fusion, hybridoma supernatants were screened by western blot for the presence of KLMT-1 specific antibodies, and antibody-secreting hybridoma single clone 1A3-3A5 was established. The monoclonal antibody was validated using NIC203 (positive control) and EG6180 (negative control) protein lysates.

**Worm protein lysate preparation and western blot**

Two mixed stage plates were used to prepare protein lysate if not stated otherwise. Worms were washed off plates with M9, washed twice to remove bacteria, pelleted and resuspended in ice-cold lysis buffer: 50 mM HEPES pH7.4, 150 mM NaCl, 2 mM MgCl_2_, 0.05% IGEPAL, 10% glycerol, protease inhibitors (Roche, 11836153001), and in-house 1x benzonase (10000x, 0.4 mg/ml) (Molecular Biology Service, IMBA). After snap-freezing samples were lysed by sonication in Bioruptor (UCD-200, Diagenode) 3 cycles of 30/30 sec ON/OFF for 8 min at high energy in an ice-water bath, followed by centrifugation to obtain clean supernatant. Protein concentration was quantified using Bradford assay (Thermo Scientific, 23238), and adjusted to 1.5 mg/ml with lysis buffer. Samples were incubated with an SDS loading buffer for 10 minutes at 95°C and loaded onto NuPAGE Bis-Tris 4–12% gel (Invitrogen). After electrophoresis, samples were transferred to 0.45 µm PVDF membrane (Thermo Scientific, 88518) and blocked with 4% non-fat milk in TBS-T with 1% of Tween20 (Sigma-Aldrich, P1379) for one hour at room temperature. Blocked membranes were incubated with primary antibodies in blocking buffer overnight at 4°C. Following primary antibody dilutions were used: anti-FLAG M2 1:2000 (Sigma-Aldrich, F3165), anti-actin 1:3000 (Abcam, ab13772), anti-tubulin 1:3000 (Sigma-Aldrich, 05-829-AF647) or anti-KLMT-1 1:60 (Monoclonal Antibody Facility, Max Perutz Labs). After incubation, membranes were washed with TBS-T, followed by the incubation with respective HRP-conjugated secondary antibodies: anti-mouse (1:10,000, Invitrogen, G-21040) or anti-rabbit (1:10,000, Jackson Immuno, 111-035-045). anti-alpha-tubulin antibody is conjugated with Alexa Fluor 647, so it does not require a secondary antibody. Substrate detection was performed using ECL reagent (Cytiva, RPN2106) and imaged with ChemiDoc MP (Bio-Rad). Membranes were stripped for one hour before reprobing if necessary (Thermo Scientific, 21059).

**Co-immunoprecipitation followed by mass-spectrometry**

Worm pellets were resuspended in 200 µl cold buffer A (50 mM HEPES pH 7.3, 150 mM NaCl, 10% glycerol, 0.05% NP-40, 1 mM EDTA; freshly supplemented with 2x cOmplete™ EDTA free (Roche) and 1mM TCEP). Worms were lysed using a Bioruptor Plus (Diagenode) with the following settings: 3 repeats of 30/30 sec ON/OFF for 8 min at high energy in an ice-water bath. Lysates were clarified by centrifugation at 18,000 g for 20 min at 4°C. Total protein concentration was quantified and using the Bradford assay. At this stage, an input sample was collected in the SDS buffer for later analysis. Next, 1 mg of clarified protein extract was incubated with 15 µl of settled anti-FLAG M2 magnetic agarose beads (Millipore, M8823), previously equilibrated in buffer A. Incubation was carried out while rotating for 3 hours at 4°C. Subsequently, an unbound sample was collected in SDS buffer for later analysis. Beads were then washed 4 times in 1 ml buffer A followed by 6 washes in buffer B (50 mM HEPES pH 7.3, 150 mM NaCl). 20% of beads were collected in the SDS buffer for later analysis. The remaining 80% were snap-frozen and stored at -20°C until further processing with on-beads digestion for mass spectrometry. The CoIP-experiment was validated by analyzing input, unbound, and IP samples by silver staining as well as western blotting using standard protocols. For on-beads digestion beads were resuspended in 50 µl of 100 mM ABC, supplemented with 400 ng of lysyl endopeptidase (Lys-C, Fujifilm Wako Pure Chemical Corporation) and incubated for 4 hours on a thermo-shaker with 1200 rpm at 37°C. The supernatant was transferred to a fresh tube and reduced with 0.5 mM Tris 2-carboxyethyl phosphine hydrochloride (Sigma) for 30 minutes at 60°C and alkylated in 3 mM methyl methanethiosulfonate (MMTS, Fluka) for 30 min at room temp protected from light. Subsequently, the sample was digested with 400 ng trypsin (Trypsin Gold, Promega) at 37°C overnight. The digest was acidified by addition of trifluoroacetic acid (Pierce) to 1%. A similar aliquot of each sample was analyzed by LC-MS/MS.

**Nano LC-MS/MS**

The nano HPLC system (UltiMate 3000 RSLC nano- or Vanquish Neo UHPLC-System) was coupled to an Orbitrap Exploris 480 mass spectrometer, equipped with FAIMS pro interface and a Nanospray Flex ion source (all parts Thermo Fisher Scientific). Peptides were loaded onto a trap column (PepMap Acclaim C18, 5 mm × 300 μm ID, 5 μm particles, 100 Å pore size, Thermo Fisher Scientific) at a flow rate of 25 μl/min using 0.1% TFA as mobile phase. After loading, the trap column was switched in line with the analytical column (PepMap Acclaim C18, 500 mm × 75 μm ID, 2 μm, 100 Å, Thermo Fisher Scientific). Peptides were eluted using a flow rate of 230 nl/min, starting with the mobile phases 98% A (0.1% formic acid in water) and 2% B (80% acetonitrile, 0.1% formic acid) and linearly increasing to 35% B over the next 60 (for in gel digests) or 120 min. This was followed by a steep gradient to 95% B in 1 min, stayed there for 6 min and ramped down in 2 min to the starting conditions of 98% A and 2% B for equilibration at 30°C. The Orbitrap Exploris 480 mass spectrometer was operated in data-dependent mode, performing a full scan (m/z range 350-1200, resolution 60,000, normalized AGC target 300%) at 3 different compensation voltages (CV -45V, -60V and -75V), followed by MS/MS scans of the most abundant ions for a cycle time of 0.9 seconds for each. MS/MS spectra were acquired using an isolation width of 1.2 m/z, normalized AGC target 200%, minimum intensity set to 25,000 or 50,000 (gel digests), HCD collision energy of 30 %, maximum injection time of 100 ms and resolution of 30,000. Precursor ions selected for fragmentation (include charge state 2-6) were excluded for 10 s (gel digests) or 45 s. The monoisotopic precursor selection (MIPS) mode was set to peptide and the exclude isotopes feature was enabled. Proteins with at least two peptides identified were used for volcano plots.

**KLMT-1 injection into gonads of hermaphrodites**

For each injection, a freshly prepared batch of KLMT-1 was used. Protein was concentrated and buffer was exchanged to injection buffer (50 mM Tris pH8 and NaCl 300 mM) using Vivaspin 500 Centrifugal Concentrator with 10 kDa cut-off. Final KLMT-1 concentration used for injection was 2.7 mg/ml. 0.75 µl (100 ng/µl) of co-injection marker pCFJ90-mSI (*myo-2p::mScarlet-I::unc-54* *5’UTR*) was added to 30 µl protein injection mix to be able to visually distinguish the offspring that received an injection dose. After injection into both arms of the gonads hermaphrodites were kept on a recovery plate for 2 hours, and replated on fresh plates. Next day, the worms were transferred to fresh plates again. The offspring from each plate was screened under stereomicroscope Axio Zoom.V16 (Zeiss), and the phenotypes were counted separately for the offspring with and without co-injection marker. Only offspring that received a co-injection marker were used to assess the proportion of affected offspring. Raw data is available in Table S2.

**Co-expression and purification of KSS-1::Strep–KLMT-1–SKR-1**

The coding sequences were cloned from cDNA into a baculovirus expression vector pGB-Dest (*51*). Sf9 cells were transfected, expressed the proteins at 27°C, and were harvested 4 days after proliferation arrest. Harvested samples were stored at -70°C until purification. 1L of cells were thawed on ice and resuspended in lysis buffer consisting of 1xPBS, 50 mM sodium citrate, 1x benzonase (10000x, 0.4 mg/ml) (Molecular Biology Service, IMBA), EDTA-free cOmplete protease inhibitor cocktail (Roche), 200 µl BioLock (IBA), and 0.5 mM TCEP. The cells were centrifuged for 40 min at 50,000 g at 8°C. For affinity purification the supernatant was loaded onto a 5 ml StrepTrapXT (Cytiva). The column was washed with 20 column volumes (CV) of PBS supplied with 5 mM ATP and 10 mM MgCl_2_. To avoid co-purification of Sf9 chaperones, an additional manual wash was carried out as follows: 1) the flow-through from the StrepTrapXT was retrieved, boiled at 95°C for 5 min, and spun down at 21,000 g for 10 min; 2) the supernatant from this step was diluted to 1 mg/ml with wash buffer (PBS, 5 mM ATP, and 10 mM MgCl_2_) and 3 CV were manually loaded on to the column; 3) column was incubated with this solution on room temperature for 10 min to allow the chaperones to bind to misfolded proteins; and finally the column was reattached to the FPLC. After 15 CV of PBS wash, the complex was eluted with PBS and 50 mM biotin with 0.5 ml/min in the up-flow mode. Individual bands were digested in gel (see below) and their identity was confirmed by mass spectrometry. When examining the fractions from the affinity purification, we noticed that the first ATP/Mg^2+^ wash contained proteins washed off the column. Although present at low concentrations, this protein fraction appeared pure, with the two main bands corresponding KSS-1::Strep and SKR-1. The wash fraction, without carry-over from the flow-through, was dialyzed in two steps: first against 1 L of 20 mM Tris pH7.5 with 150 mM NaCl, and then against 1 L of 20 mM Tris pH7.5 with 50 mM NaCl to remove ATP and NaCl before anion exchange. This sample was then loaded onto a 6 mL ResourceQ column (Cytiva) and eluted using increasing NaCl concentrations. Fractions containing both KSS-1::Strep and SKR-1 were then concentrated and loaded onto a Superdex 200 Increase 10/300 GL column (Cytiva) (Fig. 2E) using 20 mM Tris pH7.5 and 150 mM NaCl as a running buffer. The main peak fractions were pooled again, concentrated, and filtered for SEC-MALLS (Fig. 2E inset) using the same running buffer.

**In gel digest**

Coomassie-stained gel bands were cut to 2-3 mm pieces, transferred to 0.6 ml tubes and incubated with different solutions by shaking for 10 minutes at room temperature followed by removal of the supernatant as follows: gel pieces were washed with 200 µl 100 mM ABC, destained by 2 repeated rounds of shrinking in 200 µl 50% acetonitrile (ACN) in 50mM ABC and reswelling in 200 µl 100 mM ABC. Gel pieces were shrunk with 100 µl ACN before being reduced with 100 µl of 6 mM DTT in 100 mM ABC by incubation at 57°C for 30 min and alkylated with 100 µl of 28 mM MMTS in 100 mM ABC by incubation at RT for 30 min. Wash steps were repeated as described for destining and gel pieces were shortly dried in a speed-vac after the final shrinking step. Gel pieces were soaked in 12.5ng/ul Trypsin in ABC for 5 min at 4°C. Excess solution was removed, ABC was added to cover the pieces and samples were kept overnight at 37°C. The supernatant containing tryptic peptides was transferred to a fresh tube and gel pieces were extracted by addition of 20 µL 5% formic acid and sonication for 10 min in a cooled ultrasonic bath. This step was performed twice. All supernatants were unified. A similar aliquot of each digest was analyzed by LC-MS/MS

**Proteomic data processing**

For peptide identification, the RAW-files were loaded into Proteome Discoverer (version 2.5.0.400, Thermo Scientific). All MS/MS spectra were searched using MSAmanda v2.0.0.19924 (*52*). The peptide mass tolerance was set to ±10 ppm and fragment mass tolerance to ±10 ppm, the maximum number of missed cleavages was set to 2, using tryptic enzymatic specificity without proline restriction. Peptide and protein identification was performed in two steps. For an initial search the RAW-files were searched against a combined database for *C. tropicalis* (Uniprot and Wormbase; 27,687 sequences; 10,300,894 residues) or custom database containing predicted ORFs of EG6180 isolate, based on Funannotate prediction (25,392 sequences; 9,472,164 residues), additionally we searched against Uniprot reference database for *E. coli* (4,360 sequences; 1,354,438 residues), supplemented with common contaminants and sequences of tagged proteins of interest using β-methylthiolation of iodoacetamide derivative respectively on cysteine as a fixed modification. The result was filtered to 1 % FDR on protein level using the Percolator algorithm (*53*) integrated in Proteome Discoverer. A sub-database of proteins identified in this search was generated for further processing. For the second search, the RAW-files were searched against the created sub-database using the same settings as above and considering the following additional variable modifications: oxidation on methionine, deamidation on asparagine and glutamine, phosphorylation on serine, threonine and tyrosine, glutamine to pyro-glutamate conversion at peptide N-terminal glutamine and acetylation on protein N-terminus. The localization of the post-translational modification sites within the peptides was performed with the tool ptmRS, based on the tool phosphoRS (*54*). Identifications were filtered again to 1 % FDR on protein and PSM level, additionally an Amanda score cut-off of at least 150 was applied. Match-between-runs (MBR) was applied for peptides with high confident peak area that were identified by MS/MS spectra in at least one run. Protein areas were computed in IMP-apQuant (*55*) by summing up unique and razor peptides. Resulting protein areas were normalized using iBAQ (*56*) and sum normalization was applied for normalization between samples. Proteins were filtered to be identified by a minimum of 2 PSMs in at least 1 sample and quantified proteins were filtered to contain at least 3 quantified peptide groups. Statistical significance of differentially expressed proteins was determined using limma (*57*).

**Yeast two-hybrid**

To validate the interaction between KSS-1 and SKR-20 we used Matchmaker Gold Two-Hybrid System (Takara, 630489). We followed manufacturer protocol with minor modifications. In brief, coding sequences of both genes were cloned into pGADT7 and pGBKT7 vectors. As a positive control we used interaction between p53 and SV40 large T-antigen, as a negative control – lamin and SV40 large T-antigen interaction. To control for autoactivation we used empty plasmids. pGADT7 and pGBKT7 carrying constructs of interest and control plasmids were co-transformed into Y2HGold strain, 50 ng each, using PEG/lithium acetate method. Transformed cells were plated onto SD/-Leu/-Trp plates, and after 3 days of growth at 30°C individual colonies were inoculated into 3 ml of SD/-Leu/-Trp liquid medium with glucose and grown shaking overnight. Next morning cells were washed twice with 0.8% NaCl, then we measured OD_600_ and adjusted it to OD_600_ = 1 with 0.8% NaCl. Serial dilutions were prepared and spotted in 10 µl aliquots on SD/-Leu/-Trp and SD/–Leu/–Trp/X-α-Gal/AbA plates. Plates were incubated at 30°C for 3 days before final imaging.

**Annotation of KSL proteins and phylogenetic reconstruction**

To annotate genes homologous to the *kss* antidotes across *Caenorhabditis* species, we first constructed a hidden Markov model (HMM) to represent the conserved regions of a small hand-curated set of *C.tropicalis* KSS antidotes and homologous proteins from *C. tropicalis* and *C. wallacei*. Based on preliminary analyses, we prioritized species within the *Elegans* group, focusing on those with chromosome-level genome assemblies (*58*–*61*). These included: *C. elegans* (N2, WBcel235), *C. tropicalis* (EG6180), *C. brenneri* (CFB2252), *C. briggsae* (AF16), *C. doughertyi* (JU1771, nxCaeDoug1.1), *C. inopinata* (NKZ35, Sp34_v7), *C. latens* (PX534, ASM225923v3), *C. remanei* (PX506, CRPX506), *C. sinica* (JU800, nxCaeSini1.1), *C. wallacei* (JU1904, nxCaeWall1), and *C. zanzibari* (JU2190, nxCaeZanz1.1). Also, we included *C. sulstoni* (JU2788, nxCaeSuls) and *C. japonica* (DF5081, nxCaeJapo1.1) as outgroups. We searched genomes with tblastn (BLAST v2.14.0), applying an e-value threshold of 1e-5 to identify potential gene regions. After identifying these regions, we extended them by 2000 nucleotides upstream and downstream, and merged overlapping regions into longer fragments using bedtools (v2.30.0). We predicted the gene structures with genewise (v2.4.1), using these fragments and the HMM. We named each gene based on its chromosomal location in the format *ctr-ksl-[number]* (e.g., *ctr-ksl-29*). Next, we generated a multiple sequence alignment from the predicted genes using mafft-linsi (v7.427), optimizing for sequences with large insertions and deletions. To maintain alignment consistency, we replaced stop codons with the placeholder amino acid 'X'. We then generated nucleotide codon alignments informed by the protein alignment using revtrans (v1.4) with the -readthroughstop option. We filtered the resulting nucleotide multiple sequence alignment to retain only nucleotide positions covered by at least 10% or 80% of the sequences, and we replaced poorly covered positions with gaps. Finally, we constructed a maximum-likelihood phylogenetic tree using IQ-TREE (v2.3.6) with default parameters, allowing us to infer the evolutionary relationships among the *kss/ksl* gene family. An updated HMM including all 560 proteins identified in the first annotation was used to search for new KSL genes; however, only seven additional genes were identified (not included), suggesting that the original search was largely exhaustive. The KSL tree in Figure 5A was generated using a subset of hand-curated proteins that were most closely related to the KSS antidotes. A representative KSL from C. *doughertyi* (KSL-30) was used as an outgroup. Proteins were aligned using MAFFT v7 with default settings and the phylogenetic tree was constructed using IQ-TREE (v2.3.6).

**Phylogenetic reconstruction of *fars-3* and its toxic paralogs**

We retrieved and manually curated orthologs of *fars-3* from the Wormbase ParaSite WBPS19 (WS291) database (*62*, *63*). To minimize artifacts in the multiple sequence alignment, we excluded regions of the toxin sequences that were predicted to be intrinsically disordered based on IUPred2A (*64*), had low pLDDT scores based on AlphaFold2 predictions, and showed poor sequence identity when compared to *C. tropicalis fars-3*. If a region was predicted to be disordered but had significant homology, it was kept for further analysis. Based on these criteria, we retained KLMT-1 [AA:49-404] and HYDE-1 [AA:49-567]. For PZL-1, a chimera consisting of three distinct gene regions—*mec-15*, *zyg-9.2*, and *fars-3*—we only included the segment homologous to *fars-3*, identified through sequence conservation and structural homology predicted by AlphaFold2. This led to the inclusion of PZL-1 [AA:642-788]. All DNA sequences were aligned using MAFFT v7 (*65*) with default settings, and the resulting phylogenetic tree was constructed using IQ-TREE (v2.3.6) (*66*).

**De novo protein structure prediction and analysis**

De novo protein structure prediction and analyses We predicted protein structures using AlphaFold2 (*18*). AlphaFold2 and Alphafold2-multimer were run using the ColabFold (v1.4.0) notebook implementation in Google Colaboratory (*67*). For multiple sequence alignment, we selected the MMseqs2 option. Models were ranked based on pLDDT, and only the best ranked out of five models was selected for further analysis. We searched for structurally homologous proteins in PDB using the DALI (*19*) or Foldseek servers (*68*). Graphics were generated using PyMol (The PyMOL Molecular Graphics System v2.5, Schrödinger) Protein alignment statistics for the selected pairs were generated using the super algorithm for protein pairs with high sequence identity and cealign for protein pairs with low sequence identity. We estimated and plotted the evolutionary conservation of *C. tropicalis* KSL residues using the ConSurf server (*69*).

**Supplementary Figures**

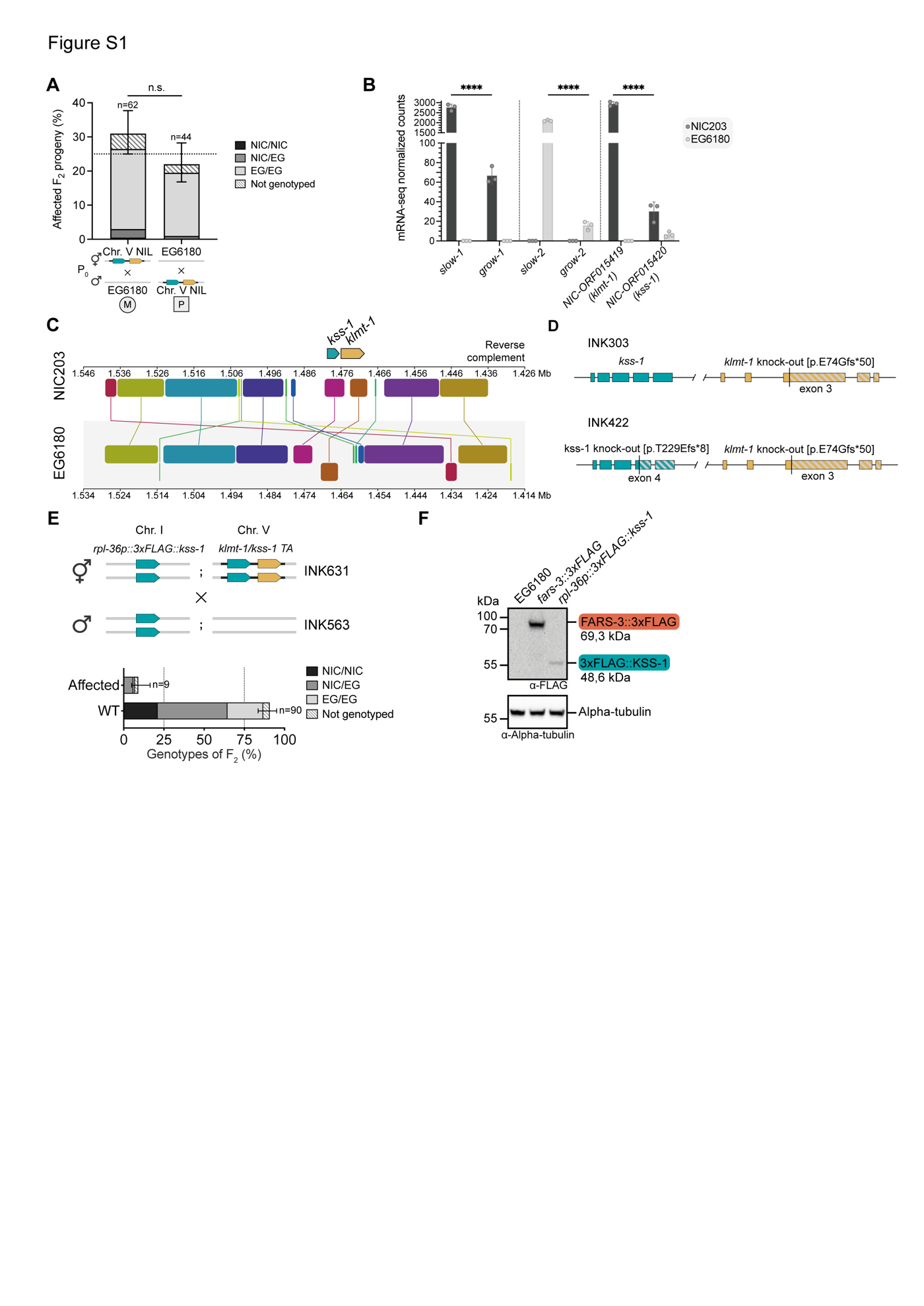

**Figure S1. Genetic mapping of the *klmt-1/kss-1* toxin-antidote element. (A)** Genotype of the F_2_ progeny from a cross between the Chr. V NIL and the EG6180 parental line**.** Affected individuals are homozygous for the EG6180 haplotype (EG/EG). The TA is equally active when inherited either maternally (M) or paternally (P) (*P* = 0.0538, Fisher’s exact test). Error bars indicate 95% confidence intervals calculated with the hybrid Wilson/Brown method. **(B)** Quantification of mRNA expression levels for two previously described *C. tropicalis* TAs, *slow-1/grow-1* and *slow-2/grow-2*, as well as a newly identified TA, *klmt-1/kss-1*. Samples correspond to mRNA-seq data from NIC203 and EG6180 gravid young adults in biological triplicates. Toxin coding genes have higher expression levels compared to antidote-coding genes (two-sided unpaired t-test; *P* < 0.0001 in all cases). Error bars indicate mean with standard deviation. EG6180 is shown in gray, NIC203 in dark gray. **(C)** Similarity profile of NIC203 and EG6180 *klmt-1/kss-1* locus. Blocks located below the center line indicate inversed regions. NIC203 was used as a reference for Mauve alignment, alignment parameters – seed 15, LCB weight 388. **(D)** Schematic of *C. tropicalis klmt-1(-)* and *klmt-1(-) kss-1(-)* mutant alleles generated using CRISPR/Cas. **(E)** Overexpression of 3xFLAG::KSS-1 as a single copy transgene (INK563) in susceptible strain EG6180 rescues the F_2_ lethality associated with the *klmt-1/kss-1* TA. Construct is expressed under a constitutive promoter (*Ctr*-*rpl-36p*). The transgene is integrated in a synthetic landing pad on Chr. I of EG6180. Both parental lines, INK631 (Chr. V NIL background) and INK563, are homozygous carriers for the transgene. Error bars indicate 95% confidence intervals calculated with the hybrid Wilson/Brown method. **(F)** Western blot confirming the expression of 3xFLAG::KSS-1 from the transgenic line used in rescue experiments. Line with endogenously tagged FARS-3::3xFLAG (INK505) was used as a positive control. Negative control is the EG6180 parental strain. Alpha-tubulin serves as a loading control.

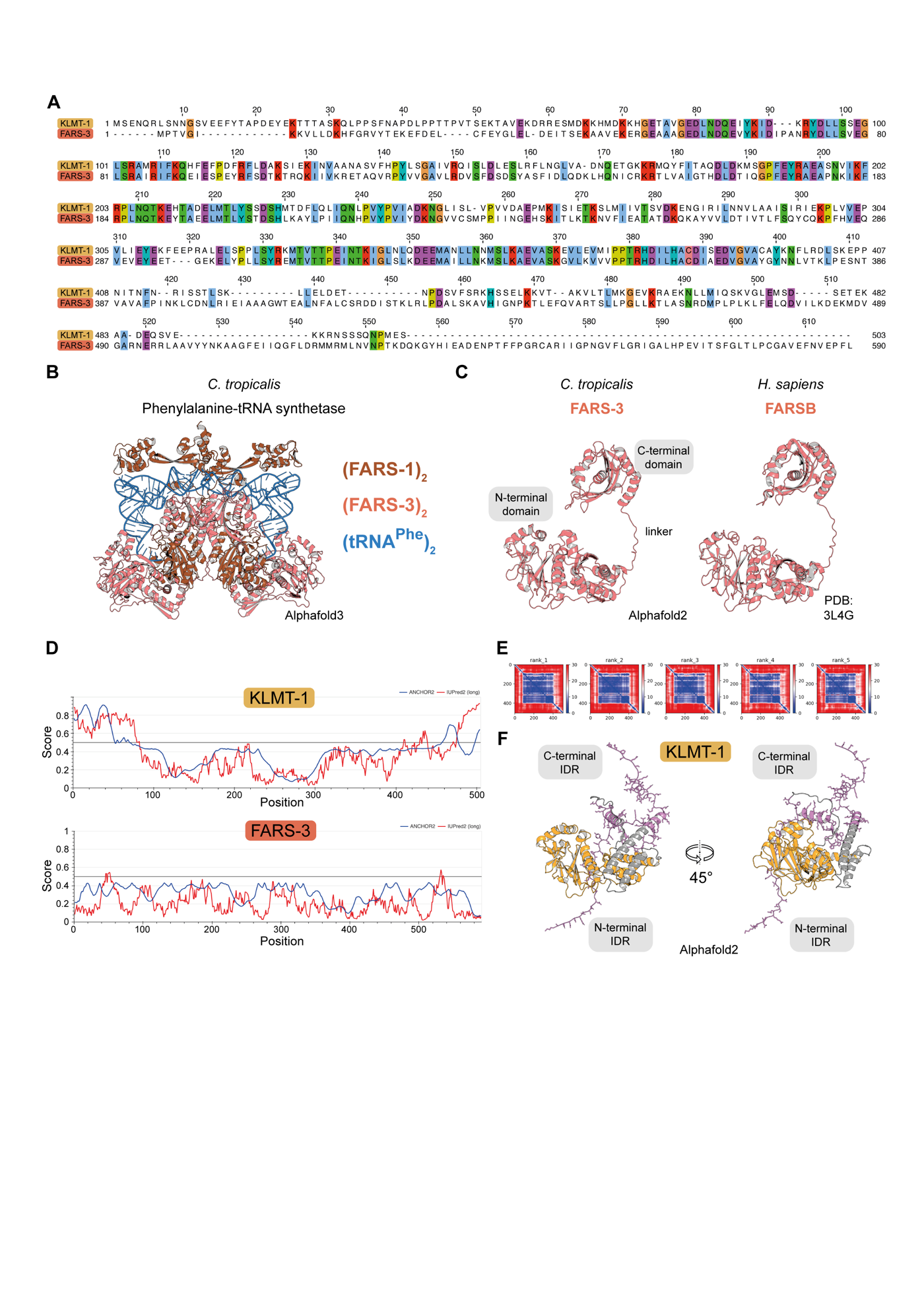

**Figure S2. The *klmt-1* toxin arose from the gene *fars-3* through gene duplication. (A)** Protein alignment of KLMT-1 and *C. tropicalis* FARS-3. Colors correspond to the Clustal X Colour Scheme. **(B)** Alphafold3 model of the *C. tropicalis* Phenylalanine-tRNA synthetase based on the known stoichiometry of the complex in bacteria and eukaryotes. **(C)** Predicted structure of *C. tropicalis* FARS-3 (left) compared to its human ortholog FARSB (PDB: 3L4G) (right). FARS-3 has two main domains: N-terminal (AA:1-376) and C-terminal (AA:390-590), which are connected by a short linker. **(D)** Prediction of Intrinsically Disordered Regions (IDR) for *C. tropicalis* KLMT-1 (top) and *C. tropicalis* FARS-3 (bottom) using IUPred2A (red line). Additionally, disordered binding regions were predicted using ANCHOR2 (blue line). **(E)** Predicted Aligned Error (PAE) of the top five KLMT-1 models generated by Alphafold2. Both N-terminal and C-terminal regions show a high PAE, which is consistent with the predictions that these regions are intrinsically disordered. **(F)** The high-confidence region, which is structurally homologous to the N-terminal domain of FARS-3, is shown in orange. The predicted intrinsically disordered regions are highlighted in violet with side chains emphasized: N-terminal (AA:1-69) and C-terminal (AA:460-503). Regions depicted in gray have low confidence (high PAE) according to AlphaFold2 but are not predicted to be disordered by IUPred2A.

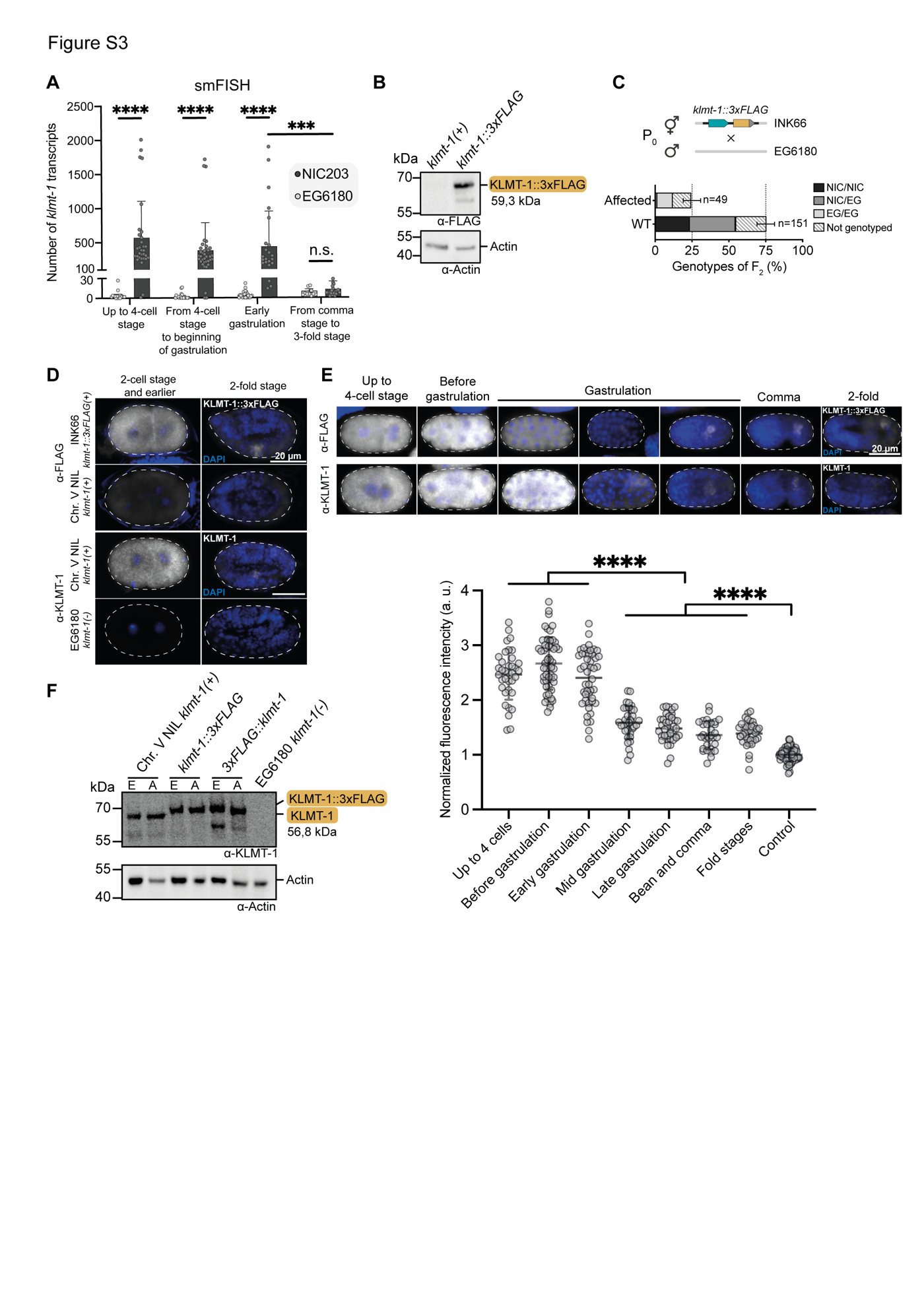

**Figure S3. Expression pattern of *klmt-1* in the adult gonad and during embryogenesis. (A)** Quantification of *klmt-1* transcripts using single molecule fluorescence in situ hybridization (smFISH) throughout embryonic development. EG6180 serves as a negative control. Difference between group means was analyzed using Kruskal-Wallis test (*H*(8) = 128.4, *P* < 0.0001), followed by Dunn's post hoc test (n.s. – not significant, *P*_adj_ > 0.9999; *P*_adj_ for other comparisons < 0.0007). 4-cell stage embryos were included in the “up to 4-cell stage” group. At least 11 embryos were analyzed per group. **(B)** The endogenous *klmt-1* locus was tagged with a C-terminal 3xFLAG using CRISPR/Cas (INK66, Chr. V NIL background). A band of the expected molecular weight (59.3 kDa) was confirmed via western blot. The parental WT Chr. V NIL parental line serves as a negative control. Actin serves as a loading control. **(C)** Genetic cross between the *klmt-1::3xflag* strain (INK66) and the WT EG6180 parental line. In agreement with an active *klmt-1/kss-1*, ~25% of their F_2_ progeny was affected and those individuals affected were homozygous carriers for the susceptible EG6180 allele, indicating that the the tag does not interfere with the activity of the toxin. Error bars indicate 95% confidence intervals calculated with the hybrid Wilson/Brown method. **(D)** Representative images of immunofluorescent staining of early and late embryos expressing KLMT-1::3xFLAG (top panel) and KLMT-1 (bottom panel), stained with α-FLAG or α-KLMT-1 monoclonal antibodies. Chr. V NIL and EG6180 serve as negative control for FLAG and KLMT-1 staining, respectively. **(E)** Representative embryos of different stages that were used for KLMT-1::3xFLAG quantification (top panel). KLMT-1 staining shows a similar expression pattern. Quantification of KLMT-1::3xFLAG throughout embryonic development (bottom panel). Embryos of all early developmental stages have higher KLMT-1::3xFLAG expression (*P_adj_* < 0.0001) compared to mid gastrulation and later developmental stages. EG6180 mixed staged embryos were used as a negative control. At least 30 embryos were quantified per group. Difference between group means was analyzed using Brown-Forsythe ANOVA test (*F**(7, 222.6) = 167.9, *P* < 0.0001), followed by Dunnett T3 post hoc test (for all stages vs control *P_adj_* < 0.0001). Error bars indicate mean with standard deviation. **(F)** KLMT-1 monoclonal antibody recognizes KLMT-1 as well as 3xFLAG::KLMT-1 and KLMT-1::3xFLAG (lines INK63 and INK66 respectively). Lysates were extracted from mixed embryos (E) or adult worms (A). EG6180 serves as a negative control. Actin serves as a loading control.

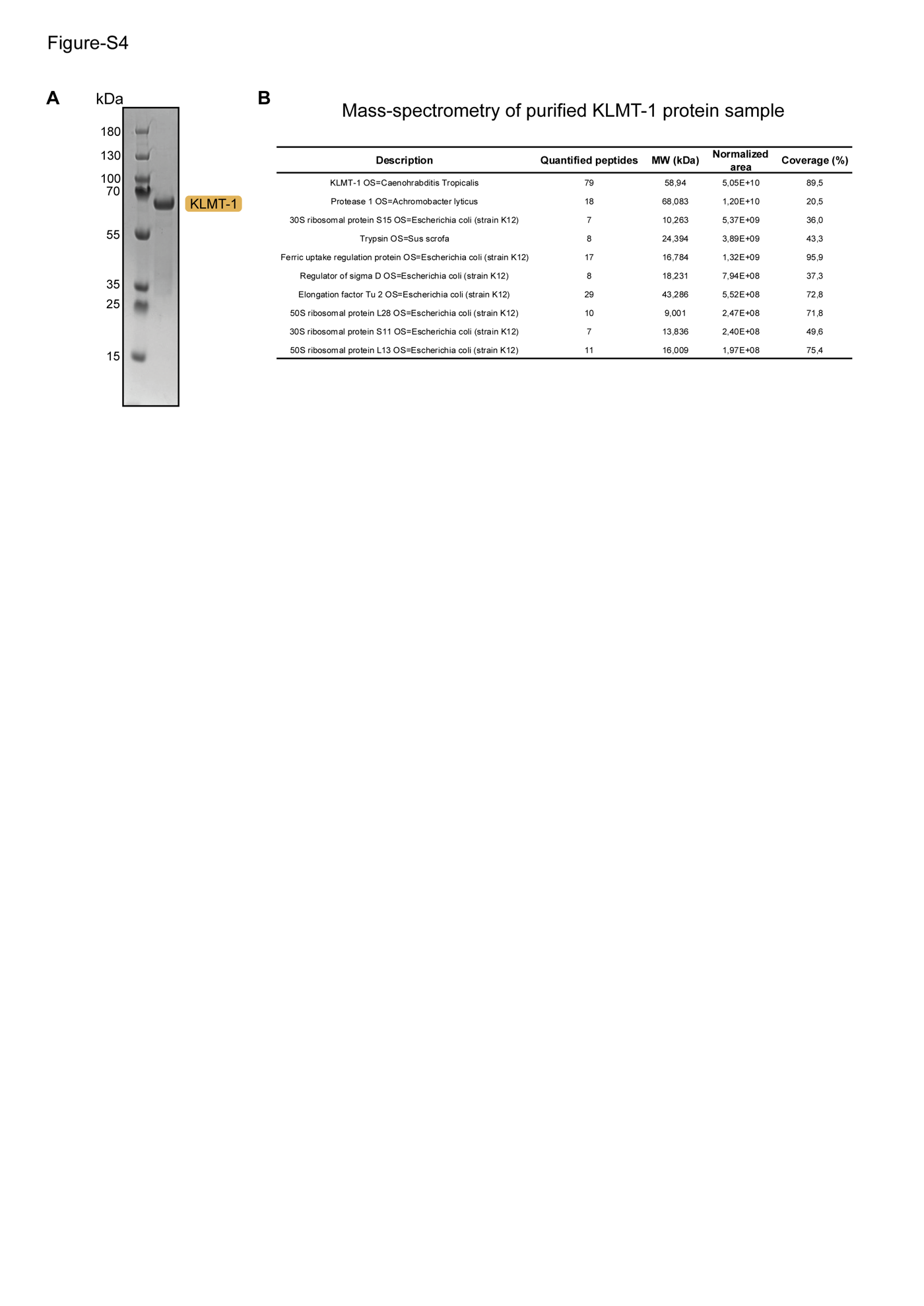

**Figure S4. Expression and purification of KLMT-1 protein from bacterial cells. (A)** KLMT-1 sample after size exclusion chromatography. Protein was additionally concentrated to 2,7 mg/ml for the injection. **(B)** Confirmation that the purified protein is KLMT-1 by mass spectrometry. Ten proteins with the highest normalized area are listed in the table. This purified KLMT-1 protein was injected into the gonad of hermaphrodites to test its *in vivo* activity (Fig. 1F).

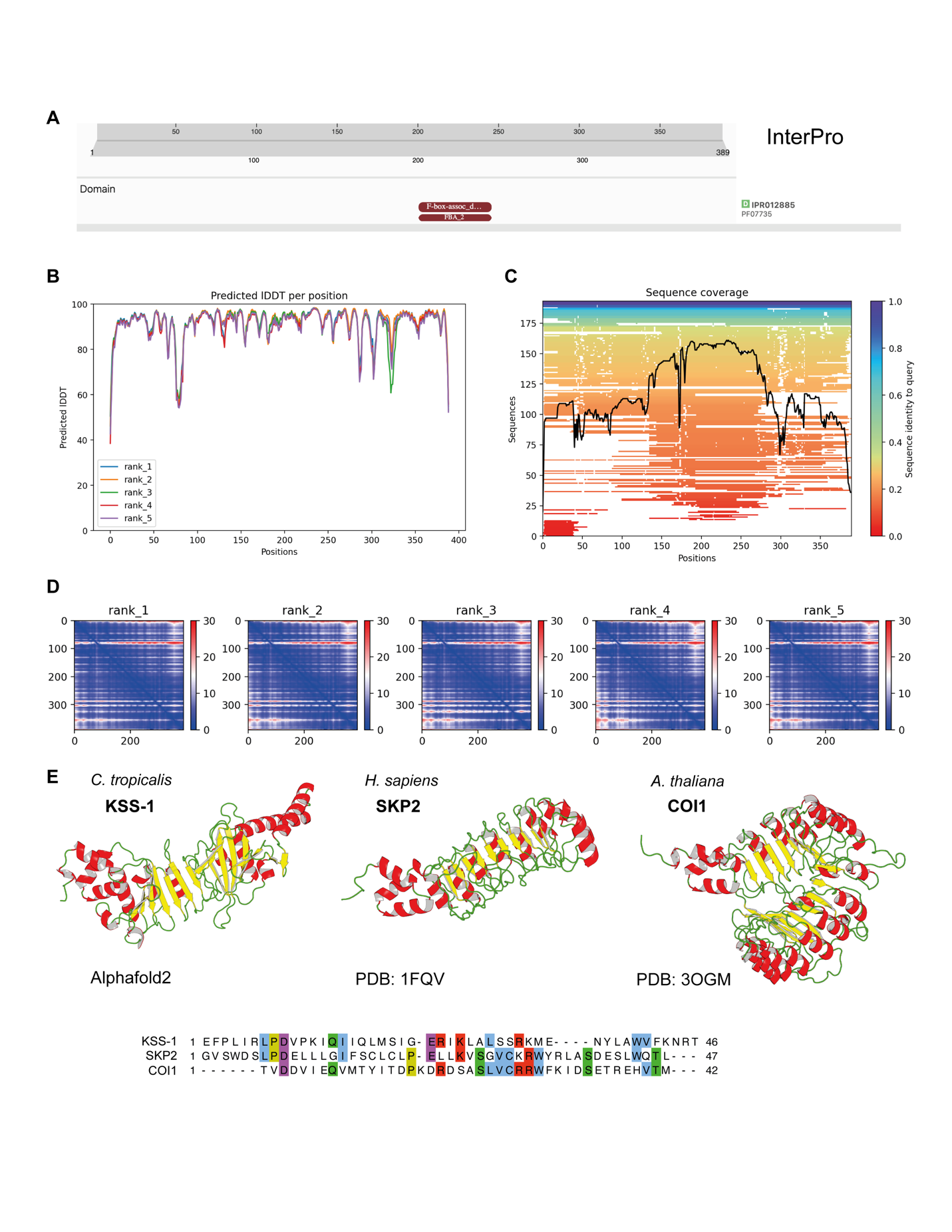

**Figure S5. Alphafold2 reveals that KSS-1 is structurally homologous to F-box proteins. (A)** Prediction of KSS-1 protein domains using InterProScan. An F-box associated domain, type 2 (IPR012885), was found in the middle of the protein but not the N-terminal F-box domain itself. **(B)** Alphafold2 KSS-1 model per-residue confidence metric pLDDT (Local Distance Difference Test). **(C)** Coverage of the multiple sequence alignment. **(D)** Predicted aligned error (confidence in the domain packing and large-scale topology of the protein) for the best five KSS-1 models. **(E)** Comparison of the predicted structure of KSS-1 with the two most similar experimentally solved structures, as identified by the DALI server: human F-box protein SKP2 (PDB: 1FQV) and Arabidopsis F-box protein COI1 (PDB: 3OGM) (top). Alpha-helices are shown in red and beta strands in yellow. Alignment of F-box domains (bottom). Colors correspond to the Clustal X Colour Scheme.

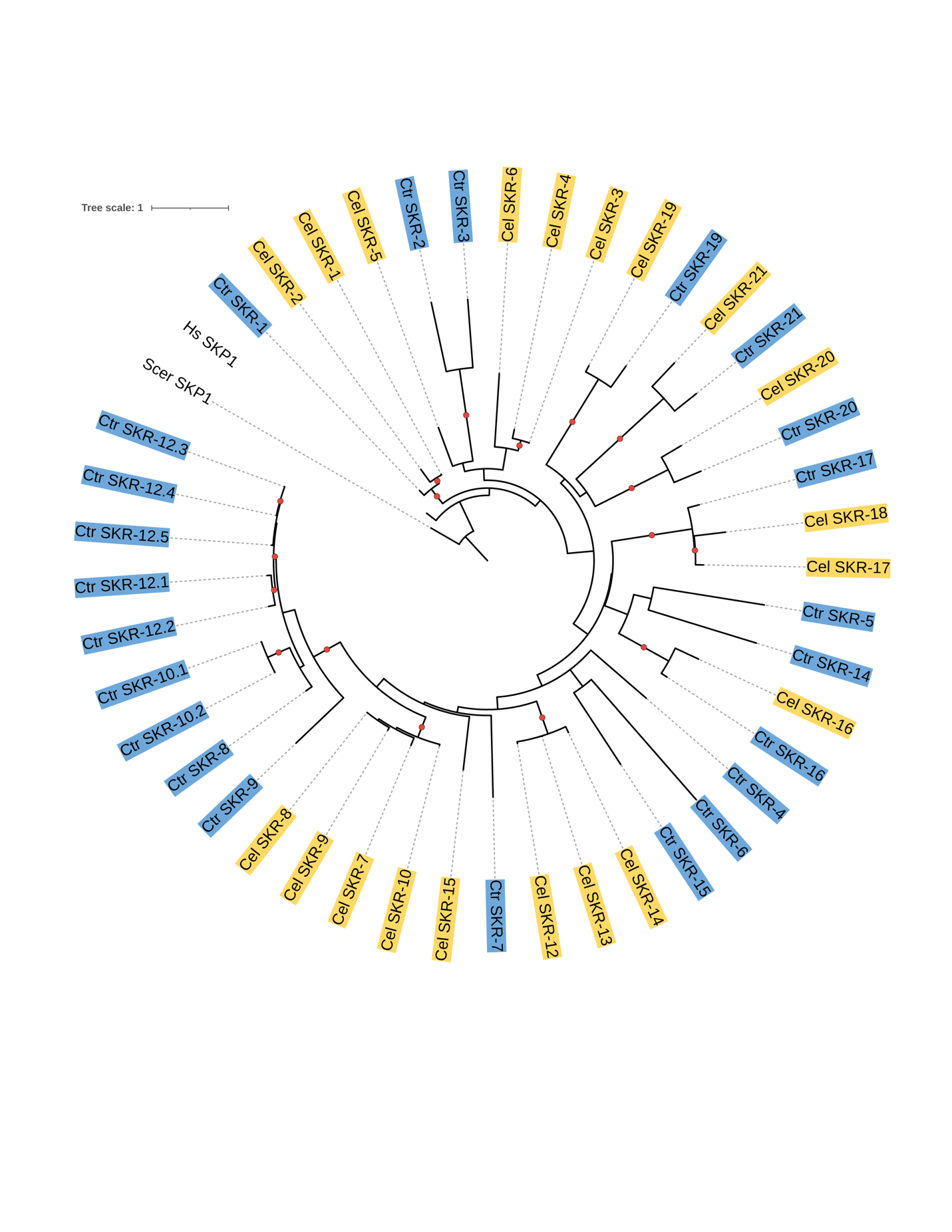

**Figure S6. Phylogeny of *C. tropicalis* SKR proteins.** Phylogenetic analysis of SKR proteins from *C. elegans* (yellow) and *C. tropicalis* (blue). Human SKP1 and *Saccharomyces cerevisiae* SKP1 are included as outgroups. Red circles indicate bootstrap values greater than 95%.

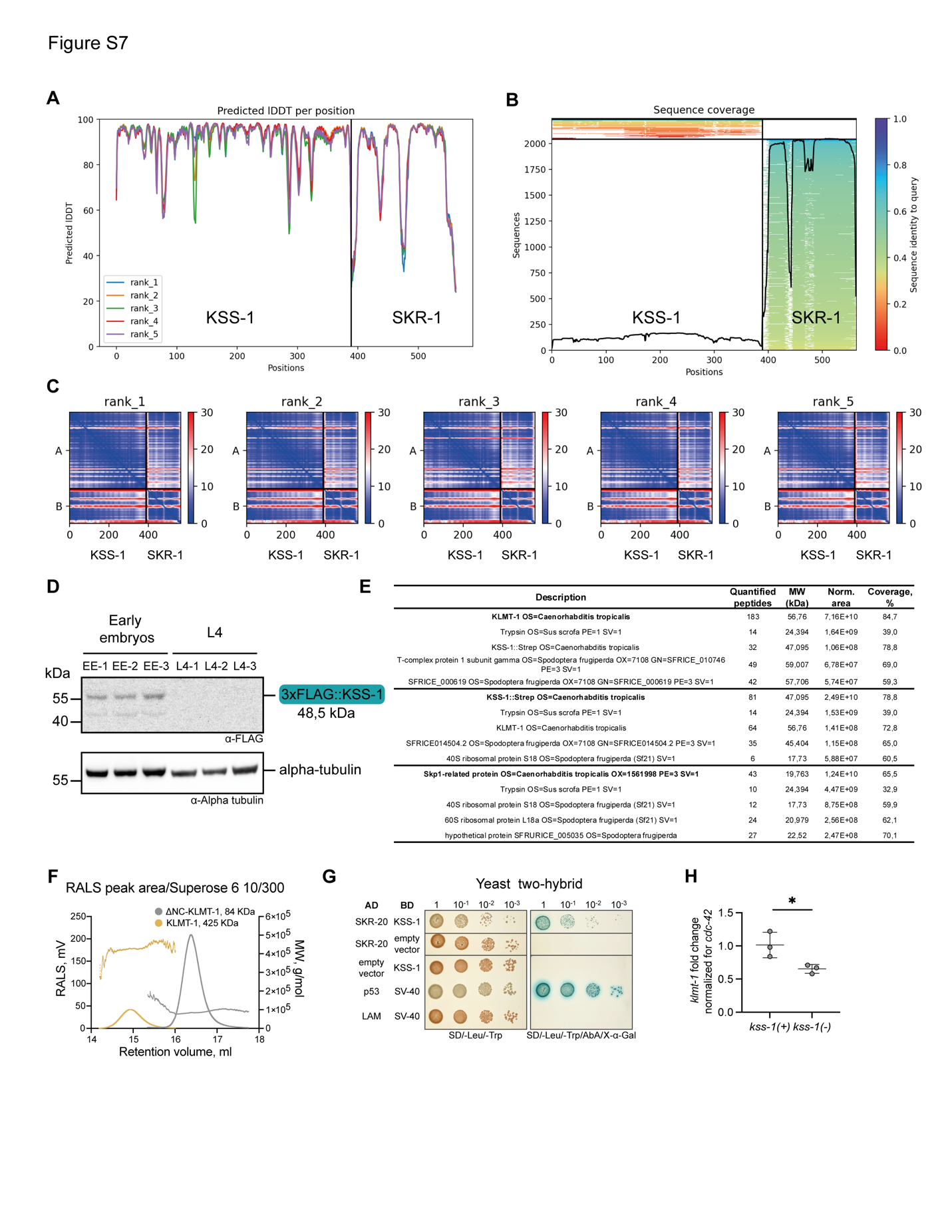

**Figure S7. Characterization of KSS-1 and SKR-1 protein binding. (A)** AlphaFold2-multimer model depicting the binding interaction between KSS-1 and SKR-1, with per-residue confidence indicated by the pLDDT (Local Distance Difference Test) metric. **(B)** Coverage of the multiple sequence alignment for the KSS-1 and SKR-1 interaction. **(C)** Predicted aligned error (confidence in the domain packing and large-scale topology of the protein) for the top five KSS-1–SKR-1 models. **(D)** Western blot analysis of endogenous KSS-1 protein expression. The *kss-1* locus was tagged with a N-terminal 3xFLAG (INK444). KSS-1 protein is detectable in early embryos (embryos collected from gravid adults allowed to lay for 2 hours and then bleached). No KSS-1 protein was observed in synchronized L4 hermaphrodites. This pattern indicates zygotic expression of KSS-1 post-fertilization. Alpha-tubulin serves as a loading control. Samples were analyzed in biological triplicates. **(E)** Bands corresponding to KLMT-1, KSS-1::Strep and SKR-1 (shown in Fig. 2D) were analyzed using liquid chromatography mass spectrometry. Five proteins with the highest normalized area for each band are listed in the table. **(F)** Static Light Scattering (SLS) determination of the molecular weight of full-length KLMT-1 and a truncated version, ΔNC-KLMT-1, which lacks both predicted N- and C-terminal intrinsically disordered regions. Both proteins were expressed in *E. coli*. Full-length KLMT-1 exhibits an apparent molecular weight of ~425 kDa and is prone to aggregation, whereas ΔNC-KLMT-1 does not aggregate and has an estimated molecular weight of 84 kDa, suggesting it forms a homodimer. Aggregation of KLMT-1 prevented determination of the KLMT-1–KSS-1 complex stoichiometry and structure. **(G)** Yeast two-hybrid assay demonstrating the direct interaction between Ctr-SKR-20 and KSS-1. p53 and SV-40 were used as a positive control, while LAM and SV-40 served as a negative control. **(H)** RT-qPCR quantification of *klmt-1* mRNA abundance from *kss-1(+)* and *kss-1(-)* lines (INK169 and INK206, respectively) normalized to *kss-1(+)* line (two-sided unpaired *t*-test; *P* = 0.0374). Error bars indicate mean with standard deviation.

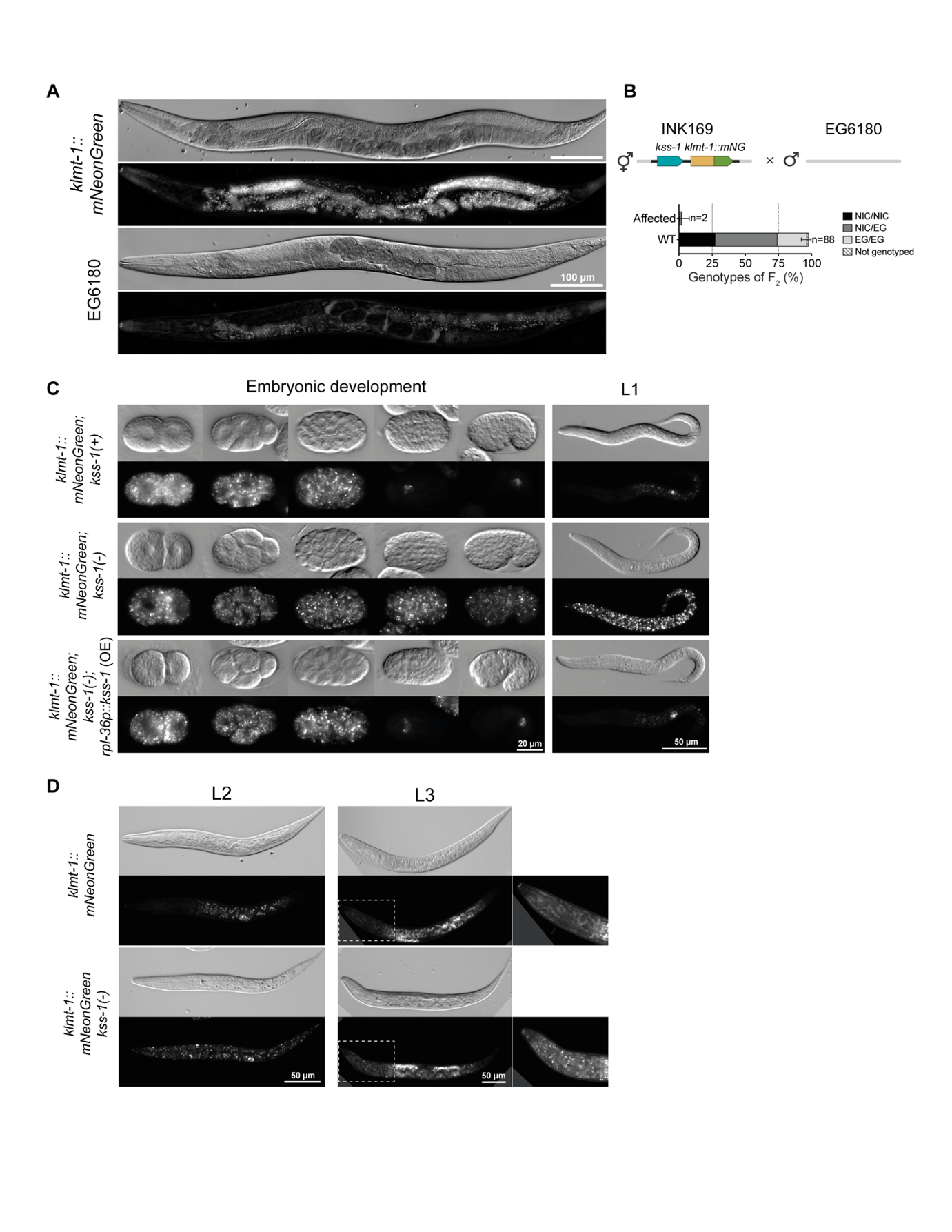

**Figure S8. Characterization of the KLMT-1::mNeonGreen reporter line. (A)** Adult hermaphrodites expressing *klmt-1::mNeonGreen* (INK169, Chr. V NIL background), and corresponding differential interference contrast (DIC) images. KLMT-1::mNG is present in the gonads of adult hermaphrodites, as well as in unfertilized eggs and embryos. The signal observed in the gut corresponds to auto-fluorescence, which is also detectable in the EG6180 strain (negative control). **(B)** Genetic cross between *klmt-1::mNG* strain (INK169) and EG6180 revealed that the *klmt-1::mNG* allele is not toxic, as evidenced by the recovery of healthy homozygous EG/EG individuals in the F_2_ generation. Error bars indicate 95% confidence intervals calculated with the hybrid Wilson/Brown method. mNG – mNeonGreen. **(C)** Representative images of KLMT-1::mNG fluorescence signal through embryonic development (2-cell stage to comma stage) and at the L1 larval stage. In the presence of a wild-type *kss-1(+)* allele, KLMT-1::mNG levels sharply decline after early gastrulation stage (top). In late embryos and larvae, KLMT-1::mNG signal is only detectable in two cells, corresponding to the Z2 and Z3 germline precursors. This signal in the germline likely does not originate from the maternal deposition of KLMT-1 but from its zygotic expression. The strain carrying *kss-1(-)* null allele in the *klmt-1::mNG* background (INK206) was healthy and viable, but showed high levels of KLMT-1::mNG even at late embryonic stages and at L1 stage (middle). To show the specificity of this mutant phenotype, we introduced a single copy transgene overexpressing KSS-1, *rpl-36p::3xFLAG::kss-1* (SLP Chr. I)*,* into the *kss-1(-)* *klmt-1::mNG* background (INK785) and showed that this transgene could rescue the mutant phenotype, leading to efficient clearance of KLMT-1::mNG. Contrast is enhanced for the L1 stage images. OE - overexpression. **(D)** In the absence of *kss-1*, maternal KLMT-1::mNG is not effectively degraded and can be detected in the soma of L2 and L3 larval stages. Zoom-in region shows KLMT-1::mNG in the head of L2 and L3 larvae (bottom). Contrast is enhanced for zoom-in regions. Larvae carrying a *kss-1(+)* allele do not have KLMT-1::mNG in somatic tissues (top).

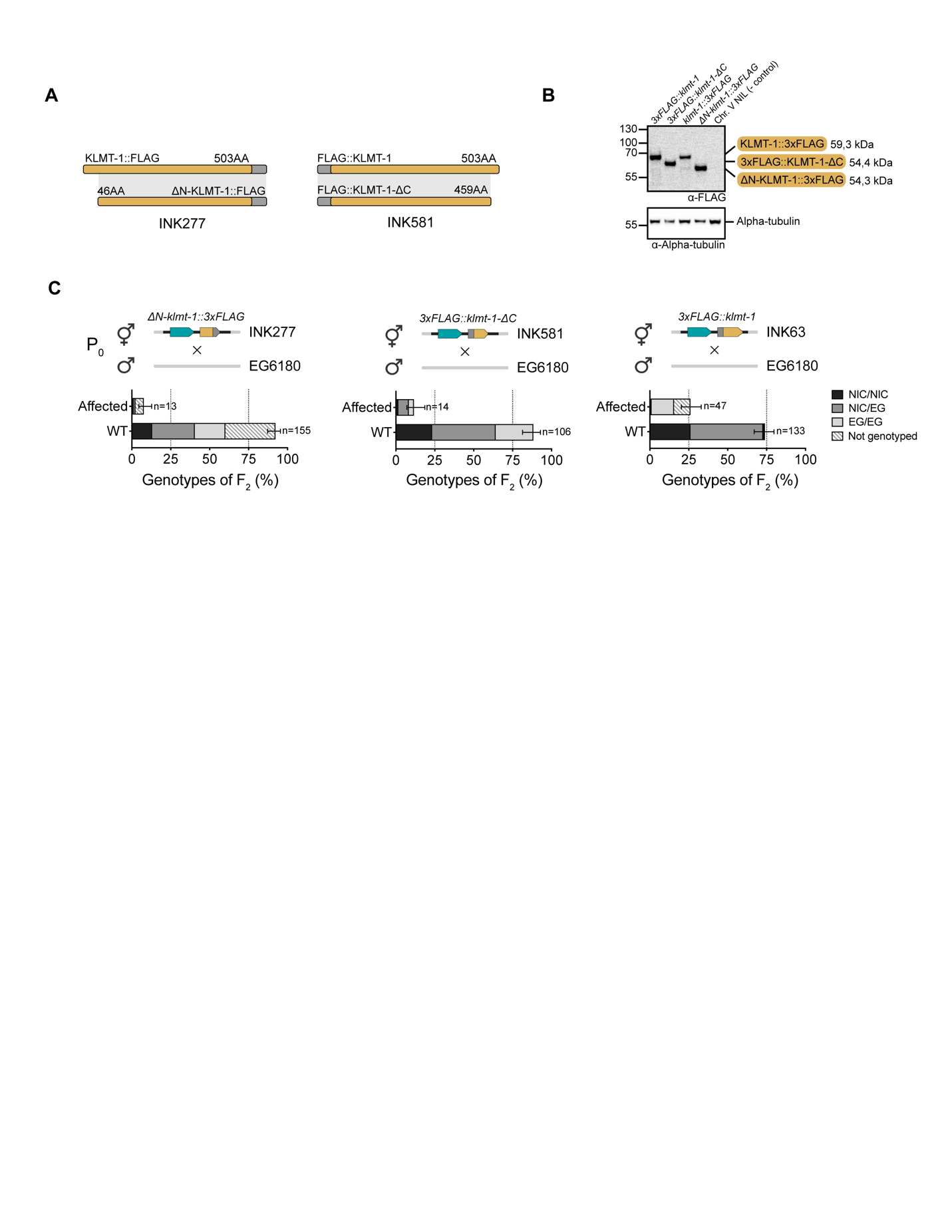

**Figure S9. Characterization of *klmt-1* N-terminal and C-terminal deletion alleles. (A)** Schematic of KLMT-1 protein variants resulting from N-terminal and C-terminal deletion alleles at the endogenous *klmt-1* locus. The N-terminal deletion carries a C-terminal 3xFLAG tag and the C-terminal deletion carries a N-terminal 3xFLAG. N- and C-terminal amino acids are labeled. N-terminal truncation was chosen based on ANCHOR2 prediction, and is 24 amino acids shorter than N-terminal IDR sequence fused to mCherry described in Fig 2H-I. **(B)** Western blot analysis of endogenous KLMT-1 N-terminal and C-terminal deletion alleles. Expression levels of N- and C- terminal deletion alleles are unchanged compared to full KLMT-1::3xFLAG. Chr. V NIL is a negative control. Alpha-tubulin serves as a loading control. **(C)** Genetic crosses to susceptible strain, EG6180, to test the activity of the KLMT-1 N-terminal (INK277, Chr. V NIL background) and C-terminal deletion strains (INK581, Chr. V NIL background). The percentage of affected F_2_ progeny and genotype of surviving progeny indicates that both terminal regions are necessary for toxicity. As a positive control for testing *klmt-1* toxicity in crosses, we used the full length 3xFLAG::KLMT-1 strain (INK63, Chr. V NIL background). Error bars indicate 95% confidence intervals calculated with the hybrid Wilson/Brown method.

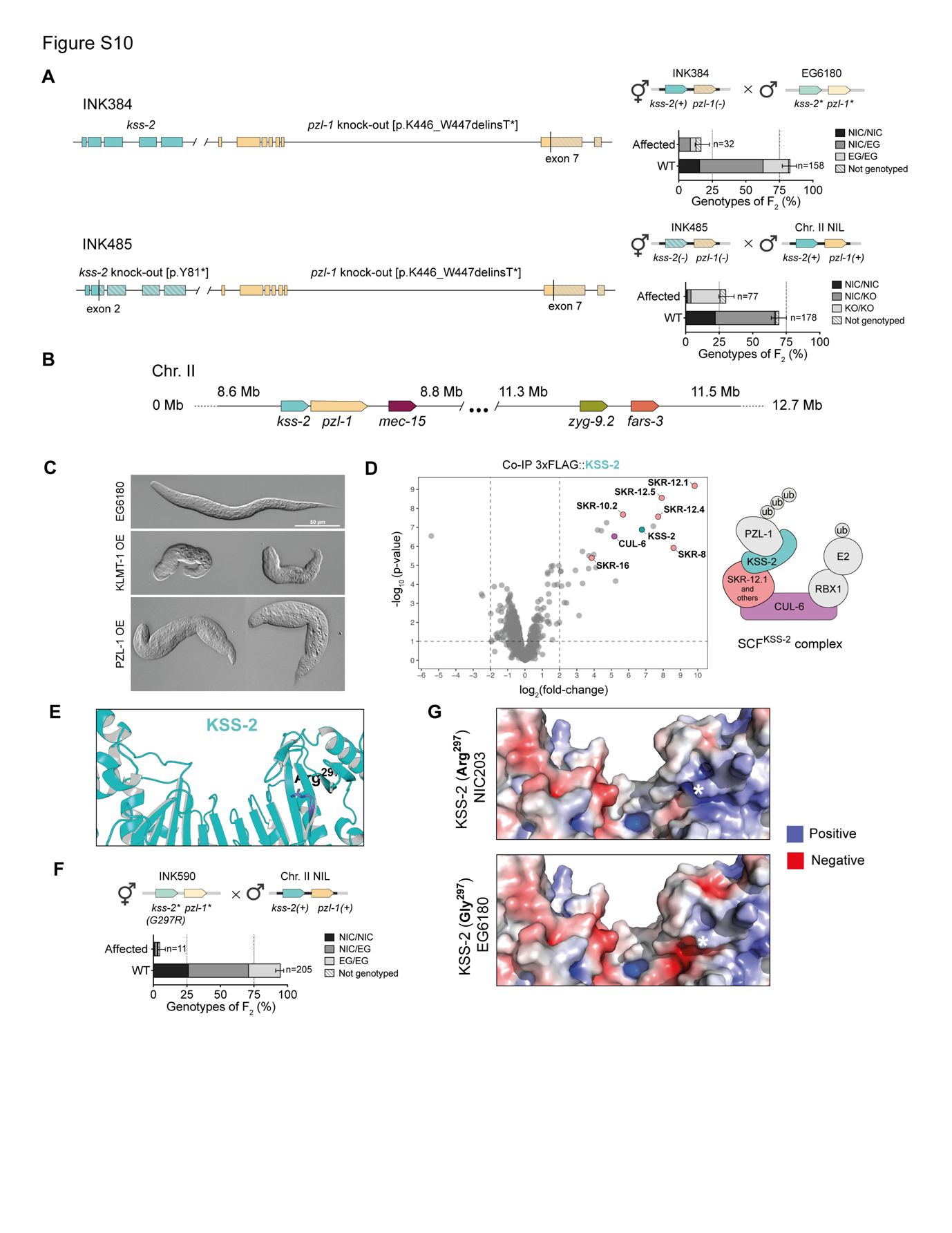

**Figure S10. The PZL-1 toxin is a chimeric gene and its antidote, KSS-2, an F-box protein. (A)** Schematic of *C. tropicalis pzl-1(-)* (INK324) and *pzl-1(-) kss-2(-)* (INK485) mutant alleles generated using CRISPR/Cas in Chr. II NIL background (left). A frameshift mutation in *pzl-1* is sufficient to abolish delay in F_2_ offspring (top right). A *pzl-1(-) kss-2(-)* double mutant allele is poisoned by the WT allele (bottom right). *pzl-1(-) kss-2(-)* double mutant exhibits stronger phenotype – embryonic and larval arrest instead of developmental delay that was observed in crosses between Chr. II NIL and EG6180. **(B)** Relative position of *mec-15*, *zyg-9*, *fars-3* and *pzl-1* on Chr. II. **(C)** Resulting phenotype of L1 larvae following overexpression of KLMT-1 (middle) or PZL-1 (bottom) in early (up to 16 cell stage) embryos. Heat-shock induction in EG6180 control strain results in wild type L1 larvae (top). Similar to *pzl-1(-) kss-2(-)* double mutant, PZL-1 overexpression exhibits embryonic and larval arrest instead of developmental delay. **(D)** Volcano plot of 3xFLAG::KSS-2 co-immunoprecipitation followed by mass spectrometry (left). Construct is expressed under a constitutive promoter (*Ctr*-*rpl-36p*), strain – INK976. Members of the SCF complex are color-coded according to their function in the complex (right). Co-IPs were performed in biological triplicates. Lysate from Chr. II NIL served as a control for fold enrichment. **(E)** AlphaFold2 model of KSS-2 with solvent-exposed residue, Arg297, located on the inner surface of the solenoid. EG6180, a susceptible haplotype, carries a copy of *kss-2* with Leu74Met and Arg297Gly substitution. **(F)** We introduced Gly297Arg point mutation in the EG6180 *kss-2* locus using CRISPR/Cas (INK590), and crossed it to Chr. II NIL. EG-KSS-2[G297R] rescues the F_2_ lethality associated with the *pzl-1/kss-2* TA. Error bars indicate 95% confidence intervals calculated with the hybrid Wilson/Brown method. **(G)** Electrostatic potential of NIC-KSS-2 (top) and EG-KSS-2 (bottom), Alphafold2 models. Residue 297 is marked with an asterisk.

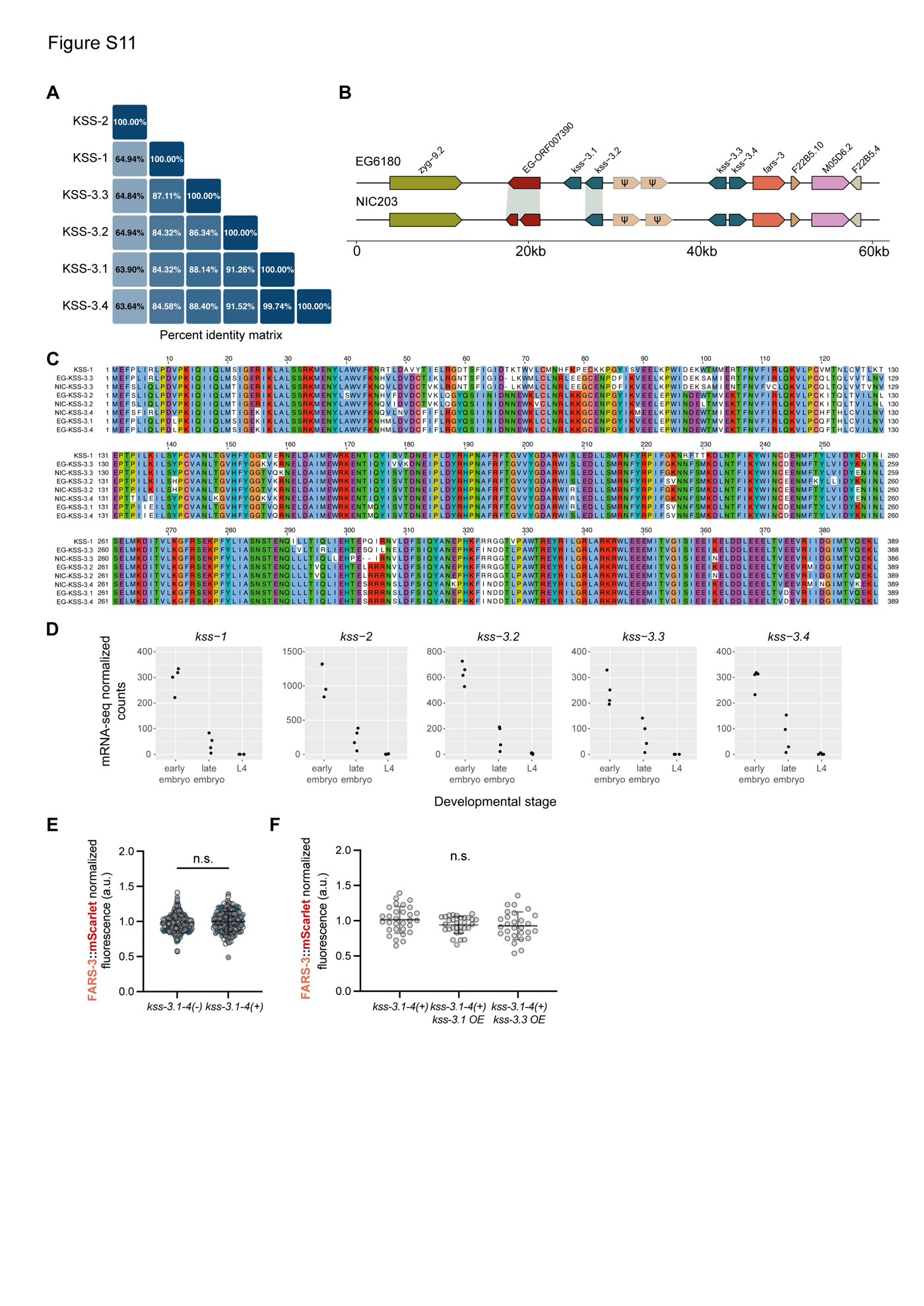

**Figure S11. A cluster of tandem F-box *kss* paralogs next to the *fars-3* locus. (A)** Protein sequence similarity among various KSS paralogs. Percent identity values are calculated in Clustal Omega. **(B)** Schematic representation of *fars-3* locus in EG6180 and NIC203. NIC203 has only three *kss-3* paralogs. **(C)** Multiple sequence alignment of KSS-1 and KSS-3 paralogs from EG6180 and NIC203. Colors correspond to the Clustal X Colour Scheme. **(D)** Expression levels of NIC203 *kss-1*, *kss-2*, *kss-3.2*, *kss-3.3*, and *kss-3.4* at three different developmental stages. Expression was quantified by mRNA-seq performed in biological quadruplicates. **(E)** Quantification of FARS-3::mScarlet levels in the wild type *kss-3.1-4(+)* and *kss-3.1-4(-)* quadruple mutant background (two-sided unpaired t-test; *P* = 0.0641, n.s. – not significant). Experiment was performed twice, individual repeats are coloured in gray and cyan. Error bars indicate mean with standard deviation. **(F)** Quantification of FARS-3::mScarlet levels in the wild type *kss-3.1-4(+)* background and in lines overexpressing *kss-3.1* or *kss-3.3* from single copy insertion transgenes (*Ctr-hsp-16.11p*). Difference between group means was analyzed using Brown-Forsythe ANOVA test (*F**(2, 73.38)= 2.094, *P* = 0.1305, n.s. – not significant). Error bars indicate mean with standard deviation.

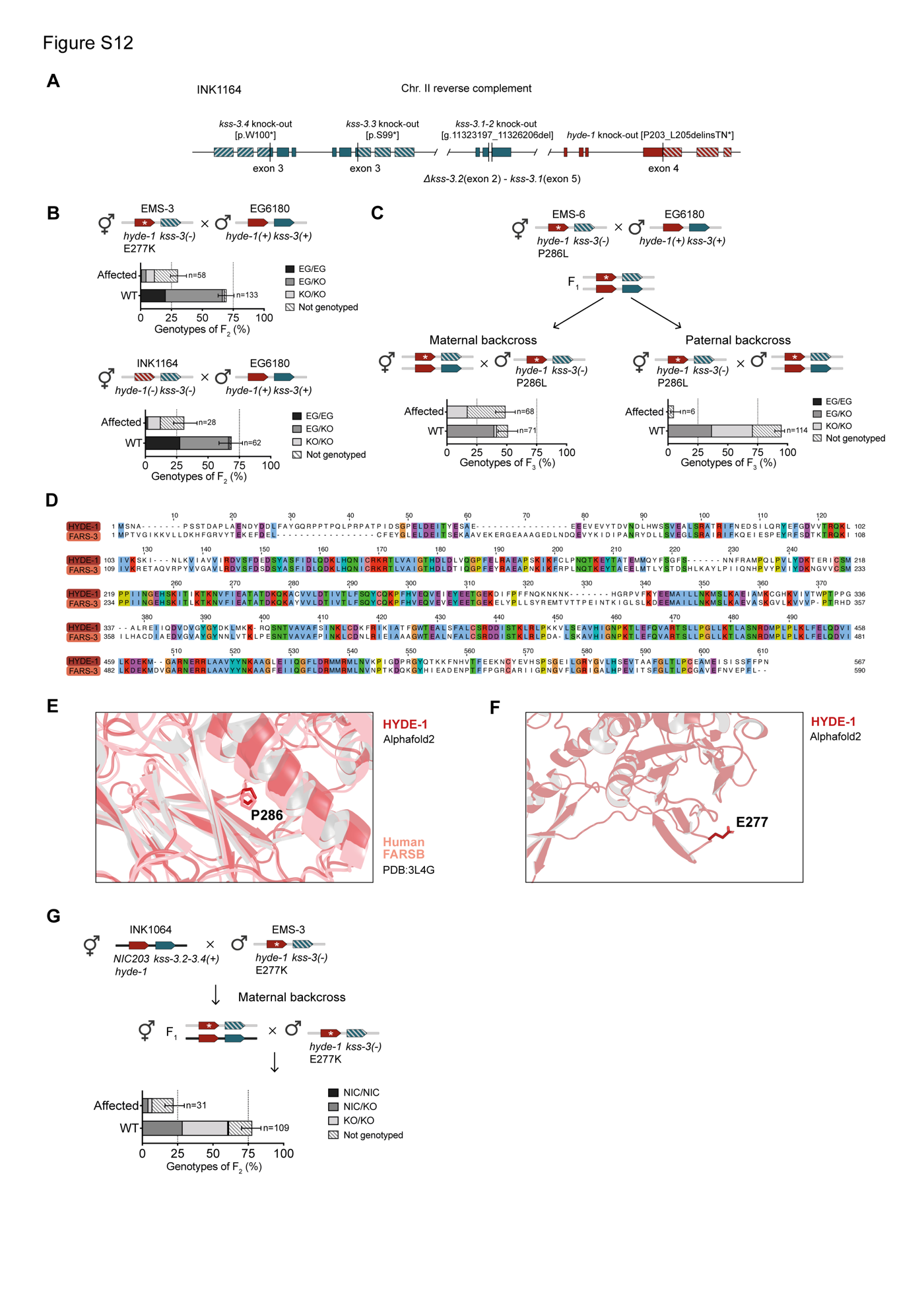

**Figure S12. *hyde-1/kss-3* is a maternal-effect TA. (A)** Schematic of the INK1164 strain carrying putative null mutations in *kss-3.*1, *kss-3.2*, *kss-3.3*, and *kss-3.4*, as well as in *ORF007390* (*hyde-1)*. All the mutant alleles were generated using CRISPR/Cas. **(B)** Crosses between the suppressor line EMS-3 (INK1092) and EG6180 (top) and the CRISPR/Cas-derived *hyde-1(-*) *kss-3.1-4(-)* mutant line (INK1164) with EG6180 (bottom). In both cases, we observed ~25% affected progeny and these were at large homozygous for the *hyde-1*(-) *kss-3.1-4(-)* mutant allele. KO/KO genotype corresponds to *hyde-1*[E277K] *kss-3.1-4(-)* or *hyde-1*(-) *kss-3.1-4(-).* **(C)** Diagnostic genetic crosses to determine whether *hyde-1* is a maternal-effect or paternal-effect toxin. In a maternal backross, heterozygous F_1_ hermaphrodites, derived from a cross between EMS-6 line (*hyde-1*[P286L] *kss-3.1-4(-),* INK1095) and EG6180, are mated with EMS-6 males (left). In a paternal backross, heterozygous F_1_ males are mated with EMS-6 hermaphrodites (right). If the toxin is maternal-effect (the toxin is deposited in eggs and not transmitted by sperm) then the TA is active in a maternal backcross (expected 50% lethality) but inactive in a paternal backcross. KO/KO genotype corresponds to *hyde-1*[P286L] *kss-3.1-4(-).* **(D)** Protein sequence alignment of HYDE-1 and *C. tropicalis* FARS-3. Colors correspond to the Clustal X Colour Scheme. **(E)** Structural alignment of HYDE-1 (Alphafold2 model) and human FARSB (PDB: 2L4G) highlighting HYDE-1 Pro286. This residue is highly conserved and mutated in the suppressor line EMS-6 (Pro286Leu). **(F)** HYDE-1 predicted structure (Alphafold2) highlighting the residue Glu277. This solvent exposed residue is mutated in the suppressor line EMS-3 (Glu277Lys). **(G)** *NIC-ORF007668* codes for HYDE-1 in NIC203. It shares only 56% of protein sequence identity to HYDE-1 toxin identified in EG6180 and carries a premature stop codon (see fig. S11B). To test whether the *NIC-hyde-1* behaves as a toxin, we crossed hermaphrodites of a NIC203 strain (INK1064) in which all known toxins (*pzl-1*, s*low-1*, and *klmt-1*) had been mutated using CRISPR/Cas with males of the suppressor EMS-3 line (*hyde-1*[E277K] *kss-3.1-4(-),* INK1092) that carries an inactive *hyde-1/kss-3* TA. Then, heterozygous F_1_ hermaphrodites were mated with EMS-3 males and the phenotype and genotype of their progeny was determined. If NIC203 *hyde-1* was active, then we would expect to see only heterozygous individuals among the F_2_. However, homozygous *hyde-1* mutants corresponding to EMS-3 allele were observed in a 1:1 ratio as heterozygous, indicating that HYDE-1 from NIC203 is not a toxin. KO/KO genotype corresponds to *hyde-1*[E277K] *kss-3.1-4(-).*

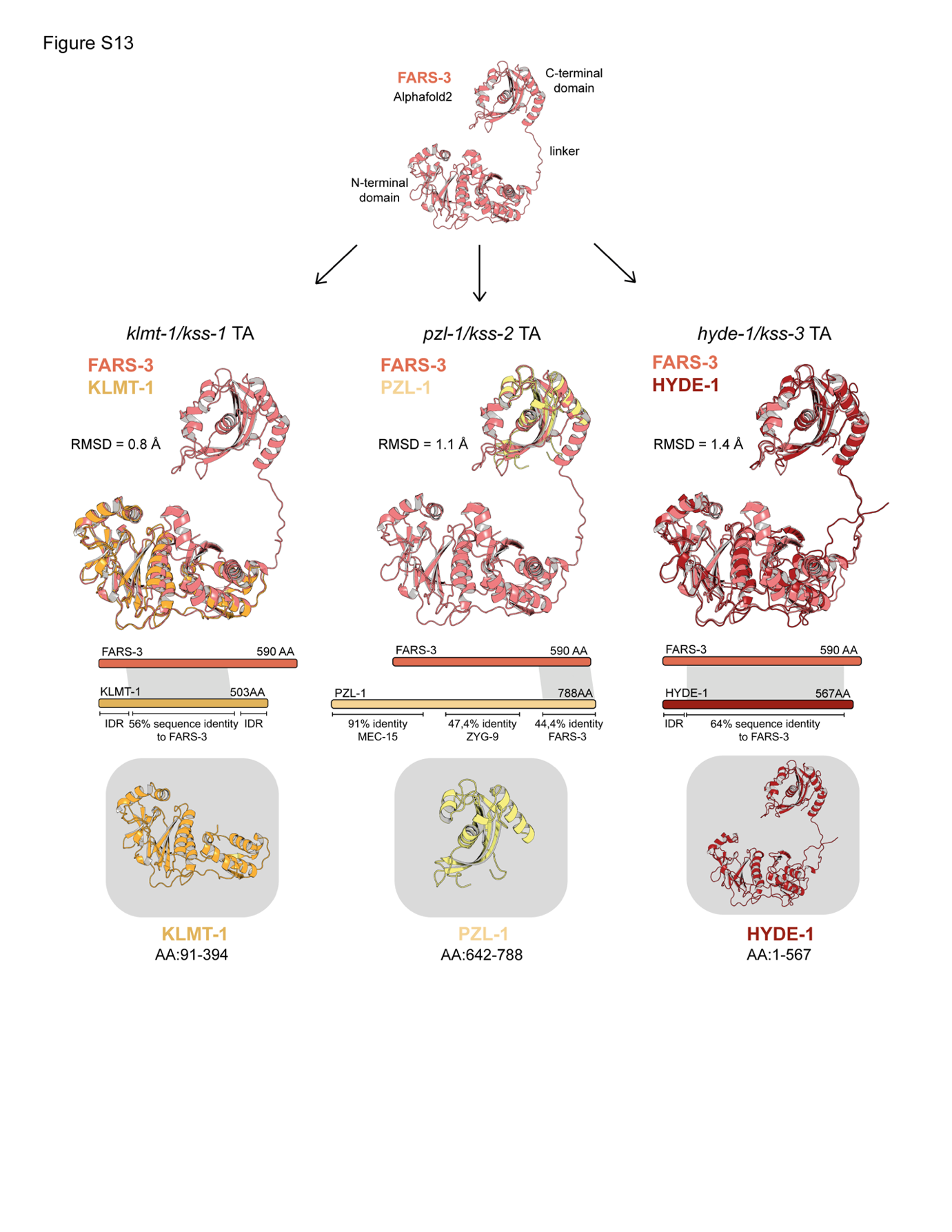

**Figure S13. KLMT-1, PZL-1, and HYDE-1 have unique FARS-3-derived domain architectures.** Comparison of the Alphafold2 predicted structures of the toxins KLMT-1, PZL-1, and HYDE-1 to *C. tropicalis* FARS-3. The structure of KLMT-1 does not include the predicted N-terminal and C-terminal intrinsically disordered regions. The structure of PZL-1 only includes the C-terminal region with homology to FARS-3.

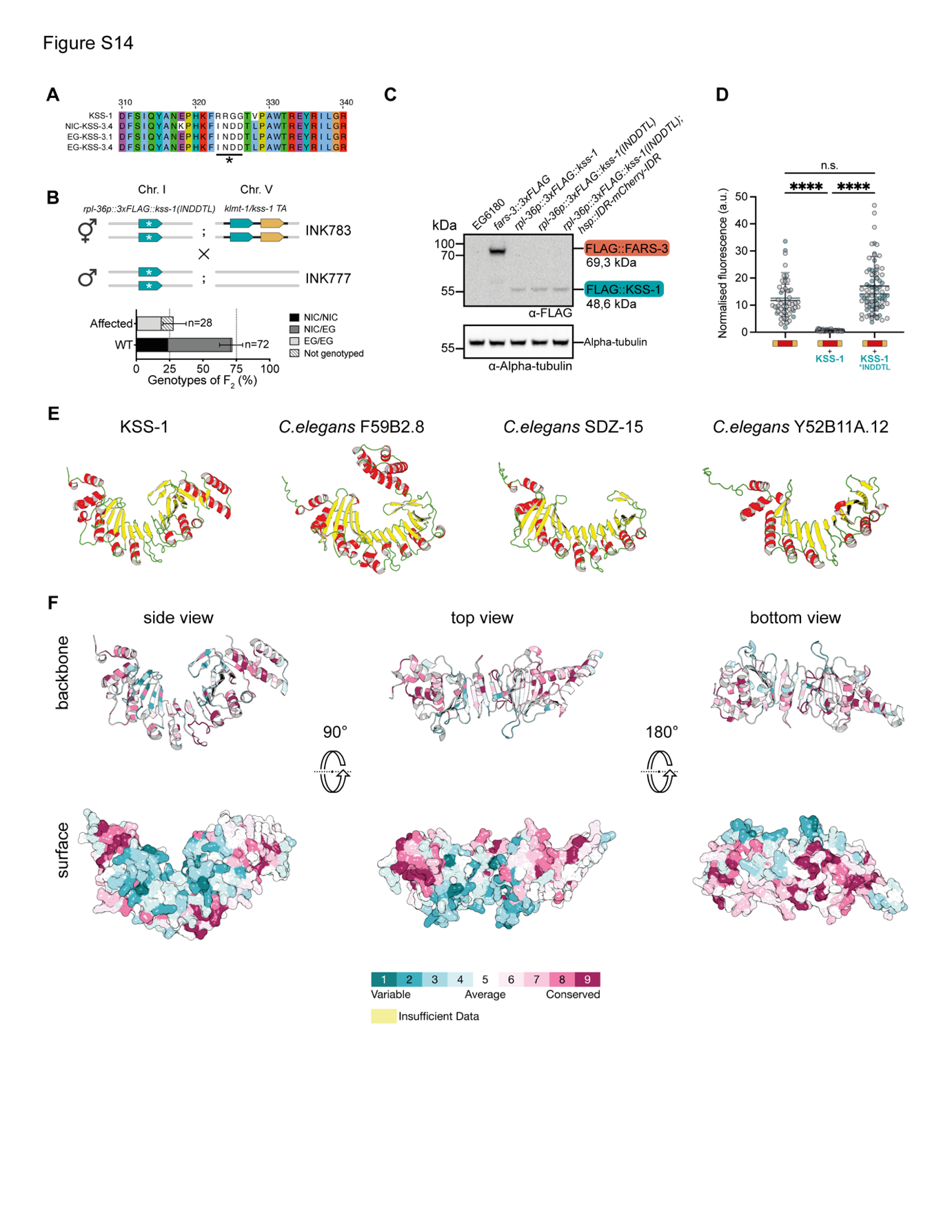

**Figure S14. A variable loop in KSS antidotes is necessary for KSS-1 substrate specificity. (A)** Multiple sequence alignment of KSS-1 and KSS-3 antidotes from EG6180 and NIC203. Asterisk marks one of the variable segments between KSS-1 and KSS-3 that is located on a loop opposing the inner surface of the solenoid. Colors correspond to the Clustal X Colour Scheme. **(B)** Overexpression of 3xFLAG::KSS-1(INDDTL) mutant from a single copy transgene does not rescue the F_2_ lethality associated with the *klmt-1/kss-1* TA. The transgene is integrated in a synthetic landing pad on Chr. I. Both parental lines, INK783 (Chr. V NIL background) and INK777 are homozygous carriers for the transgene. Error bars indicate 95% confidence intervals calculated with the hybrid Wilson/Brown method. **(C)** Western blot confirming the expression of 3xFLAG::KSS-1(INDDTL) from the transgenic line used in rescue experiments. 3xFLAG::KSS-1(INDDTL) is also expressed when introduced into a different background. An endogenously tagged FARS-3::3xFLAG and overexpression of 3xFLAG::KSS-1 were used as positive controls (INK505 and INK563). Negative control is the EG6180 parental strain. Western blot against alpha-tubulin serves as a loading control. **(D)** Fluorescent reporter assays to test if KSS-1(INDDTL) can mediate degradation of mCherry with KLMT-1 IDRs. Fluorescent mCherry reporter is under the control of a heat-shock inducible promoter (*Ctr-hsp-16.11p*) whereas the antidote is expressed under a constitutive promoter (*Ctr*-*rpl-36p*), strain – INK806. KSS-1(INDDTL) mutant does not recognise mCherry fused with IDRs (P = 0.1340). Difference between group means was analyzed using Kruskal-Wallis test (*H*(3) = 112.5, *P* < 0.0001), followed by Dunn's post hoc test. Error bars indicate mean with standard deviation. Experiment was performed twice, individual repeats are coloured in gray and cyan. **(E)** Examples of *C. elegans* proteins identified by Foldseek to be structurally homologous to *C. tropicalis* KSS-1. Alpha-helices are shown in red and beta strands in yellow. Full list available in Data S8 **(F)** Conservation pattern of the KSS antidotes and KSL paralogues generated using ConSurf. MSA included all *C. tropicalis* genes shown in Fig. 5A. Conserved amino acids are colored bordeaux, and variable amino acids are turquoise.

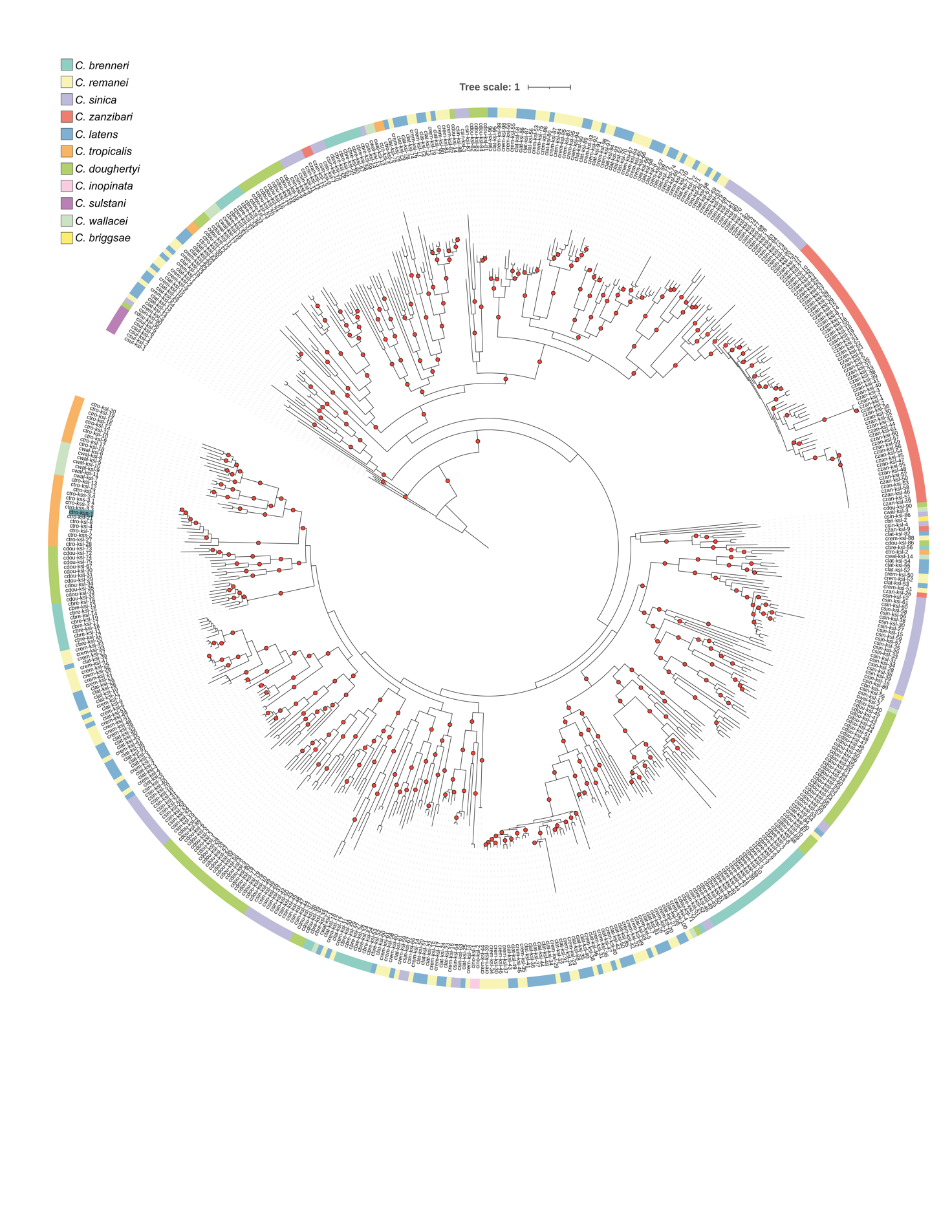

**Figure S15. Phylogenetic tree of KSL proteins across nematodes.** Phylogenetic relationships of 560 KSL proteins identified across *Caenorhabditis* species. Red dots denote branches with bootstrap values of SH-aLRT ≥ 80% and UFboot ≥ 95%. We filtered the resulting nucleotide multiple sequence alignment to retain only nucleotide positions covered by at least 80% of the sequences. Each color represents a different species (see inset).

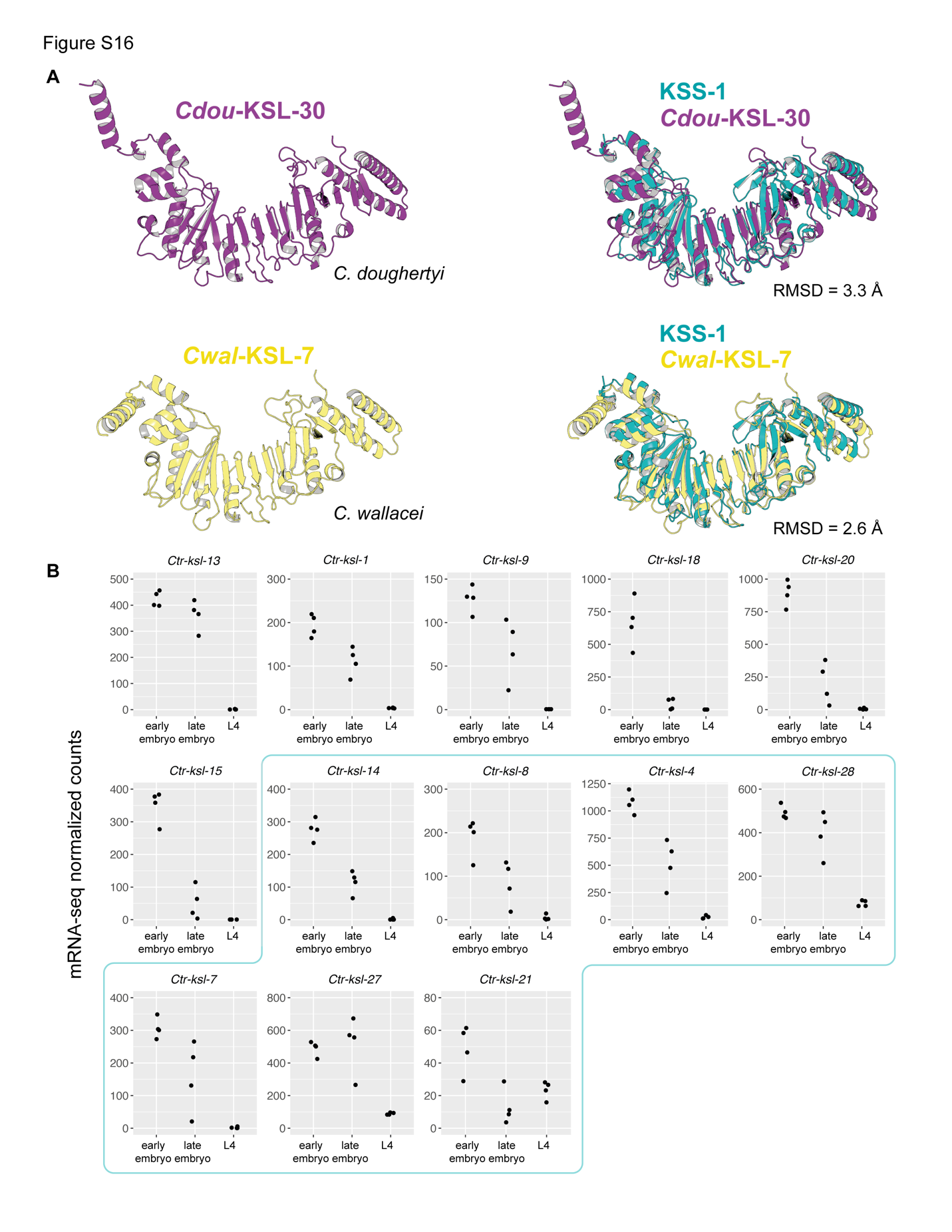

**Figure S16. KSL paralogs have a similar structure and expression pattern to KSS antidotes. (A)** Alphafold2 models of *Cdou-*KSL-30 and *Cwal-*KSL-7 (left). Predicted structures of both paralogs are highly similar to KSS-1 (right). **(B)** Expression levels of *ksl* paralogues identified on Chr. II of *C. tropicalis* (Fig. 5A). Paralogues that form a clade with characterized antidotes are marked in cyan. Expression was quantified by mRNA-seq in biological quadruplicates.

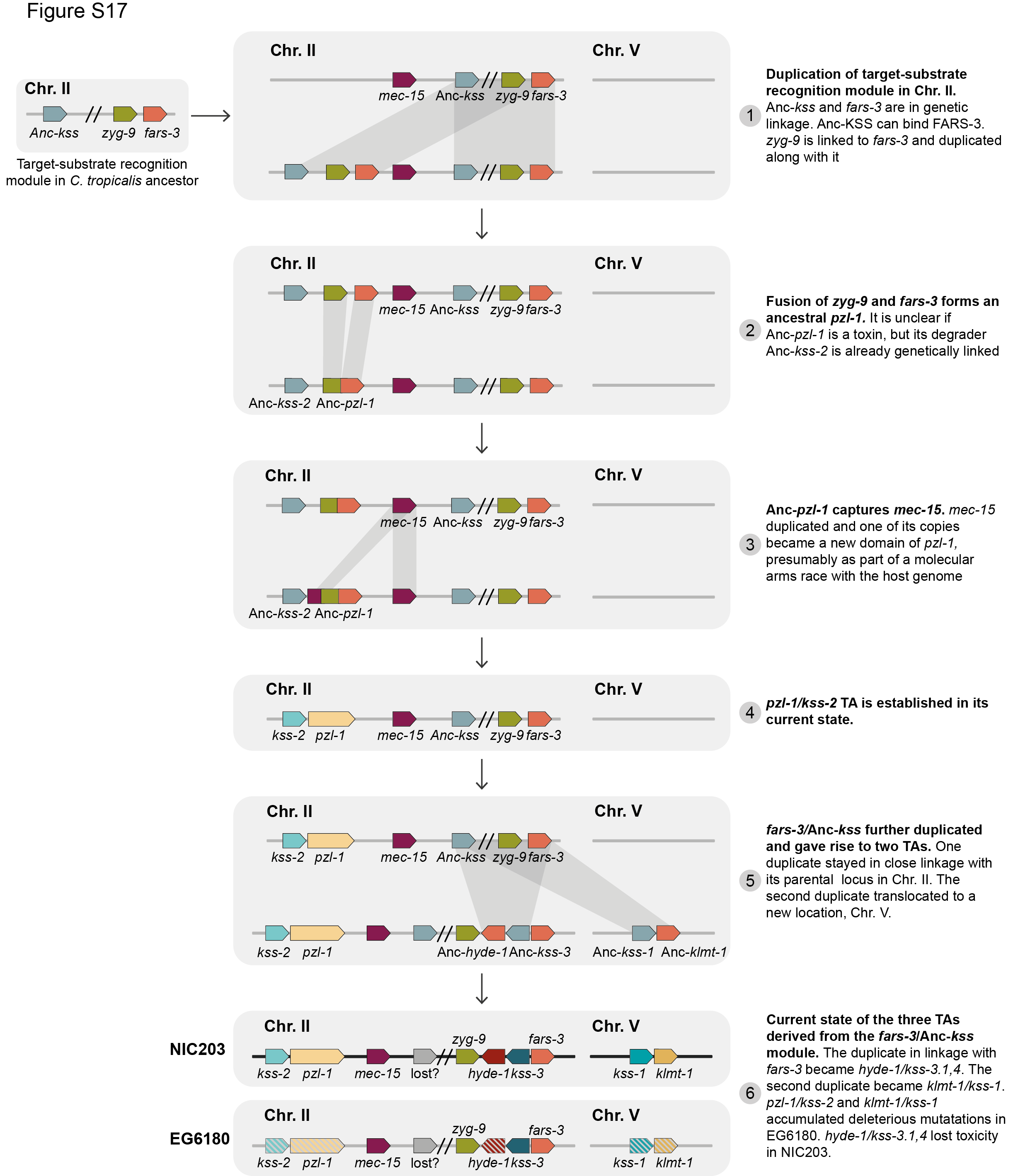

**Figure S17. Evolutionary scenarios leading to the emergence of three TA elements from *fars-3*.** Following the split of *C. wallacei* and *C. tropicalis*, a *C. tropicalis* *ksl* F-box protein in linkage with *fars-3* randomly evolved affinity for FARS-3 and became the ancestor of all *kss* antidotes (Anc*-kss*). (1) The Chr. II *fars-3/*Anc*-kss* module duplicated a first time, along with its neighboring gene, *zyg-9,* located upstream of *fars-3.* (2) The reduced selective pressure experienced by the duplicate module facilitated the evolution of a TA element. We speculate that *Anc-pzl-1* (a chimeric protein of *zyg-9* and *fars-3*) was already mildly toxic at this stage, and the Anc-*kss* paralog became Anc-*kss-2*. (3) After the formation of the first TA system, Anc-*pzl-1* fused with a paralog of *mec-15*, potentially resulting in an increase in pzl-1 toxicity. (4) Current state of *pzl-1/kss-2* TA. (5) Either a second duplication of the *fars-3/*Anc*-kss* module gave rise to a common ancestor of the *klmt-1/kss-1* and *hyde-1/kss-3* TAs, or these TA systems evolved independently from two separate duplication events of the parental module. (6) The *klmt-1/kss-1* TA translocated to Chr. V, whereas the *hyde-1/kss-3* TA remained linked to its parental locus in Chr. II, providing an evolutionary advantage to *fars-3* through genetic hitchhiking. Anc-kss likely became pseudogenized and was lost over time, as it did not confer a selective advantage to its host. For simplicity, only one antidote of hyde-1 is shown.

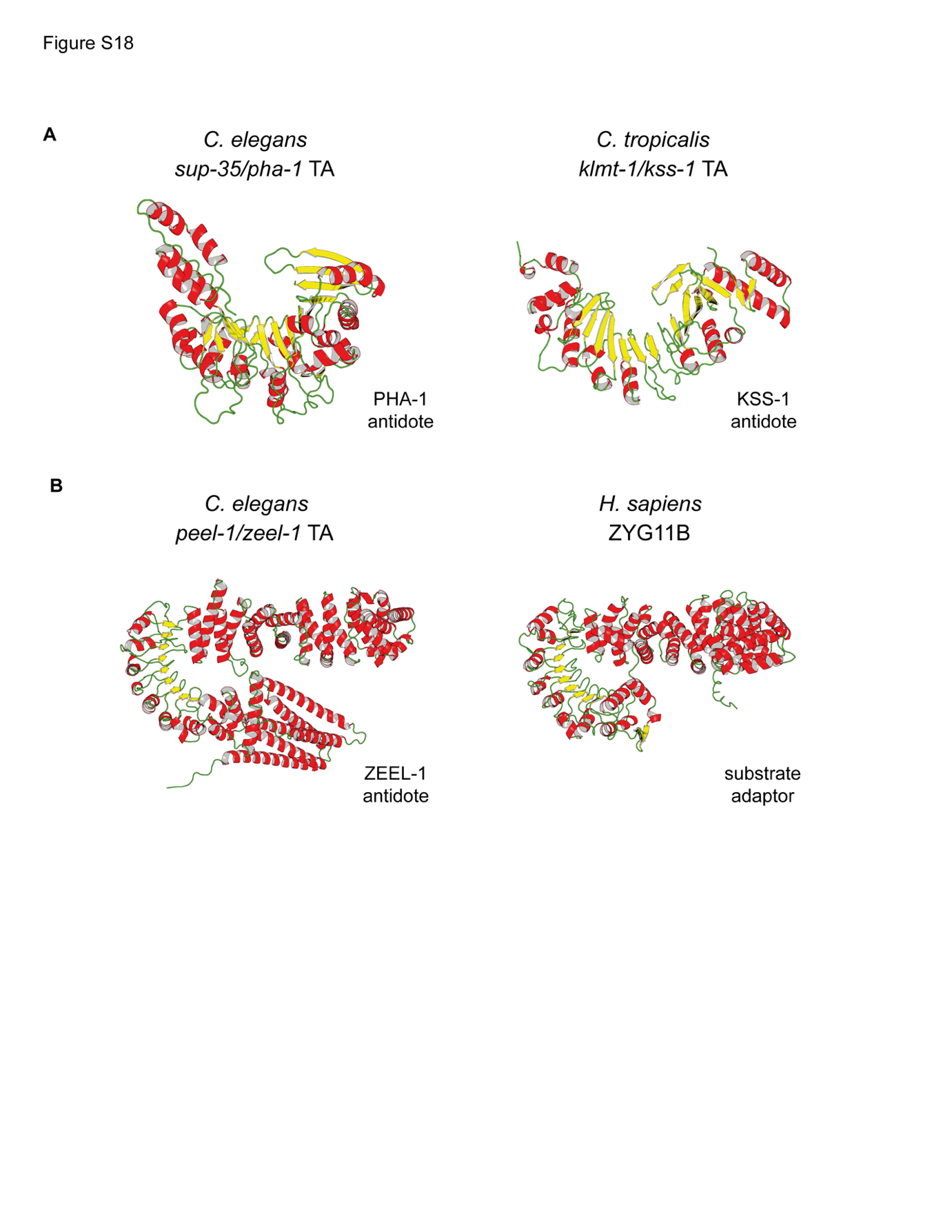

**Figure S18. *C. elegans* antidotes could also function as substrate-recognition modules of E3 ligases. (A)** Predicted Alphafold2 structures of PHA-1 and KSS-1. PHA-1 is the antidote of the *C. elegans sup-35/pha-1* TA. The molecular mechanism of PHA-1 antidote activity is currently unknown. However, PHA-1 is predicted to adopt a horseshoe fold with parallel beta sheets running on its inner face and alpha helices on its exterior, analogous to KSS-1, suggesting that it may bind and degrade SUP-35. **(B)** Predicted structures of ZEEL-1 and human ZYG11B. ZEEL-1 is the antidote of the *C. elegans peel-1/zeel-1* TA. The molecular mechanism of ZEEL-1 antidote activity is currently unknown. However, sequence and structural similarity strongly suggest a link with protein degradation. ZEEL-1 is homologous to ZYG11B, which serves as a substrate adapter subunit in the E3 ubiquitin ligase complex ZYG11B–CUL2–Elongin BC.

**Table S1. Raw data for all phenotyping of lines used in this study.**

| Strain | Short name | Mothers screened | Total number of embryos | Embryonic lethal | Larval arrest | Delay | Arrest | Sterile | Other | Wild-type (WT) | % of WT | Comments |
| --- | --- | --- | --- | --- | --- | --- | --- | --- | --- | --- | --- | --- |
| QX2341 | Chr. II NIL | 10 | 100 | 4 | 0 | 2 | 1 | 0 | 0 | 93 | 93,0 |  |
| QX2343 | Chr. V NIL | 10 | 100 | 1 | 0 | 0 | 0 | 0 | 0 | 99 | 99,0 |  |
| INK303 | NIC-ORF015419(-) = *klmt-1(-)* | 10 | 100 | 2 | 0 | 3 | 0 | 0 | 0 | 95 | 95,0 |  |
| INK324 | NIC-ORF006816 (-) = *pzl-1(-)* | 10 | 100 | 0 | 2 | 2 | 3 | 0 | 0 | 93 | 93,0 |  |
| INK422 | NIC-ORF015419(-) NIC-ORF015420(-) = *klmt-1(-) kss-1(-)* | 10 | 100 | 4 | 0 | 0 | 0 | 0 | 0 | 96 | 96,0 |  |
| INK485 | NIC-ORF006816 (-) NIC-ORF006815 (-) = *plz-1(-) kss-2(-)* | 10 | 100 | 1 | 1 | 1 | 0 | 0 | 0 | 97 | 97,0 |  |
| INK722 | FARS-3::mScarlet | 8 | 119 | 0 | 0 | 1 | 1 | 1 | 0 | 116 | 97,5 |  |
| INK793 | *kss-3.4(-)* | 8 | 80 | 1 | 1 | 0 | 0 | 0 | 0 | 78 | 97,5 |  |
| INK801 | *kss-3.3(-) kss-3.4(-)* | 10 | 100 | 1 | 0 | 6 | 2 | 0 | 0 | 91 | 91,0 |  |
| INK951 | *kss-3.1(-) kss-3.2(-)* | 11 | 110 | 1 | 0 | 0 | 0 | 0 | 0 | 109 | 99,1 |  |
| INK957 | *kss-3.2(-) kss-3.3(-) kss-3.4(-)* | 8 | 80 | 3 | 0 | 0 | 0 | 0 | 0 | 77 | 96,3 |  |
| INK1164 | *hyde-1(-) kss-3.1-3.4(-)* | 10 | 100 | 4 | 1 | 3 | 0 | 0 | 0 | 92 | 92,0 |  |
| INK1088 | mScarlet::ZYG-9.2 | 10 | 100 | 2 | 0 | 3 | 0 | 0 | 0 | 95 | 95,0 |  |
| INK1169 | *kss-3.1(-) kss-3.2(-) kss-3.3(-)* | 10 | 100 | 1 | 1 | 1 | 0 | 0 | 3 | 94 | 94,0 | Other = worms not found |
| INK169 | KLMT-1::mNeonGreen = KLMT-1::mNG | 10 | 100 | 2 | 0 | 1 | 0 | 1 | 2 | 94 | 94,0 | Other = worms not found |
| INK206 | KLMT-1::mNeonGreen;  *kss-1(-)* | 10 | 100 | 1 | 0 | 2 | 0 | 1 | 2 | 94 | 94,0 | Other = worms not found |
| INK950 | *kss-3.1-3.4(-)*; FARS-3::mScarlet | 18 | 180 | 62 | 41 | 23 | 3 | 10 | 0 | 41 | 22,8 | Day 1 - 52 affected larvae, day 4 - 11/52 wild type worms |
| INK1196 | *kss-3.1(-) kss-3.4(-)* | 10 | 100 | 69 | 15 | 2 | 3 | 0 | 0 | 11 | 11,0 | Day 1 - 24 affected larvae, day 4 - 4/24 wild type worms, 5/24 delay and arrest |
| INK1172* | *kss-3.1(-) kss-3.2(-) kss-3.4(-)* | 4 | 32 | - | - | - | - | - | - | - | - | See footnote |
| INK1091 | EMS-2 | 4 | 60 | 8 | 2 | 0 | 2 | not screened | 0 | 48 | 80,0 | Revertant, sequenced |
| INK1092 | EMS-3; *hyde-1(abu529[E277K]) II* | 4 | 61 | 5 | 0 | 0 | 2 | not screened | 0 | 54 | 88,5 | Revertant, sequenced |
| INK1095 | EMS-6 abu532; *hyde-1(abu532[P286L]) II* | 4 | 61 | 2 | 0 | 1 | 0 | not screened | 0 | 58 | 95,1 | Revertant, sequenced |
| INK1097 | EMS-8 | 4 | 60 | 5 | 2 | 0 | 0 | not screened | 0 | 53 | 88,3 | Revertant, sequenced |
| INK1100 | EMS-11 | 4 | 60 | 5 | 0 | 1 | 0 | not screened | 0 | 54 | 90,0 | Revertant, sequenced. |
| EG6180 | EG6180 | 10 | 100 | 2 | 0 | 0 | 1 | 0 | 0 | 97 | 97,0 | Wild type control |
| INK950 | *kss-3.1-3.4(-);* FARS-3::mScarlet | 18 | 180 | 62 | 41 | 23 | 3 | 10 | 0 | 41 | 22,8 | Day 1 - 52 affected larvae, day 4 - 11/52 wild type worms |

*We aimed to knock-out *kss-3.4* in the backrground of *kss-3.1(-) kss-3.1(-)* using CRISPR/Cas. After injection we selected 4 heterozygous for *kss-3.4(-)* hermaphrodites, propagated them and singled 32 offspring on individual plates, expecting to recover ~25% of desired genotype according to Mendelian segregation. 4 days later we genotyped 32 lines: 21 were heterozygous for *kss-3.4(-)* allele, 11 homozygous for wild type allele, and we did not recover *kss-3.4(-)* homozygous offspring, which suggested that this allele in homozygous state was detrimental. We decided not to proceed with isolating affected offspring as described in "Generation of *C. tropicalis* transgenic lines" Methods section.

**Table S2. Raw data for KLMT-1 injection experiments into adult gonads.** Co-injection marker helps to visually distinguish the offspring that received an injection dose.

| **Worm number** | **Injection mix** | **Strain** | **Total number of eggs** | **With co-injection marker** | | | | **Without co-injection marker** | | | |
| --- | --- | --- | --- | --- | --- | --- | --- | --- | --- | --- | --- |
|  |  |  |  | **Total** | **Affected** | **WT** | **% affected** | **Total** | **Affected** | **WT** | **% affected** |
| 1 | Buffer | EG6180 | 136 | 38 | 0 | 38 | 0,0 | 98 | 0 | 98 | 0,0 |
| 2 | Buffer | EG6180 | 110 | 36 | 1 | 35 | 2,8 | 74 | 1 | 73 | 1,4 |
| 3 | Buffer | EG6180 | 118 | 49 | 1 | 48 | 2,0 | 69 | 0 | 69 | 0,0 |
| 4 | Buffer | EG6180 | 131 | 37 | 1 | 36 | 2,7 | 94 | 0 | 94 | 0,0 |
| 5 | Buffer | EG6180 | 137 | 31 | 0 | 31 | 0,0 | 106 | 8 | 98 | 7,5 |
| 6 | Buffer | EG6180 | 110 | 40 | 0 | 40 | 0,0 | 70 | 2 | 68 | 2,9 |
| 7 | Buffer | EG6180 | 109 | 38 | 0 | 38 | 0,0 | 71 | 0 | 71 | 0,0 |
| 8 | Buffer | EG6180 | 65 | 24 | 0 | 24 | 0,0 | 41 | 7 | 34 | 17,1 |
| 9 | Buffer | EG6180 | 138 | 42 | 1 | 41 | 2,4 | 96 | 2 | 94 | 2,1 |
| 10 | Buffer | EG6180 | 119 | 33 | 0 | 33 | 0,0 | 86 | 1 | 85 | 1,2 |
| 11 | Buffer | EG6180 | 131 | 23 | 0 | 23 | 0,0 | 108 | 3 | 105 | 2,8 |
| 12 | Buffer | EG6180 | 117 | 19 | 0 | 19 | 0,0 | 98 | 3 | 95 | 3,1 |
| 13 | Buffer | EG6180 | 189 | 26 | 0 | 26 | 0,0 | 163 | 2 | 161 | 1,2 |
| 14 | KLMT-1 | EG6180 | 96 | 28 | 19 | 9 | 67,9 | 68 | 7 | 61 | 10,3 |
| 15 | KLMT-1 | EG6180 | 116 | 41 | 19 | 22 | 46,3 | 75 | 8 | 67 | 10,7 |
| 16 | KLMT-1 | EG6180 | 118 | 45 | 36 | 9 | 80,0 | 73 | 1 | 72 | 1,4 |
| 17 | KLMT-1 | EG6180 | 133 | 34 | 30 | 4 | 88,2 | 99 | 1 | 98 | 1,0 |
| 18 | KLMT-1 | EG6180 | 102 | 32 | 27 | 5 | 84,4 | 70 | 5 | 65 | 7,1 |
| 19 | KLMT-1 | EG6180 | 110 | 52 | 44 | 8 | 84,6 | 58 | 3 | 55 | 5,2 |
| 20 | KLMT-1 | EG6180 | 141 | 40 | 36 | 4 | 90,0 | 101 | 3 | 98 | 3,0 |
| 21 | KLMT-1 | EG6180 | 150 | 31 | 25 | 6 | 80,6 | 119 | 5 | 114 | 4,2 |
| 22 | KLMT-1 | EG6180 | 69 | 36 | 32 | 4 | 88,9 | 33 | 4 | 29 | 12,1 |
| 23 | KLMT-1 | EG6180 | 89 | 26 | 21 | 5 | 80,8 | 63 | 2 | 61 | 3,2 |
| 24 | KLMT-1 | EG6180 | 123 | 28 | 23 | 5 | 82,1 | 95 | 9 | 86 | 9,5 |
| 25 | KLMT-1 | EG6180 | 176 | 39 | 29 | 10 | 74,4 | 137 | 8 | 129 | 5,8 |
| 26 | KLMT-1 | EG6180 | 145 | 42 | 32 | 10 | 76,2 | 103 | 5 | 98 | 4,9 |
| 27 | KLMT-1 | EG6180 | 116 | 27 | 19 | 8 | 70,4 | 89 | 6 | 83 | 6,7 |
| 28 | KLMT-1 | EG6180 | 136 | 18 | 13 | 5 | 72,2 | 118 | 3 | 115 | 2,5 |
| 29 | KLMT-1 | EG6180 | 154 | 39 | 30 | 9 | 76,9 | 115 | 8 | 107 | 7,0 |
| 30 | KLMT-1 | EG6180 | 165 | 32 | 25 | 7 | 78,1 | 133 | 8 | 125 | 6,0 |
| 31 | KLMT-1 | EG6180 | 134 | 38 | 28 | 10 | 73,7 | 96 | 10 | 86 | 10,4 |
| 32 | KLMT-1 | INK563 (*kss-1* OE) | 101 | 42 | 0 | 42 | 0,0 | 59 | 3 | 56 | 5,1 |
| 33 | KLMT-1 | INK563 (*kss-1* OE) | 103 | 10 | 6 | 4 | 60,0 | 93 | 2 | 91 | 2,2 |
| 34 | KLMT-1 | INK563 (*kss-1* OE) | 153 | 39 | 0 | 39 | 0,0 | 114 | 1 | 113 | 0,9 |
| 35 | KLMT-1 | INK563 (*kss-1* OE) | 101 | 35 | 0 | 35 | 0,0 | 66 | 1 | 65 | 1,5 |
| 36 | KLMT-1 | INK563 (*kss-1* OE) | 150 | 23 | 0 | 23 | 0,0 | 127 | 1 | 126 | 0,8 |
| 37 | KLMT-1 | INK563 (*kss-1* OE) | 154 | 44 | 1 | 43 | 2,3 | 110 | 1 | 109 | 0,9 |
| 38 | KLMT-1 | INK563 (*kss-1* OE) | 3 | 0 | 0 | 0 | 0,0 | 3 | 0 | 3 | 0,0 |
| 39 | KLMT-1 | INK563 (*kss-1* OE) | 128 | 46 | 0 | 46 | 0,0 | 82 | 3 | 79 | 3,7 |

**Table S3. List of all *C. tropicalis* strains used in this study.** All strains except QX2341, QX2343, NIC203 and EG6180 were generated for this study.

| Strain | Short name | Genotype | First used in | Description |
| --- | --- | --- | --- | --- |
| QX2341 | Chr. II NIL | *qqIR45 (II:8.0-8.8 Mb; NIC203 > EG6180); EG6180 Mito* | Fig. 3A | NIL carrying NIC203 TA element on Chr. II in a EG6180 background. Source - Ben-David et al (2021) |
| QX2343 | Chr. V NIL | *qqIR47 (V:1.3-1.8 Mb; NIC203 > EG6180); EG6180 Mito* | Fig, 1B | NIL carrying NIC203 TA element on Chr. V in a EG6180 background. Source - Ben-David et al (2021) |
| NIC203 | NIC203 | wild type | Fig. 1A | Wild isolate from Capesterre Belle-Eau, Guadeloupe. Lat 16.05 Lon -61.63. Found in rotting flowers by N. Poullet and C. Braendle. Source - Christian Braendle |
| EG6180 | EG6180 | wild type | Fig. 1A | Wild isolate from El Yunque, Puerto Rico. Lat 18.3 Lon -65.8. Found in rotting fruit by M. Ailion and E. Jorgensen. Source - Christian Braendle |
| INK285 | Chr. IV SLP | *abuSi9[dpy-10 & sup-35 gRNA targets::(HygR(p.52-341)::rps-20 3' UTR::LoxP, IV:6.9Mb]* | - | Chr. IV synthetic landing pad with split hygromycine resistance, EG6180 background. Permissible only for somatic expression |
| INK943 | Chr. I SLP | *abuSi33[dpy-10 & sup-35 gRNA targets::(HygR(p.52-341)::rps-20 3' UTR::LoxP, I:7.5Mb]* | - | Chr. I synthetic landing pad with split hygromycine resistance, EG6180 background. Permissible only for somatic expression |
| INK63 | 3xFLAG::KLMT-1 | *klmt-1(abu32[3xFLAG::klmt-1]) V; qqIR47* | S3F | N-terminal 3xFLAG tagged KLMT-1, QX2343 background |
| INK66 | KLMT-1::3xFLAG | *klmt-1(abu33[klmt-1::3xFLAG]) V; qqIR47* | Fig. 2J | C-terminal 3xFLAG tagged KLMT-1, QX2343 background |
| INK169 | KLMT-1::mNeonGreen = KLMT-1::mNG | *klmt-1(abu107[klmt-1::2xTy1::mNeonGreen]) V; qqIR47* | Fig. 2G | C-terminal mNeonGreen tagged KLMT-1 (with 2xTy1 linker), QX2343 background |
| INK206 | KLMT-1::mNeonGreen; *kss-1(-)* | *kss-1(abu125[p.R7Ffs*28]) V; klmt-1(abu107[klmt-1::2xTy1::mNeonGreen]) V; qqIR47* | Fig. 2G | *kss-1* knockout and C-terminal mNeonGreen tagged KLMT-1 (with 2xTy1 linker), QX2343 background |
| INK277 | ΔN-KLMT-1::3xFLAG | *klmt-1(abu173[klmt-1::3xFLAG(p.S2_T45del)]) V; qqIR47* | Fig. 2J | Deletion of N-terminal IDR (internally disordered region) of KLMT-1 |
| INK303 | *NIC-ORF015419(-) = klmt-1(-)* | *klmt-1(abu191[p.E74Gfs*50]) V; qqIR47* | Fig. 1C | *klmt-1* knockout, QX2343 background |
| INK310 | heat-shock inducible mCherry | *abuSi11[hsp-16.11p::mCherry::tbb-2 3' UTR + HygR(+); abuSi9] IV* | Fig. 2H | Heat-shock inducible expression of mCherry, Chr. IV landing pad, EG6180 background |
| INK318 | heat-shock inducible KLMT-1 | *abuSi19[hsp-16.11p::klmt-1::tbb-2 3' UTR + HygR(+); abuSi9] IV* | Fig. 3A | Heat-shock inducible expression of KLMT-1, Chr. IV landing pad, EG6180 background |
| INK324 | *NIC-ORF006816 (-) = pzl-1(-)* | *pzl-1(abu205[p.K446_W447delinsT*]) II; qqIR45* | S10A | *pzl-1* knockout, QX2341 background |
| INK336 | heat-shock inducible IDR-mCherry-IDR | *abuSi36[hsp-16.11::KLMT-1(p.M1_D69)::mCherry::KLMT-1(p.A460_S503)::tbb-2 3' UTR + HygR(+); abuSi9] IV* | Fig. 2H | Heat-shock inducible expression of mCherry fused with N- and C-IDRs of KLMT-1 |
| INK422 | *NIC-ORF015419(-) NIC-ORF015420(-) = klmt-1(-) kss-1(-)* | *klmt-1(abu191[p.E74Gfs*50]) V; kss-1(abu192[p.T229Efs*8]) V; qqIR47* | Fig. 1C | *klmt-1* and *kss-1* knockout, QX2343 background |
| INK438 | heat-shock inducible KSS-1 | *abuSi40[hsp-16.11::kss-1::tbb-2 3' UTR + HygR(+), abuSi33] I* | Fig. 3H | Heat-shock inducible expression of KSS-1, Chr. I landing pad, EG6180 background |
| INK444 | 3xFLAG::KSS-1 | *kss-1(abu278[3xFLAG::kss-1]) V; qqIR47* | S7D | N-terminal 3xFLAG tagged KSS-1, QX2343 background |
| INK485 | *NIC-ORF006816 (-) NIC-ORF006815 (-) = plz-1(-) kss-2(-)* | *kss-2(abu291[p.Y81*]) II; pzl-1(abu205[p.K446_W447delinsT*]) II; qqIR45* | S10A | *pzl-1 and kss-2* knockout, QX2341 background |
| INK505 | FARS-3::3xFLAG | *fars-3(abu302[fars-3:3xFLAG]) II* | S14C | C-terminal 3xFLAG tagged FARS-3, EG6180 background |
| INK512 | heat-shock inducible KSS-1; heat-shock inducible KLMT-1 | *abuSi40[hsp-16.11::kss-1::tbb-2 3' UTR + HygR(+), abuSi33] I; abuSi19[hsp-16.11::klmt-1::tbb-2 3' UTR + HygR(+); abuSi9] IV* | Fig. 3H | Heat-shock inducible expression of KSS-1, Chr. I landing pad, and KLMT-1, Chr. IV landing pad, EG6180 background. Obtained by crossing INK438 to INK318 |
| INK563 | 3xFLAG::KSS-1 (*Ctr-rpl-36p*) | *abuSi53[rpl-36p::3xFLAG::kss-1::rpl-36 3' UTR + HygR(+), abuSi33] I* | Fig. 1F | Ribosomal promoter driven expression of N-terminal 3xFLAG tagged KSS-1, EG6180 background |
| INK577 | PZL-1::3xFLAG | *pzl-1(abu342[pzl-1::3xFLAG]) II* | Fig. 3A | C-terminal 3xFLAG tagged PZL-1, QX2341 background |
| INK581 | KLMT-1-ΔC | *klmt-1(abu346[3xFLAG::klmt-1(p.A460_S503del)]) V; qqIR47* | Fig. 2J | Deletion of C-terminal IDR of KLMT-1 |
| INK586 | heat-shock inducible KSS-2 | *abuSi56[hsp-16.11::kss-2::tbb-2 3' UTR + HygR(+), abuSi33] I* | Fig. 3H | Heat-shock inducible expression of KSS-2, Chr. I landing pad, EG6180 background |
| INK590 | EG-KSS-2[G297R] | *kss-2(abu335[p.G297R]) II* | S10F | EG6180 KSS-2 with glycine 297 changed to arginine, EG6180 background |
| INK629 | heat-shock inducible KSS-2; heat-shock inducible KLMT-1 | *abuSi56[hsp-16.11::kss-2::tbb-2 3' UTR + HygR(+), abuSi33] I; abuSi19[hsp-16.11::klmt-1::tbb-2 3' UTR + HygR(+); abuSi9] IV* | Fig. 3H | Heat-shock inducible expression of KSS-2, Chr. I landing pad and KLMT-1, Chr. IV landing pad, EG6180 background. Obtained by crossing INK586 to INK318 |
| INK631 | 3xFLAG::KSS-1 *(Ctr-rpl-36p)*; QX2343 | *abuSi53[rpl-36p::3xFLAG::kss-1::rpl-36 3' UTR + HygR(+), abuSi33] I; qqIR47* | S1E | Ribosomal promoter driven expression of N-terminal 3xFLAG tagged KSS-1, Chr. I landing pad, QX2343 background. Obtained by crossing INK563 to QX2343 |
| INK680 | heat-shock inducible PZL-1 | *abuSi66[hsp-16.11::pzl-1::tbb-2 3' UTR + HygR(+); abuSi9] IV* | Fig. 3A | Heat-shock inducible expression of PZL-1, Chr. IV landing pad, EG6180 background |
| INK722 | FARS-3::mScarlet | *fars-3(abu397[FARS-3::mScarlet]) II* | Fig. 4A | C-terminal mScarlet tagged FARS-3, EG6180 background. Used as a background for INK793, INK801, INK950, INK951, INK957 |
| INK726 | heat-shock inducible KSS-2; heat-shock inducible PZL-1 | *abuSi56[hsp-16.11::kss-2::tbb-2 3' UTR + HygR(+), abuSi33] I; abuSi66[hsp-16.11::pzl-1::tbb-2 3' UTR + HygR(+); abuSi9] IV* | Fig. 3H | Heat-shock inducible expression of KSS-2, Chr. I landing pad, and PZL-1, Chr. IV landing pad, EG6180 background. Obtained by crossing INK586 to INK680 |
| INK732 | heat-shock inducible KSS-1; heat-shock inducible PZL-1 | *abuSi40[hsp-16.11::kss-1::tbb-2 3' UTR + HygR(+), abuSi33] I; abuSi66[hsp-16.11::pzl-1::tbb-2 3' UTR + HygR(+); abuSi9] IV* | Fig. 3H | Heat-shock inducible expression of KSS-1, Chr. I landing pad, and PZL-1, Chr. IV landing pad, EG6180 background. Obtained by crossing INK438 to INK680 |
| INK767 | 3xFLAG::KSS-1 *(Ctr-rpl-36p)*; heat-shock inducible IDR-mCherry-IDR | *abuSi53[rpl-36p::3xFLAG::kss-1::rpl-36 3' UTR + HygR(+), abuSi33] I; abuSi36[hsp-16.11::KLMT-1(p.M1_D69)::mCherry::KLMT-1(p.A460_S503)tbb-2 3' UTR + HygR(+); abuSi9] IV* | Fig. 2H | Heat-shock inducible expression of mCherry fused with N- and C-IDRs |
| INK777 | 3xFLAG::KSS-1(INDDTL) *(Ctr-rpl-36p)* | *abuSi69[rpl-36p::3xFLAG::kss-1(R323_V328delinsINDDTL)::rpl-36 3' UTR + HygR(+), abuSi33] I* | S14B | Ribosomal promoter driven expression of N-terminal 3xFLAG tagged KSS-1 mutant (323RRGGTV>INDDTL), Chr. I landing pad, EG6180 background |
| INK783 | 3xFLAG::KSS-1(INDDTL) *(Ctr-rpl-36p)*; QX2343 | *abuSi69[rpl-36p::3xFLAG::kss-1(R323_V328delinsINDDTL)::rpl-36 3' UTR + HygR(+), abuSi33] I; qqIR47* | S14B | Ribosomal promoter driven expression of N-terminal 3xFLAG tagged KSS-1 mutant (323RRGGTV>INDDTL), Chr. I landing pad, Qx2343 background. Obtained by crossing INK777 to QX2343 |
| INK785 | 3xFLAG::KSS-1 *(Ctr-rpl-36p)*; KLMT-1::mNeonGreen; *kss-1(-)* | *abuSi53[rpl-36p::3xFLAG::kss-1::rpl-36 3' UTR + HygR(+), abuSi33] I; kss-1(abu125[p.R7Ffs*28]) V; klmt-1(abu107[klmt-1::2xTy1::mNeonGreen]) V; qqIR47* | S8B | Heat-shock inducible expression of KSS-1 (Chr. I landing pad), endogenous kss-1 knockout and C-terminal mNeonGreen tagged KLMT-1 (with 2xTy1 linker), Qx2343 background. Obtained by crossing INK563 to INK206 |
| INK793 | *kss-3.4(-)* | *kss-3.4(abu411[p.W100*]) II; fars-3(abu397[fars-3::mScarlet]) II* | Fig. 4A | *kss-3.4* knockout, C-terminal mScarlet tagged FARS-3, EG6180 background |
| INK801 | *kss-3.3(-) kss-3.4(-)* | *kss-3.3(abu416[p.S99*]) II; kss-3.4(abu411[p.W100*]) II; fars-3(abu397[fars-3::mScarlet]) II* | Fig. 4A | *kss-3.3* and *kss-3.4* knockout, C-terminal mScarlet tagged FARS-3, EG6180 background |
| INK806 | 3xFLAG::KSS-1(INDDTL) *(Ctr-rpl-36p)*; heat-shock inducible IDR-mCherry-IDR | *abuSi69[rpl-36p::3xFLAG::kss-1(R323_V328delinsINDDTL)::rpl-36 3' UTR + HygR(+), abuSi33] I; abuSi36[hsp-16.11::KLMT-1(p.M1_D69)::mCherry::KLMT-1(p.A460_S503)tbb-2 3' UTR + HygR(+); abuSi9] IV* | fig. S14B | Ribosomal promoter driven expression of N-terminal 3xFLAG tagged KSS-1 mutant (323RRGGTV>INDDTL), Chr. I landing pad, and heat-shock inducible expression of mCherry fused with N- and C-IDRs |
| INK808 | 3xFLAG::KSS-1 *(Ctr-rpl-36p)*; heat-shock inducible mCherry | *abuSi53[rpl-36p::3xFLAG::kss-1::rpl-36 3' UTR + HygR(+), abuSi33] I; abuSi11[hsp-16.11::mCherry::tbb-2 3' UTR + HygR(+); abuSi9] IV* | Fig. 2H | Ribosomal promoter driven expression of N-terminal 3xFLAG tagged KSS-1, Chr. I landing pad, heat-shock inducible expression of mCherry, Chr. IV landing pad, EG6180 background. Obtained by crossing INK563 to INK310 |
| INK854 | heat-shock inducible mCherry-IDR | *abuSi71[hsp-16.11::mCherry::KLMT-1(p.A460_S503)::tbb-2 3' UTR + HygR(+); abuSi9] IV* | Fig. 2I | Heat-shock inducible expression of mCherry fused with C-IDR of KLMT-1 |
| INK874 | heat-shock inducible IDR-mCherry | *abuSi82[hsp-16.11p::KLMT-1(p.M1_D69)::mCherry::tbb-2 3' UTR + HygR(+)] IV* | Fig. 2I | Heat-shock inducible expression of mCherry fused with N-IDR of KLMT-1 |
| INK877 | heat-shock inducible KSS-3.1 | *abuSi85[hsp-16.11::kss-3.1::tbb-2 3' UTR + HygR(+), abuSi33] I* | - | Heat-shock inducible expression of KSS-3.1, Chr. I landing pad, EG6180 background. Used for getting INK996 |
| INK880 | heat-shock inducible KSS-3.3 | *abuSi88[hsp-16.11::kss-3.3::tbb-2 3' UTR + HygR(+), abuSi33] I* | - | Heat-shock inducible expression of KSS-3.3, Chr. I landing pad, EG6180 background. Used for getting INK1011 |
| INK892 | 3xFLAG::KSS-1 *(Ctr-rpl-36p)*; heat-shock inducible mCherry-IDR | *abuSi53[rpl-36p::3xFLAG::kss-1::rpl-36 3' UTR + HygR(+), abuSi33] I; abuSi71[hsp-16.11::mCherry::KLMT-1(p.A460_S503)::tbb-2 3' UTR + HygR(+); abuSi9] IV* | Fig. 2I | Ribosomal promoter driven expression of N-terminal 3xFLAG tagged KSS-1, Chr. I landing pad, heat-shock inducible expression of mCherry fused with C-IDR. Obtained by crossing INK563 to INK854 |
| INK894 | 3xFLAG::KSS-1 *(Ctr-rpl-36p)*; heat-shock inducible IDR-mCherry | *abuSi53[rpl-36p::3xFLAG::kss-1::rpl-36 3' UTR + HygR(+), abuSi33] I; abuSi82[hsp-16.11p::KLMT-1(p.M1_D69)::mCherry::tbb-2 3' UTR + HygR(+)] IV* | Fig. 2I | Ribosomal promoter driven expression of N-terminal 3xFLAG tagged KSS-1, Chr. I landing pad, heat-shock inducible expression of mCherry fused with N-IDR. Obtained by crossing INK563 to INK874 |
| INK950 | *kss-3.1-3.4(-)*; FARS-3::mScarlet | *kss-3.1-3.2(abu490[g.11323197_11326206del]) II; kss-3.3(abu416[p.S99*]) II; kss-3.4(abu411[p.W100*]) II; fars-3(abu397[fars-3::mScarlet]) II* | Fig. 4A | Knockout of *kss-3.1*, *kss-3.2*, *kss-3.3*, *kss-3.4*, C-terminal mScarlet tagged FARS-3, EG6180 background |
| INK951 | *kss-3.1(-); kss-3.2(-)* | *kss-3.1-3.2(abu492[g.11323197_11326206del]) II; fars-3(abu397[fars-3::mScarlet]) II* | Fig. 4A | Knockout of *kss-3.1*, *kss-3.2*, C-terminal mScarlet tagged FARS-3, EG6180 background |
| INK957 | *kss-3.2(-); kss-3.3(-); kss-3.4(-)* | *kss-3.2(abu498[p.W47*]) II; kss-3.3(abu416[p.S99*]) II; kss-3.4(abu411[p.W100*]) II; fars-3(abu397[fars-3::mScarlet]) II* | Fig. 4A | Knockout of *kss-3.2*, *kss-3.3*, *kss-3.4*, C-terminal mScarlet tagged FARS-3, EG6180 background |
| INK976 | 3xFLAG::KSS-2 *(Ctr-rpl-36p)* | *abuSi96[rpl-36p::3xFLAG::kss-2::rpl-36 3' UTR + HygR(+), abuSi33] I* | fig. S10D | Ribosomal promoter driven expression of N-terminal 3xFLAG tagged KSS-2, Chr. I landing pad, EG6180 background |
| INK996 | heat-shock inducible KSS-3.1; *kss-3.1-3.4(-)* | *abuSi85[hsp-16.11::kss-3.1::tbb-2 3' UTR + HygR(+), abuSi33] I; kss-3.1-3.2(abu490[g.11323197_11326206del]) II; kss-3.3(abu416[p.S99*]) II; kss-3.4(abu411[p.W100*]) II; fars-3(abu397[fars-3::mScarlet]) II* | Fig. 4C | Heat-shock inducible expression of KSS-3.1, Chr. I landing pad, knockout of *kss-3.1, kss-3.2, kss-3.3, kss-3.4*, C-terminal mScarlet tagged FARS-3, EG6180 background. Obtained by crossing INK877 to INK950 |
| INK1011 | heat-shock inducible KSS-3.3; *kss-3.1-3.4(-)* | *abuSi88[hsp-16.11::kss-3.3::tbb-2 3' UTR + HygR(+), abuSi33] I; kss-3.1-3.2(abu490[g.11323197_11326206del]) II; kss-3.3(abu416[p.S99*]) II; kss-3.4(abu411[p.W100*]) II; fars-3(abu397[fars-3::mScarlet]) II* | Fig. 4C | Heat-shock inducible expression of KSS-3.3, Chr. I landing pad, knockout of *kss-3.1, kss-3.2, kss-3.3, kss-3.4*, C-terminal mScarlet tagged FARS-3, EG6180 background. Obtained by crossing INK880 to INK950 |
| INK1030 | heat-shock inducible PZL-1::mCherry | *abuSi124[hsp-16.11p::pzl-1::mCherry::tbb-2 3' UTR + HygR(+)] IV* | Fig. 3G | Heat-shock inducible expression of PZL-1 tagged with mCherry on C-terminus, Chr. IV landing pad, EG6180 background |
| INK1053 | 3xFLAG::KSS-2 *(Ctr-rpl-36p)*; heat-shock inducible PZL-1::mCherry | *abuSi96[rpl-36p::3xFLAG::kss-2::rpl-36 3' UTR + HygR(+), abuSi33] I; abuSi124[hsp-16.11p::pzl-1::mCherry::tbb-2 3' UTR + HygR(+)] IV* | Fig. 3G | Ribosomal promoter driven expression of N-terminal 3xFLAG tagged KSS-2, Chr. I landing pad, heat-shock inducible expression of PZL-1 tagged with mCherry on C-terminus, Chr. IV landing pad, EG6180 background. Obtained by crossing INK976 to INK1030 |
| INK1091 | EMS-2 | *EMS mutagenesis line 2* | Fig. 4C | Revertant line derived from EMS-induced mutagenesis of INK950, EG6180 background |
| INK1092 | EMS-3 | *hyde-1(abu529[p.E277K]) II; kss-3.1-3.2(abu490[g.11323197_11326206del]) II; kss-3.3(abu416[p.S99*]) II; kss-3.4(abu411[p.W100*]) II; fars-3(abu397[fars-3::mScarlet]) II* | Fig. 4C | Revertant line derived from EMS-induced mutagenesis of INK950, EG6180 background |
| INK1095 | EMS-6 | *hyde-1(abu532[p.P286L]) II, kss-3.1-3.2(abu490[g.11323197_11326206del]) II; kss-3.3(abu416[p.S99*]) II; kss-3.4(abu411[p.W100*]) II; fars-3(abu397[fars-3::mScarlet]) II* | Fig. 4C | Revertant line derived from EMS-induced mutagenesis of INK950, EG6180 background |
| INK1097 | EMS-8 | *EMS mutagenesis line 8* | Fig. 4C | Revertant line derived from EMS-induced mutagenesis of INK950, EG6180 background |
| INK1100 | EMS-11 | *EMS mutagenesis line 11* | Fig. 4C | Revertant line derived from EMS-induced mutagenesis of INK950, EG6180 background |
| INK1088 | mScarlet::ZYG-9.2 | *zyg-9.2(abu458[mScarlet:zyg-9.2]) II* | - | N-terminal mScarlet tagged ZYG-9.2, EG6180 background. Used as a background for INK1164, INK1169, INK1172 and INK1196 |
| INK1164 | *hyde-1(-) kss-3.1-3.4(-)* | *hyde-1(abu517[P203_L205delinsTN*]) II; kss-3.1-3.2(abu490[g.11323197_11326206del]) II; kss-3.3(abu416[p.S99*]) II; kss-3.4(abu411[p.W100*]) II; fars-3(abu397[fars-3::mScarlet]) II* | Fig. 4C | *kss-3.1, kss-3.2, kss-3.4* knockout, N-terminal mScarlet tagged ZYG-9.2, EG6180 background |
| INK1169 | *kss-3.1(-) kss-3.2(-) kss-3.3(-)* | *zyg-9.2(abu458[mScarlet:zyg-9.2]) II; kss-3.1-3.2(abu522[g.11323197_11326206del]) II; kss-3.3(abu462[p.S99*]) II* | Fig. 4A | *kss-3.1, kss-3.2, kss-3.3* knockout, N-terminal mScarlet tagged ZYG-9.2, EG6180 background |
| INK1172 | *kss-3.1(-) kss-3.2(-) kss-3.4(-)* | *kss-3.1-3.2(abu525[g.11323197_11326206del]) II; kss-3.4(abu460[p.W100*]) II;zyg-9.2(abu458[mScarlet:zyg-9.2]) II* | Fig. 4A | Knockout of *hyde-1,* knockout of *kss-3.1 to kss-3.4,* C-terminal mScarlet tagged FARS-3, EG6180 background |
| INK1196 | *kss-3.1(-) kss-3.4(-)* | *kss-3.1(abu546[g.11323196_11324454del]) II; kss-3.4(abu460[p.W100*]) II; zyg-9.2(abu458[mScarlet:zyg-9.2]) II* | Fig. 4A | *kss-3.1* and *kss-3.4* knockout, N-terminal mScarlet tagged ZYG-9.2, EG6180 background |

**Table S4.** List of gRNAs and primers used in this study.

| Lines generated | Modification | gRNA sequence 5'-3' | Repair template sequence | Genotyping primer - 1 | Genotyping primer - 2 | Amplicon size, bp |
| --- | --- | --- | --- | --- | --- | --- |
| INK303 | *klmt-1* knock-out | CCGTCTCTCCGTGCTTCTTG | no | TGCAGCTTGAAAATGATGAAA | CGATTACCGGGTAGACAGGA | 707 |
| INK63 | *3xFLAG::klmt-1* | CTGATTTTCGGACATTTCTC | tttgttttttccgtggtttttctttgctattttcgaatttaaattaaaaatacggctaataatctcaatattccagagaaatggattacaaagaccatgatggtgactataaggatcatgatattgactataaagaccatgactccgaaaatcagcgactttcgaacaacggttctgtagaag | ATCGCATGCGCCTTAAATACCG | TTTTCGTATTCGTCCGGAGCCT | 605 |
| INK66 | *klmt-1::3xFLAG* | GAATGTATTTAACTTTCCAT | tcagtgacagtgagacggagaaagccgccgacgaacaatccgtggagaaaaagagaaactcctcgtcacagaatccaatggaaagtgattacaaagaccatgatggtgactataaggatcatgatattgactataaagaccatgactaaatacattctttcccccctttccccgcttcatcaatg | ACTGCCAAAGTGCTCACCCTAA | GGGCTTAGGCGGCAAATTAAAT | 542 |
| INK169 | *klmt-1::mNeonGreen* | GAATGTATTTAACTTTCCAT | acgactcactatagggcgaattggccaagaactccttttcgacctcaaacacgtcacctttgatttgaagaatgaaaaaccaactcctagaattttcatcttgagaaaatttggtttgaaaataaaatacggatttcaggagccaccaaacatcacaaattttaatcggatctcctcaacgctttcgaagcttctggaactcgatgaaactaatccggacagtgtcttctcgagaaaacactcttccgaactgaagaaagtgactgccaaagtgctcaccctaatgaaaggagaagtcaaaagagctgagaagaatttgttaatgattcagagcaaagttggattggagatgaggtattttcgcacttcgcaacaaaacagcttcattttcgattttttcagtgacagtgagacggagaaagccgccgacgaacaatccgtggagaaaaagagaaactcctcgtcacagaatccaatggaaagtggtaccgtctccaagggagaggaggacaacatggcctccctcccagccacccacgagctccacatcttcggatccatcaacggagtcgacttcgacatggtcggacaaggaaccggaaacccaaacgacggatacgaggagctcaacctcaagtccaccaaggtaagtttaaacatatatatactaactaaccctgattatttaaattttcagggagacctccaattctccccatggatcctcgtcccacacatcggatacggattccaccaatacctcccatacccagacggaatgtccccattccaagccgccatggtcgacggatccggataccaagtccaccgtaccatgcaattcgaggacggagcctccctcaccgtcaactaccgttacacctacgagggatcccacatcaaggtaagtttaaacagttcggtactaactaaccatacatatttaaattttcagggagaggcccaagtcaagggaaccggattcccagccgacggaccagtcatgaccaactccctcaccgccgccgactggtgccgttccaagaagacctacccaaacgacaaggtaagtttaaacatgattttactaactaactaatctgatttaaattttcagaccatcatctccaccttcaagtggtcctacaccaccggaaacggaaagcgttaccgttccaccgcccgtaccacctacaccttcgccaagccaatggccgccaactacctcaagaaccaaccaatgtacgtcttccgtaagaccgagctcaagcactccaagaccgagctcaacttcaaggagtggcaaaaggccttcaccgacgtcatgggaatggacgagctctacaagtaaatacattctttcccccctttccccgcttcatcaatgtttcgaagaaaacgtttcattgtgttgtttgtgatgagatttggtttgtttttcagtgccactctttcagtggcaagtatataaattgtaatgtttttatttaaaaattatctctatatttatatttgccaaaaaatctggagtttaactgaaaaaacgcgcgtaaactccagatttttcttttttttttgacctgaaaatttaatttgccgcctaagccccccttcccctcacttttctcgattttcccccatttcccctttgttcccactcaattttcctgattcgaaaccctaatccgaaatagcacggggattacgttatcgagtctcttttttgaactcggcgcatagagctcagttcaaaaattatgccatgttttggataactttgcgttttttgattcccccaccctttcatcgtcatcccgtcatttttccagacgcggcgcaacgatttgagagtattgagcgagaaaaggaagaaaaatgctccgggaatcccaatggcttccgtggggcgtggtcaaaaaccacccgaaatgagctgttccctttagtgagggttaattgc | ACTGCCAAAGTGCTCACCCTAA | GGGCTTAGGCGGCAAATTAAAT | 1346 |
|  | *klmt-1::2xTy1::mNeonGreen* | GAGACGGTACCACTTTCCAT | gccgacgaacaatccgtggagaaaaagagaaactcctcgtcacagaatccaatggaaagtggtgaagtgcataccaatcaggacccgctggatgaagtccacacaaaccaagatccactcgatggtaccgtctccaagggagaggaggacaacatggcctccctcccagccacccacgag | ACTGCCAAAGTGCTCACCCTAA | GGGCTTAGGCGGCAAATTAAAT | 1409 |
| INK277 | *ΔN-klmt-1::3xFLAG* (p.S2_T45del) | CTGATTTTCGGACATTTCTC + CCTTCCTCCAACTACTCCAG | gttttttccgtggtttttctttgctattttcgaatttaaattaaaaatacggctaataatctcaatattccagagaaatgactccagtggtacgccattacgtttcttttattttttttatattaaaaaataacaatttaaacgacttttcgccagaaaaaccacattttcg | ATCGCATGCGCCTTAAATACCG | GATCCTTTTCCACCGCCGTTTT | 768 |
| INK581 | *3xFLAG::klmt-1-ΔC* (p.A460_S503del) | GCAAAGTTGGATTGGAGATG + GAATGTATTTAACTTTCCAT | ctgaagaaagtgactgccaaagtgctcaccctaatgaaaggagaagtcaaaagagctgagaagaatttgttaatgattcagagcaaagtttaaatacattctttcccccctttccccgcttcatcaatgtttcgaagaaaacgtttcattgtgttgtttgtgatgagatttggtttg | ACTGCCAAAGTGCTCACCCTAA | GGGCTTAGGCGGCAAATTAAAT | 302 |
| INK422 | *kss-1* knock-out | GAAAAATAGATTCACTACGA | no | ACGGTGTTTCAGGATAGACGAGA | TGTGCCGCCTCTCCTGAATTTA | 852 |
| INK206 | *kss-1* knock-out | ACGTCTGGGAGTCGAATAAG | no | TATCCCATCTGCCACGTGTTGA | TACCGATGAAAGAGGTGTCGCC | 306 |
| INK444 | *3xFLAG::kss-1* | ACGTCTGGGAGTCGAATCAA | cgaacaaccaccaattattgatttttagatctcaataaatctttggattttaatatggactacaaagatcatgacggtgactataaagatcatgacatcgattacaaggatgacgatgacaaggagtttccattgattcgactcccagacgttccaaaaattcaaattatccaattgatgtctattgg | TATCCCATCTGCCACGTGTTGA | TACCGATGAAAGAGGTGTCGCC | 415 |
| INK324 | *pzl-1* knock-out | ACATGGGTTGCCACGGAACT | ctgcaaaagataattcggaacgacgccaacatattttgtcaagtgttggcaatcagaagcgttacatgagtaactgagtcgactcggcgcggaattctccaaattctctgtttctcttctaccagatctgttggaaaagatgaaagag | AACAATGAAATGGCAGGAGCGG | TCTCTCCAATTGCATCGCCTCA | 434 |
| INK485 | *kss-2* knock-out | GAAGATGGAGTATAGTAAGC | ggagacgacgagtggagagtgtttctgggtccttcgaagatggagtaagtagctagccggataatatctcagaggaggatgtgaaaccatggtgagttcc | ATTCGACTGCCAGATGTTCCCA | *TTGCGACACAGGGATAACTCAGT* | 521 |
| INK505 | *fars-3::3xFLAG* | ATAACAATAATTTAGAGGAA | cctctttcggactcacccttccatgtggagccgtggaattcaacgtcgaaccgttcttggattacaaagaccatgatggtgactataaggatcatgatattgactataaagatgacgatgacaagtaaattattgttattgaatgttttctatggaattgttgtttttaaatttaaatattcattcc | ATTCCATTGATCTCACCCCCTGG | TTCTGGCTGGGAAAAAGGGTTCT | 711 |
| INK577 | *pzl-1::3xFLAG* | CGGAGGACGACTAATTTATTA | cctcatgttgccggttgccgccttcgagatcaaaatctttacggaggacgactacaaagaccatgatggtgactataaggatcatgatattgactataaagatgacgatgacaagtaatttattattttttaattcgttattcaattcattattttgacccccacaacc | AACCTTCTCGCCTGGACACTAC | ATCGATGGAGGCTTGGAAGGAG | 427 |
| INK590 | *kss-2* (p.G297R) | AACCAACTTTTGATGACGGT | ccagagcatcgaatacaactgagaaccaacttttgatgaccgtgagattggtatcttatcctgccgagccaggacgacgagagctcgagttctacg | TGTTCTCAAGGGTTTCCGATCCA | ACCCTCTTCTCATCCAGCCAAC | 334 |
| INK722 | *fars-3::mScarlet* | ATAACAATAATTTAGAGGAA | cccttccatgtggagccgtggaattcaacgtggaaccgttcctcggaggtggatcaatggtctcgaaaggagaggccgtcatcaaggagttcatgcgtttcaaggtccacatggagggatccatgaacggacacgagttcgagatcgagggagagggagagggacgtccatacgagggaacccaaaccgccaagctcaaggtcaccaaggtaagtttaaacatatatatactaactaaccctgattatttaaattttcagggaggaccactcccattctcctgggacatcctctccccacaattcatgtacggatcccgtgccttcatcaagcacccagccgacatcccagactactacaagcaatccttcccagagggattcaagtgggagcgtgtcatgaacttcgaggacggaggagccgtcaccgtcacccaagacacctccctcgaggacggaaccctcatctacaaggtaagtttaaacagttcggtactaactaaccatacatatttaaattttcaggtcaagctccgtggaaccaacttcccaccagacggaccagtcatgcaaaagaagaccatgggatgggaggcctccaccgagcgtctctacccagaggacggagtcctcaagggagacatcaagatggccctccgtctcaaggacggaggacgttacctcgccgacttcaaggtaagtttaaacatgattttactaactaactaatctgatttaaattttcagaccacctacaaggccaagaagccagtccaaatgccaggagcctacaacgtcgaccgtaagctcgacatcacctcccacaacgaggactacaccgtcgtcgagcaatacgagcgttccgagggacgtcactccaccggaggaatggacgaactctacaagtaaattattgttattgaatgttttctatggaattgttgtttttaaatttaaatattc | ATTCCATTGATCTCACCCCCTGG | TTCTGGCTGGGAAAAAGGGTTCT | 1506 |
| INK793 | *kss-3.4* knock-out | GGATAAACGATGAGTGGACAATG | caacaaaattcagtaaaaaaatacggtgttttaggataaacgatgagtaagtgactaagcttagagaaaactttcaatgtgttcatacgtcttcaaaaagttcttcc | CGATGGGCAAAATAGTGCGGTT | GCTCAGCGAAACATTCTGCACT | 886 |
| INK801 | *kss-3.3* knock-out | AGGATAGACGAGAAGAGCGCA | caacaaaattcagtaaaagaatacatgatttcaggatagacgagaagtgactaagctttgatagagagaactttcaatgtgttcatacgtcttcaaaaagttcttcc | GAGAACTATCTGGCGTGGGTGT | AGTCAAGCTCGTTTAGTATTTGGGA | 1025 |
| INK951 | *kss-3.1; kss-3.2* double knock-out | ACTGTGCAATCCACATCGAACA + GAAACACAACAACATTCGAGTT | ccagaatcaaactggctctttcctctcgaaaaatggagaactacttgtcatgggtgttcaaaaatcactaaatgtcctgtttcgaaaagtgaaggtacgaacgagatcaaaggggttcgagatgtccgggagcc | TTTGCCCATCGAACGACTACCT | GTGGGAAGGGGATATGACTGAGC | 1126 |
| INK1088 | *mScarlet::zyg-9.2* | AGTCAGTCAAAAAATGTCGAAT | cctctttcacccctaaaaccctcttttttcagtcagtcaaaaaatggtctcgaaaggagaggccgtcatcaaggagttcatgcgtttcaaggtccacatggagggatccatgaacggacacgagttcgagatcgagggagagggagagggacgtccatacgagggaacccaaaccgccaagctcaaggtcaccaaggtaagtttaaacatatatatactaactaaccctgattatttaaattttcagggaggaccactcccattctcctgggacatcctctccccacaattcatgtacggatcccgtgccttcatcaagcacccagccgacatcccagactactacaagcaatccttcccagagggattcaagtgggagcgtgtcatgaacttcgaggacggaggagccgtcaccgtcacccaagacacctccctcgaggacggaaccctcatctacaaggtaagtttaaacagttcggtactaactaaccatacatatttaaattttcaggtcaagctccgtggaaccaacttcccaccagacggaccagtcatgcaaaagaagaccatgggatgggaggcctccaccgagcgtctctacccagaggacggagtcctcaagggagacatcaagatggccctccgtctcaaggacggaggacgttacctcgccgacttcaaggtaagtttaaacatgattttactaactaactaatctgatttaaattttcagaccacctacaaggccaagaagccagtccaaatgccaggagcctacaacgtcgaccgtaagctcgacatcacctcccacaacgaggactacaccgtcgtcgagcaatacgagcgttccgagggacgtcactccaccggaggaatggacgagctttataagggtggatcgaattgggactatatagacgaagtggacattattccaaagcttcc | CGCTTCATTCGTTCATAAAAACCTGG | AGCCCTGTAGCGAATTTAGCGA | 1430 |
| INK1164 | *hyde-1* knock-out | TGCCCGCATTTCATTGCGATCT + CGAGCCATGCCACAGCTCCCT | cgactgagatgatgcagtacttcagcggcttctctaataatttccgagccatgactaactgaaatgcgggcacaaagtaattgtgacgtggcccactcctcctggagcgc | GTAGTGACCCGACAGAAGCTGAT | TGAAAAAGGAGCTTTGAAGGCCG | 599 |
| INK1193 | *kss-3.1* knock-out | ACAATCTATCTGCTGAGTGAAT + GAAACACAACAACATTCGAGTT | gtcatttgagattatgacgacaagacagataccatctgccacgtcatttcacaatctatctgcaaatgtcctgtttcgaaaagtgaaggtacgaacgagatcaaaggggttcgagatgtccgggagcc | ACGGAGGTTAAGTTTTTAGCGCTT | GTGGGAAGGGGATATGACTGAGC | 1106 |

**Table S5. List of repair templates and primers used for synthetic landing pad injections.** Same gRNA was used for all SLP injections: 5’-AAAGTCCACAATCTCCACGT-3’. Amplicon size for empty Chr. I SLP - 2007bp. For empty Chr. IV SLP - 2236bp.

| Lines generated | Insertion (short name) | Transgene inserted | Background | Repair template sequence (plasmid) | Genotyping primer - 1 | Genotyping primer - 2 | Genotyping primer - 3 | Amplicon size 1+2 (external), bp | Amplicon size 1+3 (internal), bp |
| --- | --- | --- | --- | --- | --- | --- | --- | --- | --- |
| INK310 | heat-shock inducible mCherry | *hsp-16.11p::mCherry::tbb-2 3' UTR + HygR(+)* | Chr. IV SLP | pAB0174 | TCCAATCTCGCTCTTCAACTCGT | TGTTCGCCGTACAGAGAACATCT | TTTTGCGGTTTGTGTTCCCT | 4966 | 1591 |
| INK318 | heat-shock inducible KLMT-1 | *hsp-16.11p::klmt-1::tbb-2 3' UTR + HygR(+)* | Chr. IV SLP | pAB0176 | TCCAATCTCGCTCTTCAACTCGT | TGTTCGCCGTACAGAGAACATCT | TGGAGGAAGGTCTGGAGCATTG | 5614 | 1573 |
| INK336 | heat-shock inducible IDR-mCherry-IDR | *hsp-16.11p::KLMT-1(p.M1_D69)::mCherry::KLMT-1(p.A460_S503)::tbb-2 3' UTR + HygR(+)* | Chr. IV SLP | pAB0192 | TCCAATCTCGCTCTTCAACTCGT | TGTTCGCCGTACAGAGAACATCT | TGGAGGAAGGTCTGGAGCATTG | 5305 | 1573 |
| INK438 | heat-shock inducible KSS-1 | *hsp-16.11p::kss-1::tbb-2 3' UTR + HygR(+)* | Chr. I SLP | pAB0260 | TGAGATGATTGATGAGGCGTCAAG | TGAAGAGAAAAAGGGCATGGTCA | TACCGATGAAAGAGGTGTCGCC | 5285 | 1588 |
| INK563 | 3xFLAG::KSS-1 *(Ctr-rpl-36p)* | *rpl-36p::3xFLAG::kss-1::rpl-36 3' UTR + HygR(+)* | Chr. I SLP | pAB0293 | TGAGATGATTGATGAGGCGTCAAG | TGAAGAGAAAAAGGGCATGGTCA | TACCGATGAAAGAGGTGTCGCC | 4864 | 1419 |
| INK586 | heat-shock inducible KSS-2 | *hsp-16.11p::kss-2::tbb-2 3' UTR + HygR(+)* | Chr. I SLP | pAB0294 | TGAGATGATTGATGAGGCGTCAAG | TGAAGAGAAAAAGGGCATGGTCA | TTGCGACACAGGGATAACTCAGT | 5955 | 1881 |
| INK680 | heat-shock inducible PZL-1 | *hsp-16.11p::pzl-1::tbb-2 3' UTR + HygR(+)* | Chr. IV SLP | pAB0336 | TCCAATCTCGCTCTTCAACTCGT | TGTTCGCCGTACAGAGAACATCT | ACTGTGTTCAGCTACGATTCCGA | 6469 | 2062 |
| INK777 | 3xFLAG::KSS-1(INDDTL) *(Ctr-rpl-36p)* | *rpl-36p::3xFLAG::kss-1(R323_V328delinsINDDTL)::rpl-36 3' UTR + HygR(+)* | Chr. I SLP | pAB0332 | TGAGATGATTGATGAGGCGTCAAG | TGAAGAGAAAAAGGGCATGGTCA | TACCGATGAAAGAGGTGTCGCC | 4864 | 1419 |
| INK854 | heat-shock inducible mCherry-IDR | *hsp-16.11p::mCherry::KLMT-1(p.A460_S503)::tbb-2 3' UTR + HygR(+)* | Chr. IV SLP | pAB0382 | TCCAATCTCGCTCTTCAACTCGT | TGTTCGCCGTACAGAGAACATCT | TTTTGCGGTTTGTGTTCCCT | 5098 | 1591 |
| INK874 | heat-shock inducible IDR-mCherry | *hsp::KLMT-1(p.M1_D69)::mCherry::tbb-2 3' UTR + HygR(+)* | Chr. IV SLP | pAB0381 | TCCAATCTCGCTCTTCAACTCGT | TGTTCGCCGTACAGAGAACATCT | TGGAGGAAGGTCTGGAGCATTG | 5173 | 1573 |
| INK877 | heat-shock inducible KSS-3.1 | *hsp-16.11p::kss-3.1::tbb-2 3' UTR + HygR(+)* | Chr. I SLP | pAB0346 | TGAGATGATTGATGAGGCGTCAAG | TGAAGAGAAAAAGGGCATGGTCA | TGGGGAGATCTGGCAGTTGGA | 5457 | 1384 |
| INK880 | heat-shock inducible KSS-3.3 | *hsp-16.11p::kss-3.3::tbb-2 3' UTR + HygR(+)* | Chr. I SLP | pAB0347 | TGAGATGATTGATGAGGCGTCAAG | TGAAGAGAAAAAGGGCATGGTCA | ACACCTGTCAAGTTCGCGAC | 5286 | 1891 |
| INK976 | 3xFLAG::KSS-2 *(Ctr-rpl-36p)* | *rpl-36p::3xFLAG::kss-2::rpl-36 3' UTR + HygR(+)* | Chr. I SLP | pAB0372 | TGAGATGATTGATGAGGCGTCAAG | TGAAGAGAAAAAGGGCATGGTCA | TTGCGACACAGGGATAACTCAGT | 5537 | 1715 |
| INK1030 | heat-shock inducible PZL-1::mCherry | *hsp-16.11p::pzl-1::mCherry::tbb-2 3' UTR + HygR(+)* | Chr. IV SLP | pAB0407 | TCCAATCTCGCTCTTCAACTCGT | TGTTCGCCGTACAGAGAACATCT | ACTGTGTTCAGCTACGATTCCGA | 7342 | 2062 |

**Other Supplementary Materials (as separate files)**

**Data S1.** Raw data and summary for all genetic crosses in this study.

**Data S2.** List of candidate TA genes within the Chr. V NIL introgression strain.

**Data S3.** *In vivo* interactome of KSS-1 revealed by IP mass spectrometry.

**Data S4.** List of candidate TA genes within the Chr. II NIL introgression strain.

**Data S5.** *In vivo* interactome of KSS-2 revealed by IP mass spectrometry.

**Data S6.** EMS-derived mutations identified in suppressor lines.

**Data S7.** Sequences of the 560 KSL proteins identified.

**Data S8.** *C. elegans* proteins with predicted structural homology to KSS-1.

**Data S9.** Sequences of toxins and antidotes.
