## Supplementary material for "A regulatory module driving the recurrent evolution of irreducible molecular complexes": Data_S9

**Data S9. DNA and protein sequences of toxins and antidotes identified in this study.** For genomic sequences exons are marked in uppercase. EG-PZL-1 and NIC-HYDE-1 don’t act as toxins and EG-KSS-1 does not protect from KLMT-1 toxicity. Their protein sequences can be found at the end of the list.

>klmt-1

ATGTCCGAAAATCAGCGACTTTCGAACAACGGTTCTGTAGAAGAATTTTATACGgtaaaaatggtttttttaaaaagacaaagtgcttttttgagatagttctaagcggttttatgcgaaaacataatatttgcggcgattattaggttgaaaaaacaaattttccatggaaaaatgctattttcaagaaaaaaaatttgtttcttcttttccagcgaataccaacagaaaagtagcaattttaatggcgaaaaccattattttctataatttttcacctgaaaatattaaattgtcccgtttgctaacccattttgcatttgaaaaacataaaatcccattcgaaatctcaaaaatatttattttcagGCTCCGGACGAATACGAAAAGACAACCACTGCATCTAAGCAGTTGCCGCCAAGTTTCAATGCTCCAGACCTTCCTCCAACTACTCCAGTGgtacgccattacgtttcttttattttttttatattaaaaaataacaatttaaacgacttttcgccagaaaaaccacattttcgacataaaataactaaaaataattttttgacatgaaaatagttttttaaataaaaaaatacgattttcagactaaaaaaatcatttccaatctgaaaattcctattttcccttgtccgaatctttttccatcttcaaaaacccatagtttgcaaaccacaattttccgagccgaaaactctacttcccgctgaaaaactgcattctcctcattaaaaaattttattttagctcaaaaaccctattttcgcccgagaaatcaccatttttctttttgaaaattctgtttcgacctaaaagacacccattttctttgaacattgaaatcctacttttccaaaactaaaaaccataattgcagcttgaaaatgatgaaaatgtggtttctccggacaaaaaaccatatcagtttctattttctatttttatggcaagaataacataattttctgttaatttttcgctgaataacccaatttttctttcagACATCCGAAAAAACGGCGGTGGAAAAGGATCGCAGAGAATCTATGGACAAGAAGCATATGGACAAGAAGCACGGAGAGACGGCGGTCGGCGAGGATCTCAACGATCAGGAGATTTACAAAATCGACAAACGATACGATTTGTTGAGTTCAGAGGGGCTCTCCAGAGCTATGAGAATCTTCAAGCAACACTTCGAATTCCCGGACTTTCGTTTCCTAGATGCAAAGTCGATTGAGAAGATAAATGTGGCTGCGAATGCATCGGTCTTCCACCCATATTTATCCGGTGCGATTGTTCGCCAGATTTCATTAGATTTGGAATCTCTCAGATTTTTGAATGGATTGGTAGCTGATAACCAGGAGACGGGCAAGAAAAGAATGCAATATTTCATCACAGCACAGGATTTGGACAAAATGAGTGGACCGTTTGAATATCGGGCCGAAGCTTCAAACGTAATCAAATTCCGTCCGTTAAATCAGACGAAGGAGCACACGGCGGATGAGCTGATGACGTTGTACTCATCTGACAGCCATATGACAGATTTCCTTCAACTCATTCAAAATCTTCCTGTCTACCCGGTAATCGCTGATAAGAATGGACTGATATCCTTGGTGCCAGTAGTCGATGCGGAACCTATGAAGATATCAAtcgagacgaagagtttgatgattatcgtaacatcagttgataaagagaacggcatccgaatcctcaacaacgtcctcgccgctatttcgattcgaattgaaaaaccacttgtcgttgagcccgtTTTGATTGAATACGAGAAATTCGAGGAACCGAGAGCCTTAGAGCTCTCCCCACCGCTCTCTTATCGCAAGATGACAGTGACGACGCCGGAGATCAACACGAAGATTGGATTGAATCTTCAAGATGAAGAGATGGCTAATCTACTAAATAATATGTCTCTGAAAGCGGAGGTCGCCTCGAAAGAAGTCCTCGAAGTGATGATACCGCCCACTCGACACGATATTCTACACGCGTGTGACATTTCAGAAGACGTCGGAGTCGCGTGTGCCTACAAAAATTTCCTAAGGGATTTGTCAAAGgttcgttttcatttttccacatttaaaactagaaatttttcaaaacttgttttttgaaccaaaacccctgtttttaccacccaaaatcacctttctccgctccaagaactccttttcgacctcaaacacgtcacctttgatttgaagaatgaaaaaccaactcctagaattttcatcttgagaaaatttggtttgaaaataaaatacggatttcagGAGCCACCAAACATCACAAATTTTAATCGGATCTCCTCAACGCTTTCGAAGCTTCTGGAACTCGATGAAACTAATCCGGACAGTGTCTTCTCGAGAAAACACTCTTCCGAACTGAAGAAAGTGACTGCCAAAGTGCTCACCCTAATGAAAGGAGAAGTCAAAAGAGCTGAGAAGAATTTGTTAATGATTCAGAGCAAAGTTGGATTGGAGATGAGgtattttcgcacttcgcaacaaaacagcttcattttcgattttttcagTGACAGTGAGACGGAGAAAGCCGCCGACGAACAATCCGTGGAGAAAAAGAGAAACTCCTCGTCACAGAATCCAATGGAAAGTTAA

>KLMT-1

MSENQRLSNNGSVEEFYTAPDEYEKTTTASKQLPPSFNAPDLPPTTPVTSEKTAVEKDRRESMDKKHMDKKHGETAVGEDLNDQEIYKIDKRYDLLSSEGLSRAMRIFKQHFEFPDFRFLDAKSIEKINVAANASVFHPYLSGAIVRQISLDLESLRFLNGLVADNQETGKKRMQYFITAQDLDKMSGPFEYRAEASNVIKFRPLNQTKEHTADELMTLYSSDSHMTDFLQLIQNLPVYPVIADKNGLISLVPVVDAEPMKISIETKSLMIIVTSVDKENGIRILNNVLAAISIRIEKPLVVEPVLIEYEKFEEPRALELSPPLSYRKMTVTTPEINTKIGLNLQDEEMANLLNNMSLKAEVASKEVLEVMIPPTRHDILHACDISEDVGVACAYKNFLRDLSKEPPNITNFNRISSTLSKLLELDETNPDSVFSRKHSSELKKVTAKVLTLMKGEVKRAEKNLLMIQSKVGLEMSDSETEKAADEQSVEKKRNSSSQNPMES

>kss-1

ATGGAATTCCCCCTTATTCGACTCCCAGACGTTCCAAAAATTCAAATTATCCAATTGATGTCTATTGGGGAGCGgtaagtactccgaaagggttttccaggacattcattaattttcagAATCAAACTGGCTCTTTCCTCTCGAAAAATGGAGAACTATCTGGCGTGGGTGTTCAAAAATCGCACTCTAGATGCGGTTTATACTATTGAATTGCGAGGCGACACCTCTTTCATCGGTATCGACACAAAAACATGGGTACTGTGTATGAACCATTTCAAACCAGAATGTAAAAAGCCGGGTTATATTTCAGTGGAAGAATTGAAACCGTGgttagttatctagagaaattcagtgaaagaatacggtgtttcagGATAGACGAGAAGTGGACAATGATGGAGAGAACTTTCAATGTGTTCATACGTCTTCAAAAGGTTCTTCCGTGTGTAATGACCAATTTATGCGTAACTCTGAAGACCGAACCGACGCCGATTCTAAAAATCCTAAGTTATCCGTGCGTCGCGAACTTGACAGGTGTGCATTTCTATGGAGGAACAGTAGAAAGAAACGAGTTGGATGCAATTATGGAATGGAGAAAGGAGAACACGATTCAGTATATTTCGGTCACGGATAACGAAATTCCATTGGATTATAGACATCCGAATgtgagttttcttttgaatagttccactttatcagaaaaaaatgtgcagaatattacatcaatgatttcagGCCTTTCGTTTCACTGGAGTAGTGTATGGTGATGCTCGATGGATAAGTCTCGAAGATTTACTCTCGATGAGAAACTTCTATCGACCGATTTTCGGGAAAAATAGATTCACTACGAAGGATCTGAACACATTTATTAAATATTGGATCAACTGTGATGAGAATATGTTCACTTATCTAGTAATTGATTACAAAGATATCAATATATCTGAATTGATGAAGGATATTACTGTTCTCAAGGGTTTCCGATCCGAAAAACCATTTTATCTGATgttagttgcttcttttttttatttagaggattattagcaataattaattgatcaaaagaaaaaagaaatagttttcagAGCATCGAATTCAACTGAGAATCAAATTCTATTGACAATTCAATTGATAGAACACACTGAACCTCAAATACGAAATGTGCTCGACTTCTCGATTCAGTATGCTAATGAACCACATAAATTCAGGAGAGGCGGCACAGTGCCTGCATGGACTCGAGAATACCGTATTCTTGGCCGATTGGCGAGAAAACGTTGGTTGGAAGAGGAGATGATTACTGTAGGAATATCGATTGAAGAAATAAAAGAACTCGATGATTTGGAGGAGGAGTTGACTGTAGAGGAAGTTCGGATCATTGATGGAATAATGACTGTTCAAGAGAAACTGTAG

>KSS-1

MEFPLIRLPDVPKIQIIQLMSIGERIKLALSSRKMENYLAWVFKNRTLDAVYTIELRGDTSFIGIDTKTWVLCMNHFKPECKKPGYISVEELKPWIDEKWTMMERTFNVFIRLQKVLPCVMTNLCVTLKTEPTPILKILSYPCVANLTGVHFYGGTVERNELDAIMEWRKENTIQYISVTDNEIPLDYRHPNAFRFTGVVYGDARWISLEDLLSMRNFYRPIFGKNRFTTKDLNTFIKYWINCDENMFTYLVIDYKDINISELMKDITVLKGFRSEKPFYLIASNSTENQILLTIQLIEHTEPQIRNVLDFSIQYANEPHKFRRGGTVPAWTREYRILGRLARKRWLEEEMITVGISIEEIKELDDLEEELTVEEVRIIDGIMTVQEKL

>pzl-1

ATGTGGTCATCGACAACTTCATCCCACCAAAATCAACAAAAAATTCATGAGAAAGTATATgtaagtgatcctttccaattccaatttcaaaaaaaactttgatttttcacgttaaaattttttctgccctacaccaggccctaaacaattgaaaaactcgactatcaatccaatttattttgttagaaacttatttctagcataataatatcctttctgataacaaacgaacaacattttttactgttttgaattaaaaactatctttcagccgcttatttgaaaatccgcttttctcagttttgaaagtcaaaaactattgttttttctccagtttttttccagtttgcaaacggaataccacttttagctgtaaaaacaattttttctaaaatctatcaaaatgttttcaaaaatttaccaaaacatatacaatttgcagAAAAACGAACATCAAGTCTGTCTTCCGCCATGTGAGCTGAAAGACGAGGAATATGAGCCGAAAAAGTCGTTCTACGAAATGCATCAGCAAACGAAGCGGTGGACGAATTGGGAATCGCAACGAATTGTAACTGCTCCGGGACACTGTGCAACAGTAGACAGTGTTCTTCTATTCGAAAAAAACGACAATAAATTCTGTCTCTCTGAAAGTCGTGATCGAGTGATTCGTTTTTGGGACGTCGATAACGTTGAACGAGGAGTTGATTCAGTGGCAAATCCATGGACAGTGGCACAAGATGATATGGCACAATTGGAATGGAGTTGGAATATGGCTCGTGATTCGAACACTGATAGATTCTACTCAACCTCATGGGATTCAACTGTTAAAAGCTGGGCGATCACCGACAACGGAGCCATTCAGAATCTGAATACAGTAAACGTCGGATCAGCAGTTCAAGCGGTTTCCTGTTCTGGAAATGAAAACGAAATCGTTTGTACTACTTTCGCAAAAAGAACAGCTGTAATTGATTCGAGAACTTTCGGAATCGTAGCTGAACACAGTCTTCATAAACGAGCGGTCATAGGATTAGCTGTCAAGGAGAATAAGATATTCACGAGTGGAGAGGATCGATTGATGATGATGGTTGATCGAAGGAATATGAGTAGACCTGTTCTTTTTgtaagtgaacatctttataatgaactacagtagttttcagGAATACTGTCAAGACGCTTACAAATCCTCTCTATCTCTTCAACGGAATCAATTGCTCACTTCAACTTCAGACGGAAAAGTCAAACTGTATGATGCGACGAATTTCCATGATTTTCAGACATATTCAgtgaggaatgaaacagttacagtaatccaatattctcttatcagGTCGGCGCCTTCACCCGTCAATCGCTTCTTCAACACGGAGCCCATTTCGTTATGGCTCGTTGTAAAAGAGGATATTATAAATTCCTTgtaagtgatcaactccctgaaaatgaacttgatccctcgagaaacacattttcacgatgcttcgaaagcatgtcacaagttctgatttttcagATGAACAGTCCTGGAATTCGGAGTCTAAAGAGATGTTCATCACATCAACTGACTGCTAAACCAGCTAAATTTGATTACTCgttagtttgccatgaataccgtaacccaaacattcatttttcagATCCGAAATGAAGACACTTACAATTGGAAATTCGATGGATAGAAGTGAAAGACGATCCATGATTgtaagtaagaaactccctaataataaaaaaaacgaacttgatcccttgagaaacatggaacactttgttgaaaaagcaatcatttataagaaaaatagcaatttcagaaaaataatatataacattttaaaagcattcgtcgctaatatcgctcatgctgtcgctttctccctcttttttgttcacatcaatatcgtcctcatcacttcccgacctcaccacagaaaatccttcattaagatttccctctttgttcatttcgtgaaagttaaattcttcctctttggcgaagttttttctctttggcactctattgtccgtatttggtttcagagtgtgctcaaatagttttcgtgcgaccttcgcgggaaaacctctcaagaatcgaagagttctatctctaactttaagaaaaattttttgaaatggaaaactcgaaatttatgtacaatagtttgctacccgtatttccttctcttctcaaataattcttcgttttagaccacactaattcaatagttgcaaaacacagtggtactgcggcaagcaaagaatacgaacacgacgcgtttggttaattcgtcgatctcgtagacagtgaacgccctgtttagcatttttcagaaaaacgggacgtgtgaaagcagagtaataaagaacattatgatcatctagaaactatttaatatcagcgttgctcctgatagaattcaaaggctgaaataagcatgaagctataatagaagaatctagattgatttttcggcacttattcgatatgttttcctgttttatccgcttccgactcaaacagaggcaactcttctcatagaaagatcataatttttcgcttttatgtcatggtggttattggttttctaatcttccacctttcaacccgacacaatgaattccgccgagtctttcggaattccatccagtgtgacaacagcaatgacgattcctagcctccctttttcttttccccttcttggacctgccatcaccgtttccatgtccaccaatcttgattttttataatttgtcaaatttttgcgttgctaaaatctttgtcattcctttatgtgcccatgtctcgtctaaatttccaaaatagggtgaacatttgtctcactcctctttctctcttccatacgcttgtctacgatatatcatagatttattgtcgattttcgatgttaaaggggtataccgctaattctctcgttgtgcacgcgaaaaattgaataaatacgtgaatcttttttgaagaattaaagagggccagtaggaaaagtggatataggctccgataaagtgccagaaaaacactgaatggtagaaaaaaataatgcacccatgagtcagttgataatataggtcatcaggtaacatatatgtcatttttgtcttctaattagtacagaaactactgaaaaccttgtttatgttttttagaacccgaaaattccagttgctctcactgacctacttatttgaccaactagttagctatcattatcttcatcctttcctggctccctttttcagctatttactctgatttcggctaaaacagtatgacctctatgttacatgatgacctatattgccagctgccttagaggtaacttcttttgtcctcctcgagattttataatgtttccagggcctttattcactttactaatcactattcatgtctcttttattcaacttcaaaaaagaattcacggggttattcatttttcgcgtgcacaacgaggacattttatactgtaaaatgcacaacgaaaccccttttttggcactttgagcgcttatatctcggttcagagacgtttatggaaaaagtgaagtatttactagagttcgatcacgctccgcgagctgaatcgaatcatataagacttaagaaaatttgagctaattttgaattagcgtattacccctttaatatgtcagaaatagaagggttttaatagttcacgattacatgacaagataaagttaggaatggaaatcaggctattttctttcttcttcctcctgtcggtatcccgtttgcctcttccatctttaaactggttcaattataataattcgatcctcatcttctgcttataccatattttttcttggtatcctccggtagtgtttcttcccttttttgaccctcatctttctccggggtcattctgctccatctccttgctgatttattttcgttcatgatatccccaatagttgtttgatctttttccatgtctgcttcgtcatagtcacgacactccttgcggccacaaacgattaagctgtatggtaataagggggataaggtaattgagactaatatgatacgctgttggctgagtatggtatgttgttagctaaaggtgatggcgtgggtacatgggaatatgctgttgctgcgtatactgtgtttggtggcattgtgggcaattgtggctcatctttgaactgaaggagtgttattcttttttcttttcctttctcgatctctgttatttctaatgatattaattgttcttggtttgataatatttcggttgttgttttaagttctactgtgctgatttcctcatctttgccatgtttcgctcatgtattttatcgataatcccgttttctaggtgtgcctttccgtagccctaccatttgaaaagctttttccatgtctcgcttattatctatgtttttgtgctaattcttaatagtatttctagcggttctggtattagtggtgggccttgatatactgcttggcttagactaatttctcctctgactgcattgctgtcctatgttgttactgtcactatgcatgtgaatcttttgactgtttatgcatagattaaagtttcctgtcgcgaagatgacgtgccggaaaattcatgaatcagatagagctactctactctgtttattgtgtttttcatgcatgtcctctactgattatcagatatttcccattccatcacccaggattatcagttggcagtacattgtcaatgatcatgatccactttaagttggctgatattagaaatgaacaaagaaaatttggttttaattttttttcaaagtgctgcaaacggttagactcaacctaatattctcgctcatttttctccgttggggcttctttgatgttattttggtctttccctgtatctgagattactaaggttttggccatgttgtcctggctctgtttcctattgttcttatggtttctcttgttccctacagttatctcattggttacattaatcttgtgtttcctgtagcgaccaagttgattatactgtaagtgtgttttctaccagcattgtatcctcttttgttattagtttatcgtcgctctccgtaggagaattatcggtatttgttttgaggtataaacctcgtgaggggctatgagcggttagagctcaactcgattttttttgttttcaatgtgtaaccttaccaacagcgacgcactaaatcatcatgcttgtaagacaagtggagattattggttcaaccttctgtgttagcatcttggaggttaattattttccaccgaagaaagctatgcctacccccttatcaaaaacatggttttaaacgttccctctttgcaagctaggactcataacttgactaaaaccggatttatctttataaaattcacacactggtcagaaactactcaagtaatgaaatttgagaattgaagacttcaaaaagaactaaaactttgtttaaatgcacctaaactgaagtaaacttgattacctgaagttctgcgccaccttgtaccactcgctctgcttctatcttttacaatctctcactgggggttgtttcttcagcagttgtaacacagctcattttcttagaagaaaaacttactctcatatagccggctaccgtcctgcatttcttggattggtggttccttttttacgaaataaggaagctctggtgcgtcttgggcggtgaatcctgataaatgtttgatgagtgggatctgtttcgttcttgagtgtaatgccctctcacatgcagtctcgtcttttcgttcaaaggtgtgttctctgtctgtctctctctctctgtctctctctctctctctctctctctgttttgctttcttggttgttattactgaaaaacacttgatgatctgtctttttatagattcctgtcatgtctttctctatgcgttctctctctatccctgtttttctgctctcttttctggttatccgtcaaattctggtatcagtgatgttttattaaaaattatcgctcttcccactttcttttgatcatatgtcccaattcttctagtttctgatttgcctctgctagtctgccttgtgtgtcttcagtgttttctatagttgcgagtacgtcatctgttactcttcctcctaattcttcacacgctgtttaggttttttgttcgtggttttctcggattttctttagattacctaggttttctctcttagctaccgtaattaaatttgattttccccgatcagtcactggtttctctagtttactctagttaccctttgctttgtttttgtgattactttttttattttcttaaattaaactaacatagtatattctctttggtcgatgcaccactttgacacttcgcgagcggttgaatattttatcaatttgttctcgatggtattaggtcggtacataagtcctgacggattttgctggtcgaacttgttgaatgtatttcactgttcaaacttaagagagacggctaagtttctgtcccgtgaaaattcaaagcttcctcttctctttcatttttagaaatttcatcatatcattttatttgtactcctatatggcgtggtattcattgaaacgagttttagtttagtggttattaacaacggacaaacactattttaaaacagttagcgatgccagtttcaatcgactagaaatagggagttatccatggatttccatccccctgcaaaaagaatgatgtcttcaaagttgagtcaattgtaaaatgagatcgaaagagattcatttcaaattttgtgttaaggatttcgggtatctggtgtatgcatttgagtaaaaaaccaaaaaaaaggattgaaaaaattcaccagagacaaaaactcaaaagatgtgactcaccctttttgtgactccacacaattccgaaacatgggagaccaccaggttgaatcttttcttttcttccgatacaggaatcagtggttcaaataatttttggtggtaataaacgagtaatgaatggttgttatgagcaaggttcttagtttgccgcttcaccctattagccgtttgatgtcattattcatgaaattcatgatgtggtcagtgtgacgcttttccagaaaagtggtgtactaggagaacaaagatgatagtatcacaatcatgcccatagtctatatttctcagaacagagacaaatgctcctgtattcatctccattcttgagaaatgtctaaattttatttggagtttgttggtattcagcatactctcgaagaaaggatcgacataaaactcactggatgtgatttgacaatgactttaaatttatcatgcagatcattgttagaagagttgattaacaagtttgcaattaccagtcgaaagttcaagatggaaaaaattcaactcagaccgtatattaaaaaagtagtgacttcgtatttcgagtcatctctataaaagttctaaaggcagagacaaatgtatgatgggggtgcattaagagacacttgtggatgaacatggacaatatttgcaatttagctattagctctgtggcaagttacacgtgaaccttgtcgacccaaacatccgtcaggacttatgggccaacctaataatatattaataatataggctaggttcaaatatttctcattttggagtgaaagaaataagaaacagaatgagtagatggttcagcaaaccttgaattttcgtgattaatacagaaggagccagctttgtggtgtcaggctgaaaaaaagtaaagtcagcttacttacttcactcaaaaatatgaaattttgaaattagtcccttattatatgtcagaaatagaacggttttaatagttcacgattacatgacaagataaagtttggaatgggaatcaggctattttctttcttcttcctcctgtcggtattctgattgctcctcccatctttagactggttcaattaatttgatcctcatcttctgctcataccatattttttcttgatatcctccggaagcgtttcttcctttttttgaccctcctctttctccgtggtcattctgctatctctaagcttcatctccctgctaatttatttccgtcatgatatcctccgatgttcctctttctctgatcctttttcttatctgctccgccgtagtcaagacaggaattacccggagaatcatttttttttccactaaaaaccttctgaattcatccattttcaacttaaaaagacatttttcgaatgagtcattcagtttttaaaaagcgtcaaatccttgagaacctacaataatttcatttactgttcccgagagtcttttttttgtggaaaaacgtcattttaatcccattcttttcctaagaattgtcgaaaactactcttaagtctcgaaatttacaatttccatttgatttccctctcaaaaaatagttttcaaatttaaaaatgcagtttctgacaaaaacttgcttctttcgaaatttcaatttaacaaaccaatttttatattatttttcttctgaaaactctcttaaaacatataatttcacaacattcatgctctttttctttctcaaaatcggatgaatcccataaagaaccgtcaaatactctcttaaccctctaatttcacggtcgttgatcgtttgataactgtaaacctctccaagtcctattttcaaccaacaatctcattttttcagTCAGAAGATCTCGATGAGAAATTCGAAGTCGAAGTGGATACATGTCCACATTTGCCCACAAACTTTGAAACAATGAAATGGCAGGAGCGGCGAAGATCCTTGGAATCCTTCCTCCAAACTTTAACTGGAAAGAAATGTATCTTGTCTAACATTTCTTACGATGCATCGATTAAAAAACTGCAAAAGATAATTCGGAACGACGCCAACATATTTTGTCAAGTGTTGGCAATCAGAAGCGTTACATGGGTTGCCACGGAACTCGGCGCGGAATTCTCCAAATTCTCTGTTTCTCTTCTACCAGATCTGTTGGAAAAGATGAAAGAGAAAAAACAGATATTGAGAAAGCCCCTTATTCGATGTACTTTGGAGGTTGGAAAGACGTTGCCGTTGGAAGATGGAATTCCAGTGATTTTATCGGCTCTCTCCATGGCAAATCCGGAGATAAAGAAGCAAACGATGTTATTTGTGGTTCAACAGCTTGAGGCGATGCAATTGGAGAGATTAAAAGGATTCATTCCATCATTACTTCCAGTTCTTGTTGAGCTTACAAAAAATGCTACTCAAGATGTCCGTGAAGTCGCTGTCCACGCCCTCGAATCAATTTACTGGAAGATAGGCCGATGTCGAATGCAATCTTTGTTATCAAAGTGTAGACATCTATTACCCAAAAGACACCCTTGGCATTGGATTAAAAAAGGATCTGACGAAAAAAAGAAAACAGAAAAACCGCCTTCTTCTTTTTTTCATGAAGCTATTGGCTCCATCATTTACTCCGGAGAAGAACTTCATGCCGCCAGATTCAGCAGATCCAACTGTATCCGGGTCGATCTGGAGGATCAGTTTATCCGACCGAATCTTCTTGTGGGACTTTTGAAGATCCTGGAAACTCATGCAGCCGACACTACATCTTTGAATCTGTACGAGTGCAGTGATGTTTTGGTGAGAGATGATACGGAGAAAAACGGAGCAAGAGAAGAGCGACGTTTGGCTGCTGCCATTCTCAATGAATCGGCCGATTTCGCTGCAATAGTGAACTGCCTCATCAAGCTGTTGGATGCGTTAAAACTGAGTCCGACGAAAGACGGCGATAATTTTCATTTGGAAATATCGGATAgtgagttcttgttcgaaacctcaaaacccttggaatgatcattgaatttggggatttttacctagtttccattcgagttttctttatactccctaaaaacatctgattccatgaatgctctcatgaatattatgaatcctcagaatatcaatgaagatcttttagaagcctagaatctcattcctctagaatcttttttccatcttaaattccaagaaccctttaatgaattctcttagatgctactttgaattcctctagaatccctgaaattccatgaacctccatcctgaatcttctgaatctccttttctctctttcagATCAAACCTTCTCGCCTGGACACTACGCGCGAATGGTTGGCCCGAACGACGTATTTCTCGGCCATGTTGGAGTCGTTCATCCGGAGGTACTCAGGAAATTCAACCTCATGTTGCCGGTTGCCGCCTTCGAGATCAAAATCTTTACGGAGGACGACTAA

>PZL-1

MWSSTTSSHQNQQKIHEKVYKNEHQVCLPPCELKDEEYEPKKSFYEMHQQTKRWTNWESQRIVTAPGHCATVDSVLLFEKNDNKFCLSESRDRVIRFWDVDNVERGVDSVANPWTVAQDDMAQLEWSWNMARDSNTDRFYSTSWDSTVKSWAITDNGAIQNLNTVNVGSAVQAVSCSGNENEIVCTTFAKRTAVIDSRTFGIVAEHSLHKRAVIGLAVKENKIFTSGEDRLMMMVDRRNMSRPVLFEYCQDAYKSSLSLQRNQLLTSTSDGKVKLYDATNFHDFQTYSVGAFTRQSLLQHGAHFVMARCKRGYYKFLMNSPGIRSLKRCSSHQLTAKPAKFDYSSEMKTLTIGNSMDRSERRSMISEDLDEKFEVEVDTCPHLPTNFETMKWQERRRSLESFLQTLTGKKCILSNISYDASIKKLQKIIRNDANIFCQVLAIRSVTWVATELGAEFSKFSVSLLPDLLEKMKEKKQILRKPLIRCTLEVGKTLPLEDGIPVILSALSMANPEIKKQTMLFVVQQLEAMQLERLKGFIPSLLPVLVELTKNATQDVREVAVHALESIYWKIGRCRMQSLLSKCRHLLPKRHPWHWIKKGSDEKKKTEKPPSSFFHEAIGSIIYSGEELHAARFSRSNCIRVDLEDQFIRPNLLVGLLKILETHAADTTSLNLYECSDVLVRDDTEKNGAREERRLAAAILNESADFAAIVNCLIKLLDALKLSPTKDGDNFHLEISDNQTFSPGHYARMVGPNDVFLGHVGVVHPEVLRKFNLMLPVAAFEIKIFTEDD

>kss-2

ATGGAATTCTCTCTTATTCGACTGCCAGATGTTCCCAAAATTCAAATTATTCAATTGATGACGATTGGCGAGAGgtgagtatctcgaggttttctttttgttatattttgttttacagAATTAAACTGGTCATCTCTTCTCGAAGAATGGAGAACTATTTGTCCAGGGTATTCAAGAAGCCCAATACAAATTCCGATTATAATATGAACTTGAAAGGCAAATTTTCTTTCATTTCTATCGGAGACGACGAGTGGAGAGTGTTTCTGGGTCCTTCGAAGATGGAGTATAGTAAGCCGGATAATATCTCAGAGGAGGATGTGAAACCATGgtgagttcccagaaatctcgagtaatgtactcaaagaaaacggagattcagGATAAACGAAAAGTGTACAGTGATCGAAAATACTCTTAACGTGTTCGTACGTCTCCAAAACGTGTTCCCTTGTGGGACTACCAACTTGTGCGTAGATTTGAACAAAATCGATCCGATGCCGATTCCGAATATACTGAGTTATCCCTGTGTCGCAAATTTGACAGGTATTCATTTCTACGGAGGAACAGTACAGAAATACGAGTTGGATGCAATTATGGAGTGGAGAAAAGAGAATACGATTCAGTTCATTACAATTTCAGATAACAAAATTCCATTGGACTATAGACACCCAAACgtgagtcaactttcaaacatttatatcttaattgccagtggagacattgggaaatggtcaacttcgaagttgctcatcgtaaaatttagttgatcaaaataaacagaacgacgctgttatttttaatcacgttgtttcacctcttctgtaagagctggcaatatcggaacccttaggacacgcccactattatgccaatctcgaaaaatggcagttctggccgaagatccgacaagttcttccacgtgaggtcgggaagcgaaaataaaattggtacagtaagaagttcaaaccataacccatgcgcctttaaggttttcaatagagagcgttaaaggcgcacgacaaaaagtaaaaagttcttactgtaccaattttattttcgctccccgacctcacgtggaagaactgtcggatcttcggccagaactgccatttttcgagattggcataatattggagaggaggaggaaggagaggggagaattgtggtagttgaggctgcgcggagaagagggagatggtgacccttctatactcctttatattttatacttcttatgataataaataatgatagagtggaaccaagacttagtttgaactcaagtatcaagggtctgaagtataataggtactcgcctattacacttccaatatttacaaaaaacaactcggcatcgataaagcgttgaccactttctgatatctacactgcagaccaagatgcatcgatattcagGCTCTTCGCTTCTCCGGGGTGATGTACAACGACGCTCGATGGGTACGAATCGAAGACTTACTATCAATGAGAAACGTGATCCGATCGTTTTTCGTAGACAATAATTTCAGTATGCAAGATCTGAACACATTTATCAAATATTGGACCAACTGCGACGAAAATATGTTTACATATCTCTATATTGGATACAAAAGTTTAGATATGTTAGAATTGATGAAGGATATCACTGTTCTCAAGGGTTTCCGATCCAAAAAATCATTTTATCTCATgttagtccttttattttataacgagaaacctcgagatttcaatgataaataagccataaattccaaaaaaaaaattttccagAGCATCGAATACAACTGAGAACCAACTTTTGATGACGGTTAGATTGGTATCTTATCCTGCCGAACCCGGACGACGAGAGCTCGAGTTCTACGTTCTGTCTGCCAACACACCTCATCATGTACAGGGTGGCAGCACGGAACCCGCGTGGACTCGAGAGTATCGCATTTTGCGACGATTGGCGAGAAAACGTTGGCTGGATGAGAAGAGGGTTACGGTAGAAGAGGCAAAGGAACTGGATGATTTGGAGAAGGAGTTGGCTGCTGAGGGAGTTCGGATGATTGATGGAATAATGACTGTAGAACAGAAACTGTAA

>KSS-2

MEFSLIRLPDVPKIQIIQLMTIGERIKLVISSRRMENYLSRVFKKPNTNSDYNMNLKGKFSFISIGDDEWRVFLGPSKMEYSKPDNISEEDVKPWINEKCTVIENTLNVFVRLQNVFPCGTTNLCVDLNKIDPMPIPNILSYPCVANLTGIHFYGGTVQKYELDAIMEWRKENTIQFITISDNKIPLDYRHPNALRFSGVMYNDARWVRIEDLLSMRNVIRSFFVDNNFSMQDLNTFIKYWTNCDENMFTYLYIGYKSLDMLELMKDITVLKGFRSKKSFYLIASNTTENQLLMTVRLVSYPAEPGRRELEFYVLSANTPHHVQGGSTEPAWTREYRILRRLARKRWLDEKRVTVEEAKELDDLEKELAAEGVRMIDGIMTVEQKL

>EG-kss-2

ATGGAATTCTCTCTTATTCGACTGCCAGATGTTCCCAAAATTCAAATTATTCAATTGATGACGATTGGCGAGAGgtgagtatctcgaggttttctttttgttatattttgttttacagAATTAAACTGGTCATCTCTTCTCGAAGAATGGAGAACTATTTGTCCAGGGTATTCAAGAAGCCCAATACAAATTCCGATTATAATATGAACTTGAAAGGCAAATTTTCTTTCATTTCTATCGGAGACGACGAGTGGAGAGTGTTTATGGGTCCTTCGAAGATGGAGTATAGTAAGCCGGATAATATCTCAGAGGAGGATGTGAAACCATGgtgagttcccagaaatctcgagtaatgtactcaaagaaaacggagattcagGATAAACGAAAAGTGTACAGTGATCGAAAATACTCTTAACGTGTTCGTACGTCTCCAAAACGTGTTCCCTTGTGGGACTACCAACTTGTGCGTAGATTTGAACAAAATCGATCCGATGCCGATTCCGAATATACTGAGTTATCCCTGTGTCGCAAATTTGACAGGTATTCATTTCTACGGAGGAACAGTACAGAAATACGAGTTGGATGCAATTATGGAGTGGAGAAAAGAGAATACGATTCAGTTCATTACAATTTCAGATAACAAAATTCCATTGGACTATAGACACCCAAACgtgagtcaactttcaaacatttatatcttaattgccagtggagacattgggaaatggtcaacttcgaagttgctcatcgtaaaatttagttgatcaaaataaacagaacgacgctgttatttttaatcacgttgtttcacctcttctgtaagagctggcaatatcggaacccttaggacacgcccactattatgccaatctcgaaaaatggcagttctggccgaagatccgacaagttcttccacgtgaggtcgggaagcgaaaataaaattggtacagtaagaagttcaaaccataacccatgcgcctttaaggttttcaatagagagcgttaaaggcgcacgacaaaaagtaaaaagttcttactgtaccaattttattttcgctccccgacctcacgtggaagaactgtcggatcttcggccagaactgccatttttcgagattggcataatattggagaggaggaggaaggagaggggagaattgtggtagttgaggctgcgcggagaagagggagatggtgacccttctatactcctttatattttatacttcttatgataataaataatgatagagtggaaccaagacttagtttgaactcaagtattaagggtctgaagtataataggtactcgcctattacacttccaatatttacaaaaaacaactcggcatcgataaagcgttgaccactttctgatatctacactgcagaccaagatgcatcgatattcagGCTCTTCGCTTCTCCGGGGTGATGTACAACGACGCTCGATGGGTACGAATCGAAGACTTACTATCAATGAGAAACGTGATCCGATCGTTTTTCGTAGACAATAATTTCAGTATGCAAGATCTGAACACATTTATCAAATATTGGACCAACTGCGACGAAAATATGTTTACATATCTCTATATTGGATACAAAAGTTTAGATATGTTAGAATTGATGAAGGATATCACTGTTCTCAAGGGTTTCCGATCCAAAAAATCATTTTATCTCATgttagtccttttattttataacgagaaacctcgagatttcaatgataaataagccataaattccaaaaaaaaaattttccagAGCATCGAATACAACTGAGAACCAACTTTTGATGACGGTTGGATTGGTATCTTATCCTGCCGAGCCAGGACGACGAGAGCTCGAGTTCTACGTTCTGTCTGCCAACACACCTCATCATGTACAGGGTGGCAGCACGGAACCCGCGTGGACTCGAGAGTATCGCATTTTGCGACGATTGGCGAGAAAACGTTGGCTGGATGAGAAGAGGGTTACGGTAGAAGAGGCAAAGGAACTGGATGATTTGGAGAAGGAGTTGGCTGCTGAGGGAGTTCGGATGATTGATGGAATAATGACTGTAGAACAGAAACTGTAA

>EG-KSS-2

MEFSLIRLPDVPKIQIIQLMTIGERIKLVISSRRMENYLSRVFKKPNTNSDYNMNLKGKFSFISIGDDEWRVFMGPSKMEYSKPDNISEEDVKPWINEKCTVIENTLNVFVRLQNVFPCGTTNLCVDLNKIDPMPIPNILSYPCVANLTGIHFYGGTVQKYELDAIMEWRKENTIQFITISDNKIPLDYRHPNALRFSGVMYNDARWVRIEDLLSMRNVIRSFFVDNNFSMQDLNTFIKYWTNCDENMFTYLYIGYKSLDMLELMKDITVLKGFRSKKSFYLIASNTTENQLLMTVGLVSYPAEPGRRELEFYVLSANTPHHVQGGSTEPAWTREYRILRRLARKRWLDEKRVTVEEAKELDDLEKELAAEGVRMIDGIMTVEQKL

>hyde-1

ATGAGTAACGCTCCCAGCTCGACAGACGCACCGTTAGCGgtatgttcatattaagctggtttttagtgaaaactggccacttttcacacattttaataaagaaaaaaaaacagtttttcacataaaatgctattttcaaataaaaataacatttctcagtcgaaatacgctgtttgcttctgaaaagttgttctaatgaaaaggttactaatttgaattcaataattttttccagcgaattttaacagaaaaaagctattttgctacaaaaagcgtgatttcctacaatttttcaccttaaaacccaaactgcgcacttttttaagctagtttgcatttataaaacgcctaatcctataataaacaataattttttgcagGAGAACGACTACGACGACCTTTTTGCCTACGGTCAGCGACCGCCAACACCTCAACTTCCACGCCCAgtacgtttatttcttttttaaccgaataatattttcaattatgattttcagGCTACCCCAATCGACTCTGGCCCCGAGTTGGACGAGATTgtaggttttttaatccggccgacgctgttttcgctgttttcactgcagaaaacgcgttccgtgcctgagaaatcgctgtttctagcgaatttcgaaagaaacccgctattttctagcggaattatcatttattttattatttaaatttttaatttttaatttaaatttttcatgacggaaaccataatttttcattcttttcgaccgtaaaacctctttttttcgaggggtttaactggaaaaacgctatttttaactcaaaaaatttttgtatcttcgaattccgaacagaaacgtcgtttttgggggaaaaacgaatttttcaaatttaaaatagctattttttacctaaagtcctggttttacagctgacaatgctattttcgattattttcgatctaaaaccctctttttacgggtaaaaacgcttctttcggttattttttcatctaaaaaccctcatttcaagcttggtaccggaaaacgtcatttttagacaaaaaacggttatttttactgttttgaacctagaaccctcttaaaacactcaaaaaaaatctatttttgactattttcgaccttgacccgtctttttacagctgaaaagtgctactttcgggtgttttttacctaaaaacccttcagttcaagcgatgtaccggaaaagttcatttttagaaaaaaaacgttatttttactgttttgaacctaaaaccctcttaaaacactcaaaaaaaacctatttttgactattttcgaccttgaaccgtctttttacagctgaaaagtgctactttcgggtgtttttcacctaaaaaccccgtagttcaagcgatgtaccgaaaaacttcatttttagagaaaaaccgttatttttactgttttgaacctaaaaccctcttaaaacactcaaaaaaaaacctatttttgactattttcgaccttgacccgtctttttacagctgaaaagtgctactttcgggtgttttttacctaaaaacccttcagttcaagcgatgtaccggaaaacttcatttttagaaaaaaaacgttatttttactgttttgaacctaaaaccctcttaaaacactcaaaaaaaacctatttttgactattttcgaccttgaaccgtctttttacagctgaaaagtgctactttcgggtgtttttcacctaaaaaccccgtagttcaagcgatgtaccgaaaaacttcatttttagagaaaaaccgttatttttactattttgaacctaaaaccctctttaaaacacatctgaaacaaaaatatctgttttgacagctaaaaaccctatattttccttcagACATATGAATCAGCAGAGGAAGAAGAGGTGGAGGTTTATACGGATGTAAACGATCTTCATTGGTCAAGCGTTGAGGCGCTCTCCAGAGCTACCCGCATCTTCAACGAAGACAGTATATTGCAACGATACGAGTTCGGAGACGTAGTGACCCGACAGAAGCTGATTGTAAAGTCAAAGATCAATCTCAAAGTTATCGCCGTTGTTATTCGCGATGTTAGCTTCGACGAGGATAGCTACGCGTCGTTCATCGATCTCCAGGACAAACTCCATCAGAATATCTGTCGGAAACGAACGCTGGTCGCCATCGGAACCCATGACTTGGATTTAGTTCAAGGTCCGTTCGAATTGCGGGCAGAAGCCCCAAGTAAGATTAAATTCTGTCTTCCGAATCAGACGAAGGAGTACACGGCGACTGAGATGATGCAGTACTTCAGCGGCTTCTCTAATAATTTCCGAGCCATGCCACAGCTCCCTGTTTTATATGATAAAACAGAACGGATCTGCTCCATGCCGCCTATTATCAATGGAGAACACTCGAAGATCACTATCAAGACCAAGAACGTGTTCATTGAGGCGACGGCGACGGATAAGCAGAAGGCGTGCGTGGTCCTTGACACCATCGTCACCCTCTTCTCCCAATACTGCCAAAAACCGTTCCATGTCGAACAGGTGGAGATAGAATACGAGGAGACTGGAGAAAAGGATATTTTTCCTTTTTTCAATCAAAAAAATAAAAATAAACATGGAAGACCAGTCTTTAAATACGAGGAAATGGCGATTCTCCTAAACAAGATGTCCCTGAAAGCGGAGATCGCAATGAAATGCGGGCACAAAGTAATTGTGACGTGGCCCACTCCTCCTGGAGCGCTCCGTGAAATTATCCAAGACGTCGACGTTGGGTACGGCTACGACAAACTGATGAAAAAGCGTCAGgtgcgggtttaaagattatttttccctatttgtagcacaaaaatgctggtttcaacttgaaaaccctatttaccataaaaaaaactcttttccatcttgaaaaacccctattttaatttggaaacccgtttttcggccttcaaagctcctttttcaagagaaaaacctcattttccactcgtaaaatcccatttttcaccccttggatccctttttctaaggagaaatcctctttttccactcaaaaggccccttttccaatagaaaaccccatttttccagtcttaggatccctttctcctagtggaaatcctcttttctacctctaaaatccccatttccacccttaggatccttttcttcccctagaatcctcttttctcctccccaaaatcctccttttcttccagTCGAACACTGTCGCCGTCGCCTTCTCAATCAACAAGCTCTGCGACAAATTCCGGATCAAAATCGCCACATTCGGATGGACAGAGGCCCTCAGCTTCGCCCTCTGCTCCCGAGACGACATCTCGACGAAGCTGCGCCTCCCGAAAAAAGTACTCTCGGAAGCCGTCCACATTGGAAATCCGAAGACACTGGAATTCCAGGTCGCTCGGACCTCACTTCTTCCGGGTCTTCTGAAGACGTTGGCTTCAAATCGTGATATGCCTCTTCCCCTGAAGCTCTTCGAGCTCCAGGATGTCATTCTGAAAGATGAAAAGATGGGAGCAAGAAACGAGAGACGACTCGCCGCTGTCTACTACAACAAAGCTGCCGGATTAGAGATTATCCAAGGATTTTTGGATCGAATGATGAGGATGCTGAATGTGAAGCCGATAGGTGATCCTCGAGGATATCAGACTAAAAAGTTTAATCgtgagtttttactttatgatcttcagggatttccctagatccttcctcccccccaagagatcccatggattcttggaatccctaaaaatatccccagaatttcatgggtcctcctccctttctcccatctccttaatccatcctctcttcttccagATGTGACATTCGAGGAAAAGAATTGCTACGAAGTCCATTCCCCCAGCGGCGAAATCCTCGGACGATACGGAGTGCTCCATTCAGAGGTCACCGCCGCTTTCGGACTCACCCTTCCATGTGAAGCAATGGAGATTAGCATCTCCTCATTTTTCCCAAACTAA

>HYDE-1

MSNAPSSTDAPLAENDYDDLFAYGQRPPTPQLPRPATPIDSGPELDEITYESAEEEEVEVYTDVNDLHWSSVEALSRATRIFNEDSILQRYEFGDVVTRQKLIVKSKINLKVIAVVIRDVSFDEDSYASFIDLQDKLHQNICRKRTLVAIGTHDLDLVQGPFELRAEAPSKIKFCLPNQTKEYTATEMMQYFSGFSNNFRAMPQLPVLYDKTERICSMPPIINGEHSKITIKTKNVFIEATATDKQKACVVLDTIVTLFSQYCQKPFHVEQVEIEYEETGEKDIFPFFNQKNKNKHGRPVFKYEEMAILLNKMSLKAEIAMKCGHKVIVTWPTPPGALREIIQDVDVGYGYDKLMKKRQSNTVAVAFSINKLCDKFRIKIATFGWTEALSFALCSRDDISTKLRLPKKVLSEAVHIGNPKTLEFQVARTSLLPGLLKTLASNRDMPLPLKLFELQDVILKDEKMGARNERRLAAVYYNKAAGLEIIQGFLDRMMRMLNVKPIGDPRGYQTKKFNHVTFEEKNCYEVHSPSGEILGRYGVLHSEVTAAFGLTLPCEAMEISISSFFPN

>kss-3.1

ATGGAATTCCCTCTTATCCAACTGCCAGATCTCCCCAAAATTCAAATTATTCAATTGATGTCTATTGGGGAGAAgtaagtattactagagagtttcccaagacactcatttatttccagAATCAAACTGGCTCTTTCTTCTCGAAAAATGGAGAACTATCTGGCGTGGGTGTTCAAAAATCATATGCTCGATGTGGATTGTTTCATTTTTTTGCGAGGCTACCAGTCTATTATCAATATAGATAACAACGAATGGAAACTGTGTTTGAACCGTTTAAAAAAGGGTTGTGAAAATCCAGGTTATATTAAAGTGGAAGAATTAGAACCGTGgtgagttctgagaagtttcaacaaaattcagtaaaagaatacataatttcagGATAAACGATGAGTGGACAATGGTAGAGAAAACTTTCAATGTGTTCATACGTCTTCAAAAAGTTCTTCCGTGCCAGTTTACCCATTTGTGCGTAATTCTGAATGTCGAACCGACGCCGATTATAGAAATTCTGAGTTATCCGTGCGTCGCGAACTTGACAGGTGTGCATTTCTATGGAGGAACAGTTCAGAGGAACGAGTTGGATGCAATTATGGAATGGAGAAAAGAGAACACGATGCAGTATATTTCGGTTACAGATAACGAAATTCCATTGGATTATAGACATCCGAATgtgagccttcttttgaatagttatatcctatcaaaacacaaaaagtgcagaatgtttcgctgagcaatttcttagttgaccgtattctaatatctccactcttaaccaaaatacataaatagattcagGCATTTCGTTTCACTGGAGTAGTGTATGGCGATGCTCGATGGATACGTCTCGAAGATTTACTCTCGATGAGAAACTTCTATCGACCGATTTTCAGTGTGAATAATTTCAGTATGAAGGATCTGAACACATTTATTAAATATTGGATCAACTGCGATGAGAATATGTTCACGTATCTAGTTATTGATTATAAAAATATCAATTTATCAGAATTAATGAAGGATATCACTGTTCTCAAGGGATTCCGATCAGAAAAACCATTTTATCTGATgttagttccttcttttttatattcagagaatgtttaagactaattaattgggctcccggacatttcgaaacacaacaacattcgagttttcaaatgtcctgtttcgaaaagtgaaggtacgaacgagatcaaaggggttcgagatgtccgggagccactaattgatcaaaagaaaaatggtttccagAGCATCGAATTCAACTGAAAACCAACTCCTATTGACAATTCAATTGATAGAACACACTGAATCTCGAAGACGAAATTCGCTTGACTTCTCGATTCAGTATGCCAATGAACCTCATAAATTCATAAACGACGACACACTACCTGCATGGACTCGAGAATACCGTATTCTGGGACGATTGGCGAGAAAACGTTGGTTGGAAGAGGAGATGATTACTGTAGGAATATCGATTGAAGAAATAAAAGAACTCGATGATTTGGAGGAGGAGTTGACAGTGGATGAAGTTCGGATCATTGATGGAATAATGACTGTTCAAGAGAAATTGTAA

>KSS-3.1

MEFPLIQLPDLPKIQIIQLMSIGEKIKLALSSRKMENYLAWVFKNHMLDVDCFIFLRGYQSIINIDNNEWKLCLNRLKKGCENPGYIKVEELEPWINDEWTMVEKTFNVFIRLQKVLPCQFTHLCVILNVEPTPIIEILSYPCVANLTGVHFYGGTVQRNELDAIMEWRKENTMQYISVTDNEIPLDYRHPNAFRFTGVVYGDARWIRLEDLLSMRNFYRPIFSVNNFSMKDLNTFIKYWINCDENMFTYLVIDYKNINLSELMKDITVLKGFRSEKPFYLIASNSTENQLLLTIQLIEHTESRRRNSLDFSIQYANEPHKFINDDTLPAWTREYRILGRLARKRWLEEEMITVGISIEEIKELDDLEEELTVDEVRIIDGIMTVQEKL

>kss-3.4

ATGGAATTCCCTCTTATCCAACTGCCAGATCTCCCCAAAATTCAAATTATTCAATTGATGTCTATTGGGGAGAAgtaagtattactagagagtttcccaagacactcatttatttccagAATCAAACTGGCTCTTTCTTCTCGAAAAATGGAGAACTATCTGGCGTGGGTGTTCAAAAATCATATGCTCGATGTGGATTGTTTTATTTTTTTGCGAGGATACCAGTCTATTATCAATATAGATAACAACGAATGGAAACTGTGTTTGAATCGTTTAAAAAAGGGTTGTGAAAATCCAGGCTATATTAAAGTGGAAGAATTAGAACCGTGgtgagttctgagaagtttcaacaaaattcagtaaaaaaatacggtgttttagGATAAACGATGAGTGGACAATGGTAGAGAAAACTTTCAATGTGTTCATACGTCTTCAAAAAGTTCTTCCGTGCCAGTTTACCCATTTGTGCGTAATTCTGAATGTCGAACCGACGCCGATTATAGAAATTCTGAGTTATCCGTGCGTCGCGAACTTGACAGGTGTGCATTTCTATGGAGGAACAGTTCAGAGGAACGAGTTGGATGCAATTATGGAATGGAGAAAAGAGAACACGATGCAGTATATTTCGGTTACAGATAACGAAATTCCATTGGATTATAGACATCCGAATgtgagccttcttttgaatagttatatcctatcaaaacacaaaaagtgcagaatgtttcgctgagcaatttcttagttgaccgtattctaatatctccactcttaaccaaaatacataaatagattcagGCATTTCGTTTCACTGGAGTAGTGTATGGCGATGCTCGATGGATAAGTCTCGAAGATTTACTCTCGATGAGAAACTTCTATCGACCGATTTTCAGTGTGAATAATTTCAGTATGAAGGATCTGAACACATTTATTAAATATTGGATCAACTGCGATGAGAATATGTTCACGTATCTAGTTATTGATTATAAAAATATCAATTTATCAGAATTAATGAAGGATATCACTGTTCTCAAGGGTTTCCGATCAGAAAAACCATTTTATCTGATgttagttcattcttttttattttcagagaatacttaataataattaatttttttttcaaacttttctgtctgaacctgaggtgccaaacatcctgaaaaggtccaataagagcgagatgctcatggtcatgttactaaatgttaacaaacaatgcaccgaacccaggatcctctaatcctactctgtaataattaattgatcaaaataagaaagaaatggtttccagAGCATCGAATTCAACTGAAAACCAACTCCTATTGACAATTCAATTGATAGAACACACTGAATCCCGAAGACGAAATTCGCTTGACTTCTCGATTCAGTATGCCAATGAACCTCATAAATTCATAAACGACGACACACTACCTGCATGGACTCGAGAATACCGTATTCTGGGACGATTGGCGAGAAAACGTTGGTTGGAAGAGGAGATGATTACTGTAGGAATATCGATTGAAGAAATAAAAGAACTCGATGATTTGGAGGAGGAGTTGACAGTGGATGAAGTTCGGATCATTGATGGAATAATGACTGTTCAAGAGAAATTGTAA

>KSS-3.4

MEFPLIQLPDLPKIQIIQLMSIGEKIKLALSSRKMENYLAWVFKNHMLDVDCFIFLRGYQSIINIDNNEWKLCLNRLKKGCENPGYIKVEELEPWINDEWTMVEKTFNVFIRLQKVLPCQFTHLCVILNVEPTPIIEILSYPCVANLTGVHFYGGTVQRNELDAIMEWRKENTMQYISVTDNEIPLDYRHPNAFRFTGVVYGDARWISLEDLLSMRNFYRPIFSVNNFSMKDLNTFIKYWINCDENMFTYLVIDYKNINLSELMKDITVLKGFRSEKPFYLIASNSTENQLLLTIQLIEHTESRRRNSLDFSIQYANEPHKFINDDTLPAWTREYRILGRLARKRWLEEEMITVGISIEEIKELDDLEEELTVDEVRIIDGIMTVQEKL

>NIC-kss-3.4

ATGGAATTTTCTTTTATTCGACTCCCAGATGTTCCAAAAATTCAAATTATCCAATTGATGACTATTGGGGAGAAgtaagtatctctagagaggttcccaagacactcatttattttcagAATCAAACTGGCCCTTTCCTCTCGAAAAATGGAGAACTATCTGGCGTGGGTGTTCAAAAATCAAGTGCTCAATGTGGATTGTTTCATTTTTTTGCGAGGCTACCAGTCTATTATCAATATAGATAACAACGAATGGAAACTGTGTTTGAACCGTTTAAAAAAGGGTTGTGAAAATCCAGGTTATATTAAAATGGAAGAATTGAAACCGTGgtgagcaatatcgagaaattcagtgaaagaatacggtgttttagGATAAACGATGAGTGGACAATGATAGAGAAAACTTTCAATGTGTTCATACGTCTCCAAAAGGTTCTTCCGTGCCACTTTACCCATTTGTGCGTAATTCTGAATGTCGAACCGACGACGATTCTAGAAATTCTGAGTTATCCGTGCGTCGCGAACTTAAAAGGTGTGCATTTCTATGGAGGAAAAGTGAAAAGAAACGAGTTGGATGCAATTATGGAATGGAGAAAAGAGAATACGATTCAGTATATTTCAGTTACAGATAACGAAATTCCATTGGATTATAGACATCCGAATgtgagccttttttcaaatagttacatcctatcagttaaaaaaatgtgcagaatgttccgctgagcaatttcttagttgaccgtattctattatcttcattcttaaccgaaatacataaatatattcagGCATTTCGTTTCACTGGAGTTGTGTATGGTGATGCTCGATGGATACGTCTCGAAGATTTACTCTCGATGAGAAACTTCTATCGACCGATTTTCGGAAAAAATAATTTCAGTATGAAGGATCTGAACACATTTATTAAATATTGGATCAACTGCGAAGAGAATATGTTCACGTATCTAGTTATTGATTACGAAAATATCAATTTATCAGAATTAATGAAGGATATCACTGTTCTCAAGGGTTTCCGATCAGAAAAACCATTTTATCTGATgttagtttcttcttttttatattcagataatacttattattaattaattgatcaaaagaaaaaagaaacaattttcagAGCATCGAATTCAACTGAAAACCAACTCTTATTGACAATTCAATTAATAGAACACACTGAATCCCGAAGACGAAATTCGCTTGACTTCTCGATTCAGTATGCTAACAAACCACATAAATTCATAAACGACGACACACTACCTGCATGGACTCGAGAATACCGTATTCTGGGGCGATTGGCGAGAAAACGTTGGTTAGAAGAGGAAATGATTACTGTAGGAATATCCATTGAAGAAATAAAAGAACTCGATGATTTGGAGGAGGAGTTGACTGTGGAGGAAGTTCGGATGATTAATGGAATAATGACTGTCAAAGAGAAATTGTAA

>NIC-KSS-3.4

MEFSFIRLPDVPKIQIIQLMTIGEKIKLALSSRKMENYLAWVFKNQVLNVDCFIFLRGYQSIINIDNNEWKLCLNRLKKGCENPGYIKMEELKPWINDEWTMIEKTFNVFIRLQKVLPCHFTHLCVILNVEPTTILEILSYPCVANLKGVHFYGGKVKRNELDAIMEWRKENTIQYISVTDNEIPLDYRHPNAFRFTGVVYGDARWIRLEDLLSMRNFYRPIFGKNNFSMKDLNTFIKYWINCEENMFTYLVIDYENINLSELMKDITVLKGFRSEKPFYLIASNSTENQLLLTIQLIEHTESRRRNSLDFSIQYANKPHKFINDDTLPAWTREYRILGRLARKRWLEEEMITVGISIEEIKELDDLEEELTVEEVRMINGIMTVKEKL

>EG-KSS-1

MEFPLIRLPDVPKIQIIQLMSIGERIKLALSSRKMENYLEWVFKNRALDAVYTIELRGDTSFIGIDTKTWVLCMNHFKPECKKPGYISVEELKPWIDEKWTMMERTFNVFIRLQKVIPCELTQLFVTLKTEPTPILEILSYPCVANLTGVHFFGGTVQRKELDAIMEWRKENTIQYISVTDNEIPLDYRHPNAFRFTGVVYGDARWISLEDLLSMRNFYRPIFGKNRFSMKDLNTFIKYWINCDENMFTYLVIDYKDINISELMKDITVLKSFRSEKPFYLIASNSTQILLTIQLIEHTEPQIRNVLDFSIQYANEPHKFRRGGTVPAWTREYRILGRLARKRWLEEEMITVGISIEEIKELDDLEEELTVEEVRMIDGIMTVQEKL

>EG-PZL-1

MWSSTTSSHQNQQKIHEKVYKNEHQVRVPPCELKREEYEPKKSFYEMHQQTKRWTNWESQRIVTAPGHCATVDSVLLFEKNDNKFCLSESRDRVIRFWDVDNVERGVDSVANPWTVAQDDMAQLEWSWNMARDSNTDRFYSTSWDSTVKSWAITDNGAIQNLNTVNVGSAVQAVSCSGNENEIVCTTFAKRTAVIDSRTFGIVAEHSLHKRAVIGLAVKENKIFTCGEDRLMMMVDRRNMSRPVLFEYCQDAYKSSLSLQRNQLLTSTSDGKVKLYDATNFHDFQTYSVGAFTRQSLLQHGAHFVMARCKRGYYKFLMNSPGIRSPKRCSSHQLTAKPAKFDYSSEMKTLTIGNSMDRSERRSMISEDLDEKFEVEVDTCPHLPTNFETMKWQERRRSLESFLQTLTGKKCILSNISYDASIKKLQKIIRNDANIFGQVLAIRSVKWVATELGAEFSKFSVSLLPDLLEKMKEKKQILRKPLIRCTLEVGKTLPLEDGIPVILSALSMANPEIKKQTMLFVVQQLEAMQMERLKGFIPSLLPVLVELTKNATQDVREVAVHALESIYWKIGRCRMQSLLSKCRHLLPKRHPWHWIKKGSDEKKKTEKPPSSFFHEAIGSIIYSGEELHAARFSRSNCIRVDLEDQFIRPNLLVGLLKILETHAADTTSLNLYECSDVLVRDDTEKNGAREERRLAAAILNESADFAAIVNCLIKLLDALKLSPTKDGDNFHLEISDNQTFSPGHYARMVGPNDVFLGHVGVVHPEVLRKFNLMLPVAAFEIKIFTEDD

>NIC-HYDE-1

MAERRIYENVVGDIGNLYDEIAHVRLPAAPIPTPRLNLPTVPLPANRYDLLSVEGLSRAIRIFKQEVESPEYRFSDTKTRQKIIVKRETAQVRPYVVGVILSDVCFDEDSYASFIRIQDKLHQNICRKRTLVAIGTHDLDTIQGPFEYRAEAPNKIKFRPLNQTKDYTAEELMTLYSKDKNMKDTIKTFKKKSLVPVIYDKNGVVCSMPPFISGAHSEITQKTKNVFIEATATDKQKAYVVLDTIVTLFSQYCQKPFHVEQVEVEYEDKNEKEYFPFTCSKKMKNNTPEIRTKIVLNFKDEEMAILLNKMSLKAEVASKGGLKVLVPTTRHDILQACDLGEDVGVAYGRKFFVTKLHESNSVAVAVTSPFNYLCDNLRIKLSVFGWTEALNFALCSRDDISTKLRLPDALSEAVHTLEIRRHWNSKSLGPLFFRVF
